# Individual-Specific Areal-Level Parcellations Improve Functional Connectivity Prediction of Behavior

**DOI:** 10.1101/2021.01.16.426943

**Authors:** Ru Kong, Qing Yang, Evan Gordon, Aihuiping Xue, Xiaoxuan Yan, Csaba Orban, Xi-Nian Zuo, Nathan Spreng, Tian Ge, Avram Holmes, Simon Eickhoff, B.T. Thomas Yeo

**Affiliations:** Department of Electrical and Computer Engineering, National University of Singapore, Singapore; Centre for Sleep and Cognition (CSC) & Centre for Translational Magnetic Resonance Research (TMR), National University of Singapore, Singapore; N.1 Institute for Health and Institute for Digital Medicine (WisDM), National University of Singapore, Singapore; Department of Radiology, Washington University School of Medicine, St. Louis, MO, USA; State Key Laboratory of Cognitive Neuroscience and Learning/IDG McGovern Institute for Brain Research, Beijing Normal University, Beijing, China; National Basic Public Science Data Center, Chinese Academy of Sciences, Beijing, China; Laboratory of Brain and Cognition, Department of Neurology and Neurosurgery, Montreal; Neurological Institute, Departments of Psychiatry and Psychology, McGill University, Montreal, Canada; McConnell Brain Imaging Centre, Montreal, Canada; Psychiatric & Neurodevelopmental Genetics Unit, Center for Genomic Medicine, Massachusetts General Hospital, Boston, MA, USA; Martinos Center for Biomedical Imaging, Massachusetts General Hospital, Charlestown, MA, USA; Yale University, New Haven, CT, USA; Institute for Systems Neuroscience, Medical Faculty, Heinrich-Heine University Düsseldorf, Düsseldorf, Germany; Institute of Neuroscience and Medicine, Brain & Behaviour (INM-7), Research Center Jülich, Jülich, Germany; Integrative Sciences and Engineering Programme (ISEP), National University of Singapore, Singapore

## Abstract

Resting-state functional MRI (rs-fMRI) allows estimation of individual-specific cortical parcellations. We have previously developed a multi-session hierarchical Bayesian model (MS-HBM) for estimating high-quality individual-specific network-level parcellations. Here, we extend the model to estimate individual-specific areal-level parcellations. While network-level parcellations comprise spatially distributed networks spanning the cortex, the consensus is that areal-level parcels should be spatially localized, i.e., should not span multiple lobes. There is disagreement about whether areal-level parcels should be strictly contiguous or comprise multiple non-contiguous components, therefore we considered three areal-level MS-HBM variants spanning these range of possibilities. Individual-specific MS-HBM parcellations estimated using 10min of data generalized better than other approaches using 150min of data to out-of-sample rs-fMRI and task-fMRI from the same individuals. Resting-state functional connectivity (RSFC) derived from MS-HBM parcellations also achieved the best behavioral prediction performance. Among the three MS-HBM variants, the strictly contiguous MS-HBM (cMS-HBM) exhibited the best resting-state homogeneity and most uniform within-parcel task activation. In terms of behavioral prediction, the gradient-infused MS-HBM (gMS-HBM) was numerically the best, but differences among MS-HBM variants were not statistically significant. Overall, these results suggest that areal-level MS-HBMs can capture behaviorally meaningful individual-specific parcellation features beyond group-level parcellations. Multi-resolution trained models and parcellations are publicly available (https://github.com/ThomasYeoLab/CBIG/tree/master/stable_projects/brain_parcellation/Kong2022_ArealMSHBM).

## Introduction

The human cerebral cortex comprises hundreds of cortical areas with distinct function, architectonics, connectivity, and topography (Kaas, 1987; Felleman and Van Essen, 1991; Eickhoff et al., 2018a). These areas are thought to be organized into at least six to twenty spatially distributed large-scale networks that broadly subserve distinct aspects of human cognition (Goldman-Rakic, 1988; Mesulam, 1990; Smith et al., 2009; Bressler and Menon, 2010; Uddin et al., 2019). Accurate parcellation of the cerebral cortex into areas and networks is therefore an important problem in systems neuroscience. The advent of in-vivo non-invasive brain imaging techniques, such as functional magnetic resonance imaging (fMRI), has enabled the delineation of cortical parcels that approximate these cortical areas (Sereno et al., 1995; Van Essen and Glasser, 2014; Eickhoff et al., 2018b).

A widely used approach for estimating network-level and areal-level cortical parcellations is resting-state functional connectivity (RSFC). RSFC reflects the synchrony of fMRI signals between brain regions, while a participant is lying at rest without performing any explicit task, i.e., resting-state fMRI (rs-fMRI; Biswal et al., 1995; Fox and Raichle, 2007; Buckner et al., 2013). Most RSFC studies have focused on estimating group-level parcellations obtained by averaging data across many individuals (Power et al., 2011; Yeo et al., 2011; Craddock et al., 2012; Zuo et al., 2012; Gordon et al., 2016). These group-level parcellations have provided important insights into brain network organization, but fail to capture individual-specific parcellation features (Harrison et al., 2015; Laumann et al., 2015; Braga and Buckner, 2017; Gordon et al., 2017a). Furthermore, recent studies have shown that individual-specific parcellation topography is behaviorally relevant (Salehi et al., 2018; Bijsterbosch et al., 2019; Kong et al., 2019; Li et al., 2019b; Mwilambwe-Tshilobo et al., 2019; Seitzman et al., 2019; Cui et al., 2020), motivating significant interest in estimating individual-specific parcellations.

Most individual-specific parcellations only account for inter-subject (between-subject) variability, but not intra-subject (within-subject) variability. However, inter-subject and intra-subject RSFC variability can be markedly different across brain regions (Mueller et al., 2013; Chen et al., 2015; Laumann et al., 2015). For example, the sensory-motor cortex exhibits low inter-subject variability, but high intra-subject variability (Mueller et al., 2013; Laumann et al., 2015). Therefore, it is important to consider both inter-subject and intra-subject variability when estimating individual-specific parcellations (Mejia et al., 2015, 2018; Kong et al., 2019). We have previously proposed a multi-session hierarchical Bayesian model (MS-HBM) of individual-specific network-level parcellation that accounted for both inter-subject and intra-subject variability (Kong et al., 2019). We demonstrated that compared with several alternative approaches, individual-specific MS-HBM networks generalized better to new resting-fMRI and task-fMRI data from the same individuals (Kong et al., 2019).

In this study, we extend the network-level MS-HBM to estimate individual-specific areal-level parcellations. While network-level parcellations comprise spatially distributed networks spanning the cortex, the consensus is that areal-level parcels should be spatially localized (Kaas, 1987; Amunts and Zilles, 2015), i.e., an areal-level parcel should not span multiple cortical lobes. Consistent with invasive studies (Amunts and Zilles, 2015), most areal-level parcellation approaches estimate spatially contiguous parcels (Shen et al., 2013; Honnorat et al., 2015; Gordon et al., 2016; Chong et al., 2017). However, a few studies have suggested that individual-specific areal-level parcels can be topologically disconnected (Glasser et al., 2016; Li et al., 2019b). For example, according to Glasser and colleagues, area 55b might comprise two disconnected, but spatially close, components in some individuals (Glasser et al., 2016). Given the lack of consensus, we considered three different spatial localization priors. Across the three priors, the resulting parcels ranged from being strictly contiguous to being spatially localized with multiple non-contiguous components.

We compared MS-HBM areal-level parcellations with three other approaches (Laumann et al., 2015; Schaefer et al., 2018; Li et al., 2019b) in terms of their generalizability to out-of-sample rs-fMRI and task-fMRI from the same individuals. Furthermore, a vast body of literature has shown that RSFC derived from group-level parcellations can be used to predict human behavior (Hampson et al., 2006; Finn et al., 2015; Rosenberg et al., 2016; Li et al., 2019a). Therefore, we also investigated whether RSFC derived from individual-specific MS-HBM parcellations could improve behavioral prediction compared with two other parcellation approaches (Schaefer et al., 2018; Li et al., 2019b).

## Methods

### Overview

We proposed the spatially-constrained MS-HBM to estimate individual-specific areal-level parcellations. The model distinguished between inter-subject and intra-subject functional connectivity variability, while incorporating spatial contiguity constraints. Three different contiguity constraints were considered: distributed MS-HBM, contiguous MS-HBM and gradient-infused MS-HBM. The resulting MS-HBM parcels ranged from being strictly contiguous (contiguous MS-HBM) to being spatially localized with multiple topologically disconnected components (distributed MS-HBM). Subsequent analyses proceeded in four stages. First, we explored the pattern of inter-subject and intra-subject functional variability across the cortex. Second, we examined the intra-subject reproducibility and inter-subject similarity of MS-HBM parcellations on two different datasets. Third, the MS-HBM was compared with three other approaches using new rs-fMRI and task-fMRI data from the same participants. Finally, we investigated whether functional connectivity of individual-specific parcellations could improve behavioral prediction.

### Multi-session rs-fMRI datasets

The Human Connectome Project (HCP) S1200 release (Van Essen et al., 2012a; Smith et al., 2013) comprised structural MRI, rs-fMRI and task-fMRI of 1094 young adults. All imaging data were collected on a custom-made Siemens 3T Skyra scanner using a multiband sequence. Each participant went through two fMRI sessions on two consecutive days. Two rs-fMRI runs were collected in each session. Each fMRI run was acquired at 2 mm isotropic resolution with a TR of 0.72 s and lasted for 14 min and 33 s. The structural data consisted of one 0.7 mm isotropic scan for each participant.

The Midnight Scanning Club (MSC) multi-session dataset comprised structural MRI, rs-fMRI and task-fMRI from 10 young adults (Gordon et al., 2017b; Gratton et al., 2018). All imaging data were collected on a Siemens Trio 3T MRI scanner using a 12-channel Head Matrix Coil. Each participant was scanned for 10 sessions of resting-state fMRI data. One rs-fMRI run was collected in each session. Each fMRI run was acquired at 4mm isotropic resolution with a TR of 2.2 s and lasted for 30 min. The structural data was collected across two separate days and consisted of four 0.8 mm isotropic T1-weighted images and four 0.8 mm isotropic T2-weighted images.

It is worth noting some significant acquisition differences between the two datasets, including scanner type (e.g., Skyra versus Trio), acquisition sequence (e.g., multiband versus non-multiband), and scan time (e.g. day versus midnight). These differences allowed us to test the robustness of our parcellation approach.

### Preprocessing

Details of the HCP preprocessing can be found elsewhere (Van Essen et al., 2012a; Glasser et al., 2013; Smith et al., 2013; HCP S1200 manual). Of particular importance is that the rs-fMRI data has been projected to the fs_LR32k surface space (Van Essen et al., 2012b), smoothed by a Gaussian kernel with 2mm full width at half maximum (FWHM), denoised with ICA-FIX (Griffanti et al., 2014; Salimi-Khorshidi et al., 2014) and aligned with MSMAll (Robinson et al., 2014). To eliminate global and head-motion related artifacts (Burgess et al., 2016; Siegel et al., 2016), additional nuisance regression and censoring was performed (Kong et al., 2019; Li et al., 2019a). Nuisance regressors comprised the global signal and its temporal derivative. Runs with more than 50% censored frames were removed. Participants with all four runs remaining (N = 835) were considered.

In the case of the MSC dataset, we utilized the preprocessed rs-fMRI data of 9 subjects on fs_LR32k surface space. Preprocessing steps included slice time correction, motion correction, field distortion correction, motion censoring, nuisance regression and bandpass filtering (Gordon et al., 2017b). Nuisance regressors comprised whole brain, ventricular and white matter signals, as well as motion regressors derived from Volterra expansion (Friston et al., 1996). The surface data was smoothed by a Gaussian kernel with 6mm FWHM. One participant (MSC08) exhibited excessive head motion and self-reported sleep (Gordon et al., 2017b; Seitzman et al., 2019) and was thus excluded from subsequent analyses.

### Functional connectivity profiles

As explained in the previous section, the preprocessed rs-fMRI data from the HCP and MSC datasets have been projected onto fs_LR32K surface space, comprising 59412 cortical vertices. A binarized connectivity profile of each cortical vertex was then computed as was done in our previous study (Kong et al., 2019). More specifically, we considered 1483 regions of interest (ROIs) consisting of single vertices uniformly distributed across the fs_LR32K surface meshes (Kong et al., 2019). For each rs-fMRI run of each participant, the Pearson’s correlation between the fMRI time series at each spatial location (59412 vertices) and the 1483 ROIs were computed. Outlier volumes were ignored when computing the correlations. The 59412 × 1483 RSFC (correlation) matrices were then binarized by keeping the top 10% of the correlations to obtain the final functional connectivity profile (Kong et al., 2019).

We note that because fMRI is spatially smooth and exhibits long-range correlations, therefore considering only 1483 ROI vertices (instead of all 59412 vertices) would reduce computational and memory demands, without losing much information. To verify significant information has not been lost, the following analysis was performed. For each HCP participant, a 59412 × 59412 RSFC matrix was computed from the first rs-fMRI run. We then correlated every pair of rows of the RSFC matrix, yielding a 59412 × 59412 RSFC similarity matrix for each HCP participant. An entry in this RSFC similarity matrix indicates the similarity of the functional connectivity profiles of two cortical locations. The procedure was repeated but using the 59412 × 1483 RSFC matrices to compute the 59412 × 59412 RSFC similarity matrices. Finally, for each HCP participant, we correlated the RSFC similarity matrix (generated from 1483 vertices) and RSFC similarity matrix (generated from 59412 vertices). The resulting correlations were high with r = 0.9832 ± 0.0041 (mean ± std) across HCP participants, suggesting that very little information was lost by only considering 1483 vertices.

### Group-level parcellation

We have previously developed a set of high-quality population-average areal-level parcellations of the cerebral cortex (Schaefer et al., 2018), which we will refer to as “Schaefer2018”. Although the Schaefer2018 parcellations are available in different spatial resolutions, we will mostly focus on the 400-region parcellation in this paper (Figure 4A) given that previous work has suggested that there might be between 300 and 400 human cortical areas (Van Essen et al., 2012b). The 400-region Schaefer2018 parcellation will be used to initialize the areal-level MS-HBM for estimating individual-specific parcellations. The Schaefer2018 parcellation will also be used as a baseline in our experiments.

### Areal-level multi-session hierarchical Bayesian model (MS-HBM)

The areal-level MS-HBM (Figure 1A) is the same as the network-level MS-HBM (Kong et al., 2019) except for one crucial detail, i.e., spatial localization prior Φ (Figure 1A). Nevertheless, for completeness, we will briefly explain the other components of the MS-HBM, although further details can be found elsewhere (Kong et al., 2019).

**Figure 1.**
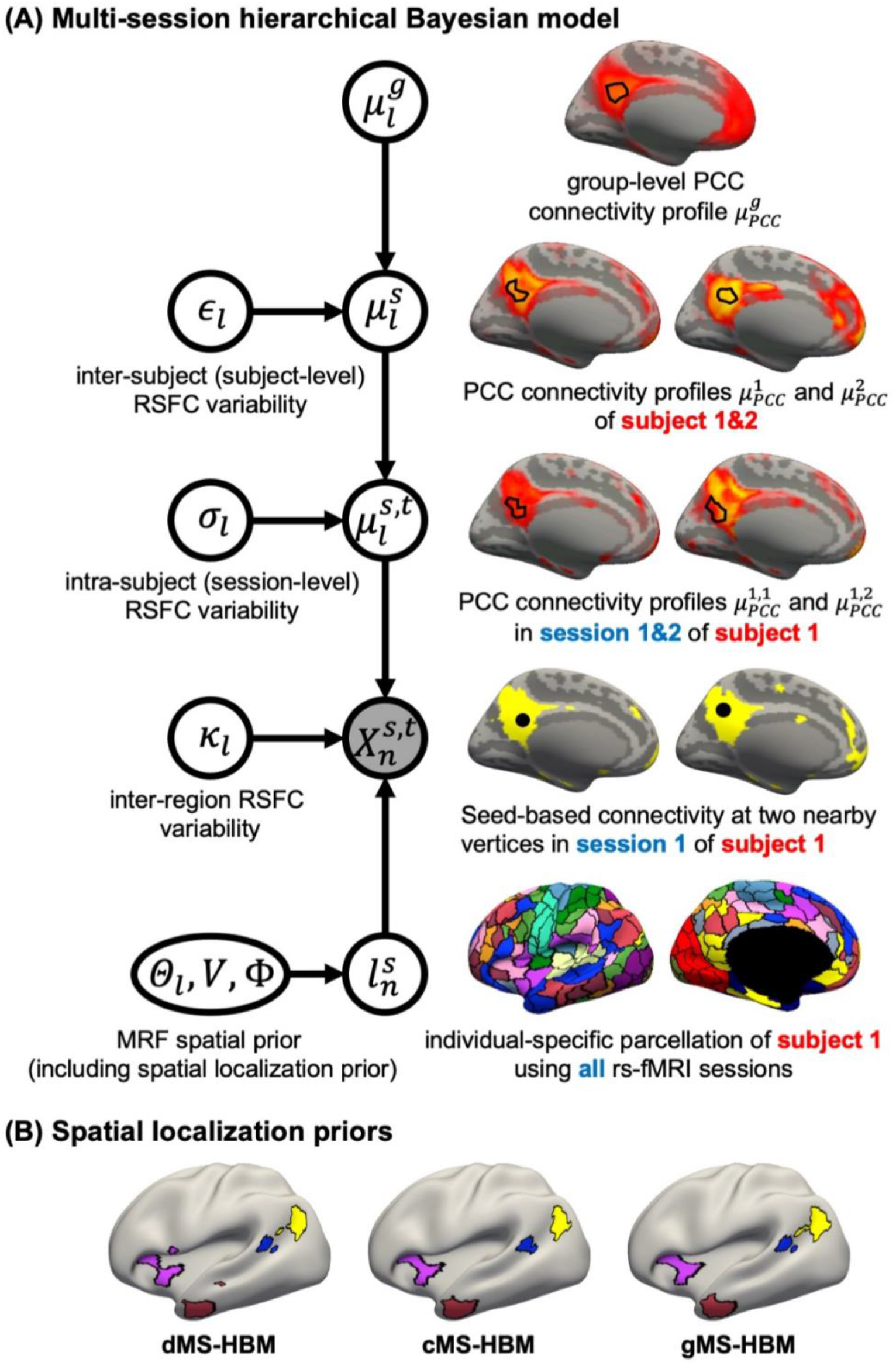
(A) Multi-session hierarchical Bayesian model (MS-HBM) of individual-specific areal-level parcellations. 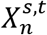 denote the RSFC profile at brain location *n* of subject *s* during rs-fMRI session *t*. The shaded circle indicates that 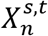 are the only observed variables. The goal is to estimate the parcel label 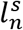 for subject s at location *n* given RSFC profiles from all sessions. 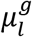 is the group-level RSFC profile of parcel *l*. 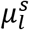 is the subject-specific RSFC profile of parcel *l*. A large *ϵ*_l_ indicates small inter-subject RSFC variability, i.e., the group-level and subject-specific RSFC profiles are very similar. 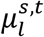 is the subject-specific RSFC profile of parcel *l* during session *t*. A large *σ*_l_ indicates small intra-subject RSFC variability, i.e., the subject-level and session-level RSFC profiles are very similar. *κ_l_* captures inter-region RSFC variability. A large *κ_l_* indicates small inter-region variability, i.e., two locations from the same parcel exhibit very similar RSFC profiles. Finally, *Θ_l_* captures inter-subject variability in the spatial distribution of parcels, smoothness prior *V* encourages parcel labels to be spatially smooth, and the spatial localization prior Φ ensures each parcel is spatially localized. The spatial localization prior Φ is the crucial difference from the original network-level MS-HBM (Kong et al., 2019). (B) Illustration of three different spatial localization priors. Individual-specific parcellations of the same HCP participant were estimated using distributed MS-HBM (dMS-HBM), contiguous MS-HBM (cMS-HBM), and gradient-infused MS-HBM (gMS-HBM). Four parcels depicted in pink, red, blue and yellow are shown here. All four parcels estimated by dMS-HBM were spatially close together but contained two separate components. All four parcels estimated by cMS-HBM were spatially contiguous. Three parcels (pink, red, yellow) estimated by gMS-HBM were spatially contiguous, while the blue parcel contained two separate components.

We denote the binarized functional connectivity profile of cortical vertex *n* during session *t* of subject *s* as 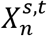. For example, the binarized functional connectivity profiles of a posterior cingulate cortex vertex 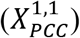 and a precuneus vertex 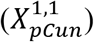 from the 1st session of the 1st subject are illustrated in Figure 1A (fourth row). The shaded circle indicates that 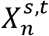 are the only observed variables. Based on the observed connectivity profiles of *all* vertices during *all* sessions of a single subject, the goal is to assign a parcel label 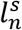 for each vertex *n* of subject s. Even though a vertex’s connectivity profiles are likely to be different across fMRI sessions, the vertex’s parcel label was assumed to be the same across sessions. For example, the individualspecific areal-level parcellation of the 1st subject using data from all available sessions is illustrated in Figure 1A (last row).

The multiple layers of the areal-level MS-HBM explicitly differentiate inter-subject (between-subject) functional connectivity variability from intra-subject (within-subject) functional connectivity variability (*ϵ_l_* and *σ_l_* in Figure 1A). The connectivity profiles of two vertices belong to the same parcel will not be identical. This variability is captured by *κ_l_* (Figure 1A). Some model parameters (e.g., group-level connectivity profiles) will be estimated from a training set comprising multi-session rs-fMRI data from multiple subjects. A new participant (possibly from another dataset) with single-session fMRI data could then be parcellated without access to the original training data.

The Markov random field (MRF) spatial prior (Figure 1A last row) is important because the observed functional connectivity profiles of individual subjects are generally very noisy. Therefore, additional priors were imposed on the parcellation. First, the spatial smoothness prior *V* encouraged neighboring vertices (e.g., PCC and pCun) to be assigned to the same parcels. Second, the inter-subject spatial variability prior *Θ_l,n_* denote the probability of parcel *l* occurring at a particular spatial location *n*. The two priors (*V* and *Θ_l,n_*) are also present in the network-level MS-HBM (Kong et al., 2019).

However, an additional spatial prior is necessary because of well-documented long-range connections spanning the cortex. Therefore, with the original MRF prior (Kong et al., 2019), brain locations with similar functional connectivity profiles could be grouped together regardless of spatial proximity. In the case of network-level MS-HBM, this is appropriate because large-scale networks are spatially distributed, e.g., the default network spans frontal, parietal, temporal and cingulate cortex. In the case of areal-level parcellations, there is the expectation that a single parcel should not span large spatial distances (Glasser et al., 2016; Gordon et al., 2016; Schaefer et al., 2018). Therefore, the areal-level MS-HBM incorporates an additional prior Φ constraining parcels to be spatially localized (Figure 1A last row).

As mentioned in the introduction, even though there is consensus that individual-specific areal-level parcels should be spatially localized, there are differing opinions about whether they should be spatially contiguous. Some studies have enforced spatially contiguous cortical parcels (Laumann et al., 2015; Gordon et al., 2016; Chong et al., 2017) consistent with invasive studies (Amunts and Zilles, 2015). Other studies have estimated parcels that might comprise multiple spatially close components (Glasser et al., 2016; Li et al., 2019b). For example, Glasser and colleagues suggested that area 55b might be split into two disconnected components in close spatial proximity. Given the lack of consensus, we consider three possible spatial localization priors (i.e, Φ in Figure 1A):

1. *Distributed MS-HBM (dMS-HBM)*. Previous studies have suggested that after registering cortical folding patterns, inter-individual variability in architectonic locations are different across architectonic areas (Fischl et al., 2008). One of the most spatially variable architectonic area is hOc5, which can be located in an adjacent sulcus away from the group-average location (Yeo et al., 2010a, 2010b). This variability corresponded to about 30mm. Therefore, similar to Glasser and colleagues (2016), Φ comprises a spatial localization prior constraining each individual-specific parcel to be within 30mm of the group-level Schaefer2018 parcel boundaries. We note that this prior only guarantees an individual-specific parcel to be spatially localized, but the parcel might comprise multiple distributed components (Figure 1B left panel). We refer to this prior as distributed MS-HBM (dMS-HBM).
2. *Contiguous MS-HBM(cMS-HBM)*. In addition to the 30mm prior from dMS-HBM, we include a spatial localization prior encouraging vertices comprising a parcel to not be too far from the parcel center, as was done in our previous study (Schaefer et al., 2018). If this spatial contiguity prior is sufficiently strong, then all individual-specific parcels will be spatially connected (Figure 1B middle panel). However, an overly strong prior will result overly round parcels, which is not biologically plausible (Vogt 2009). To ameliorate this issue, the estimation procedure starts with a very small weight on this spatial contiguity prior and then progressively increases the weights to ensure spatial contiguity. Thus, we refer to this prior as contiguous MS-HBM model (cMS-HBM). We note that requiring parcels to be spatially connected within an MRF framework is non-trivial; our approach is significantly less computationally expensive than competing approaches (Nowozin and Lampert, 2010; Honnorat et al., 2015).
3. *Gradient-infused MS-HBM (gMS-HBM)*. A well-known areal-level parcellation approach is the local gradient approach, which detects local abrupt changes (i.e., gradients) in RSFC across the cortex (Cohen et al., 2008). Our previous study (Schaefer et al., 2018) has suggested the utility of combining local gradient (Cohen et al., 2008; Gordon et al., 2016) and global clustering (Yeo et al., 2011) approaches for estimating areal-level parcellations. Therefore, we complemented the spatial contiguity prior in cMS-HBM with a prior based on local gradients in RSFC, which encouraged adjacent brain locations with gentle changes in functional connectivity to be grouped into the same parcel. In practice, we found that the gradient-infused prior, together with a very weak spatial contiguity prior, dramatically increased the number of spatially contiguous parcels (Figure 1B right panel). Furthermore, the parcels are also less round than cMS-HBM, which is in our opinion more biologically plausible. We refer to this prior as gradient-infused MS-HBM (gMS-HBM).

A more detailed mathematical explanation of the model can be found in Supplemental Methods S1. Given a dataset of subjects with multi-session rs-fMRI data, a variational Bayes expectation-maximization (VBEM) algorithm can be used to estimate the following model parameters (Kong et al., 2019): group-level parcel connectivity profiles 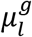, the inter-subject functional connectivity variability *ϵ_l_*, the intra-subject functional connectivity variability *σ_l_*, the spatial smoothness prior *V* and the inter-subject spatial variability prior *Θ_l_*. The individual-specific areal-level parcellation of a new participant could then be generated using these estimated group-level priors without access to the original training data. Furthermore, although the model requires multi-session fMRI data for parameter estimation, it can be applied to a single session fMRI data from a new participant (Kong et al., 2019). Details of the VBEM algorithm can be found in Supplementary Methods S2.

### Characterizing inter-subject and intra-subject functional connectivity variability

Previous studies have shown that sensory-motor regions exhibit lower inter-subject, but higher intra-subject functional connectivity variability than association regions (Mueller et al., 2013; Laumann et al., 2015; Kong et al., 2019). Therefore, we first evaluate whether estimates of areal-level inter-subject and intra-subject variability were consistent with previous work (Figure 2A). The HCP dataset was divided into training (N = 40), validation (N = 40) and test (N = 755) sets. Each HCP participant underwent two fMRI sessions on two consecutive days. Within each session, there were two rs-fMRI runs. All four runs were utilized.

**Figure 2.**
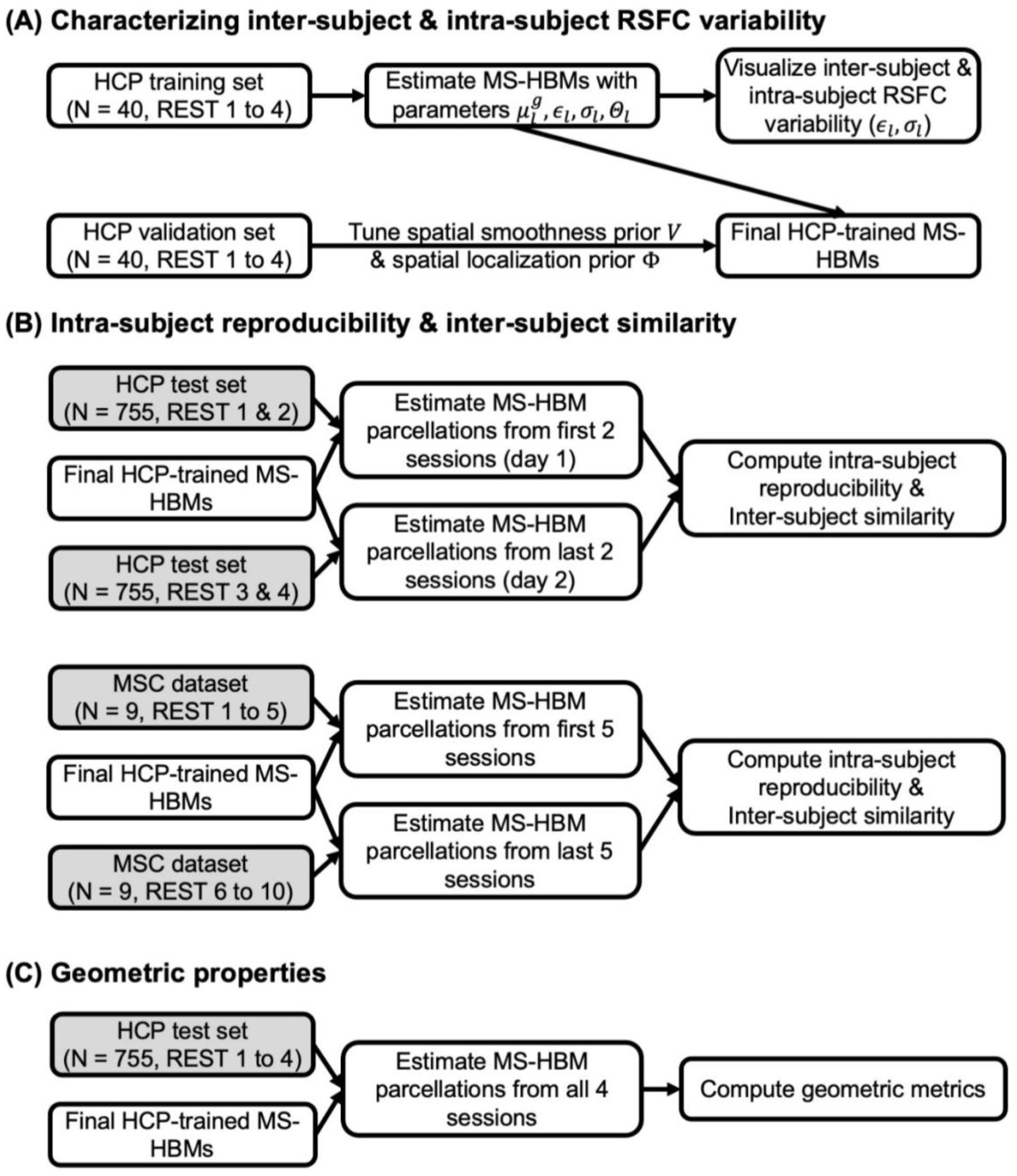
Flowcharts of analyses characterizing MS-HBMs. (A) Training MS-HBMs with HCP training and validation sets, as well as characterizing inter-subject and intra-subject RSFC variability. (B) Exploring intra-subject reproducibility and inter-subject similarity of MS-HBM parcellations using HCP test set and MSC dataset. (C) Characterizing geometric properties of MS-HBM parcellations using HCP test set. Shaded boxes (HCP test set and MSC dataset) were solely used for evaluation and not used at all for training or tuning the MS-HBM models.

The parameters of three MS-HBM variants (dMS-HBM, cMS-HBM and gMS-HBM) were estimated. More specifically, the group-level parcel connectivity profiles 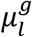, the inter-subject resting-state functional connectivity variability *ϵ_l_*, the intra-subject resting-state functional connectivity variability *σ_l_* and inter-subject spatial variability prior *Θ_l_* were estimated using the HCP training set (Figure 2A). We tuned the “free” parameters (associated with the spatial smoothness prior *V* and spatial localization prior Φ) using the HCP validation set (Figure 2A). The Schaefer2018 400-region group-level parcellation (Figure 4A) was used to initialize the optimization procedure. The final trained MS-HBMs (Figure 2A) were used in all subsequent analyses.

### Intra-subject reproducibility and inter-subject similarity of MS-HBM parcellations

Within-subject reliability is important for clinical applications (Shehzad et al., 2009; Birn et al., 2013; Zuo and Xing, 2014; Zuo et al., 2019). Having verified that the spatial patterns of inter-subject and intra-subject functional connectivity variability were consistent with previous work, we further characterized the intra-subject reproducibility and inter-subject similarity of individual-specific MS-HBM parcellations (Figure 2B). The three trained models (dMS-HBM, cMS-HBM and gMS-HBM) were applied to the HCP test set. Individual-specific MS-HBM parcellations were independently estimated using the first 2 runs (day 1) and the last 2 runs (day 2).

To evaluate the reproducibility of individual-specific parcellations, the Dice coefficient was computed for each parcel from the two parcellations of each participant:

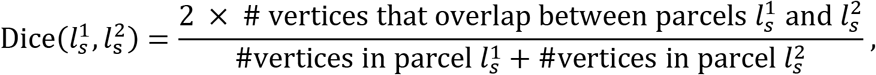

where 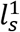 and 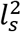 are parcel *l* from the two parcellations of subject s. The Dice coefficient is widely used for comparing parcellation or segmentation overlap (Destrieux et al., 2010; Sabuncu et al., 2010; Birn et al., 2013; Blumensath et al., 2013; Arslan et al., 2015; Honnorat et al., 2015; Salehi et al., 2018). The Dice coefficient is equal to 1 if there is perfect overlap between parcels and zero if there is no overlap between parcels. The Dice coefficients were averaged across all participants to provide insights into regional variation in intra-subject parcel similarity. Finally, the Dice coefficients were averaged across all parcels to provide an overall measure of intra-subject parcellation reproducibility.

To evaluate inter-subject parcellation similarity, for each pair of participants, the Dice coefficient was computed for each parcel. Since there were two parcellations for each participant, there were a total of four Dice coefficients for each parcel, which were then averaged. Furthermore, the Dice coefficients were averaged across all pairs of participants to provide insights into regional variation in inter-subject parcel similarity. Finally, the dice coefficients were averaged across all parcels to provide an overall measure of inter-subject parcellation similarity.

To evaluate whether the parameters of MS-HBM algorithms from one dataset could be generalized to another dataset with different acquisition protocols and preprocessing pipelines, we used the HCP model parameters to estimate individual-specific parcellations in the MSC dataset. More specifically, the MS-HBM parcellations were independently estimated using the first 5 sessions and the last 5 sessions for each MSC participant (Figure 2B).

### Geometric properties of MS-HBM parcellations

The three MS-HBM variants impose different spatial priors on areal-level parcellations. To characterize the geometric properties of the MS-HBM parcels (Figure 2C), the three trained models (dMS-HBM, cMS-HBM and gMS-HBM) were applied to the HCP test set using all four rs-fMRI runs. We then computed two metrics to characterize the geometry of the parcellations. First, for each parcellation, the number of spatially disjoint components was computed for each parcel and averaged across all parcels. Second, for each parcellation, a roundness metric was computed for each parcel and averaged across all parcels. Here, the roundness of a parcel is defined as 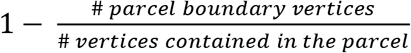; a larger value indicates that a parcel is rounder.

### Comparison with alternative approaches

Here, we compared the three MS-HBM approaches (dMS-HBM, cMS-HBM, and gMS-HBM) with three alternative approaches. The first approach was to apply the Schaefer2018 400-region group-level parcellation to individual subjects. The second approach is the well-known gradient-based boundary mapping algorithm that has been widely utilized to estimate individual-specific areal-level parcellation (Laumann et al., 2015; Gordon et al., 2017b). We will refer to this second approach as “Laumann2015” (https://sites.wustl.edu/petersenschlaggarlab/resources). The third approach is the recent individual-specific areal-level parcellation algorithm of Li and colleagues (Li et al., 2019b; http://nmr.mgh.harvard.edu/bid/DownLoad.html), which we will refer to as “Li2019”.

Evaluating the quality of individual-specific resting-state parcellations is difficult because of a lack of ground truth. Here, we considered two common evaluation metrics (Gordon et al., 2016; Chong et al., 2017; Schaefer et al., 2018; Kong et al., 2019): resting-state connectional homogeneity and task functional inhomogeneity (i.e., uniform task activation; see below). These metrics encode the principle that if an individual-specific parcellation captured the areal-level organization of the individual’s cerebral cortex, then each parcel should have homogeneous connectivity and function. Furthermore, we also compared the relative utility of the different parcellation approaches for RSFC-based behavioral prediction.

### Resting-state connectional homogeneity

Resting-state connectional homogeneity was defined as the averaged Pearson’s correlations between rs-fMRI time courses of all pairs of vertices within each parcel, adjusted for parcel size and summed across parcels (Schaefer et al., 2018; Kong et al., 2019). Higher resting-state homogeneity means that vertices within the same parcel share more similar time courses. Therefore, higher resting-state homogeneity indicates better parcellation quality.

For each participant from the HCP test set (N = 755), we used one run to infer the individual-specific parcellation and computed resting-state homogeneity with the remaining 3 runs. For the MSC dataset (N = 9), we used one session to infer the individual-specific parcellation and computed resting-state homogeneity with the remaining 9 sessions (Figure 3A).

**Figure 3.**
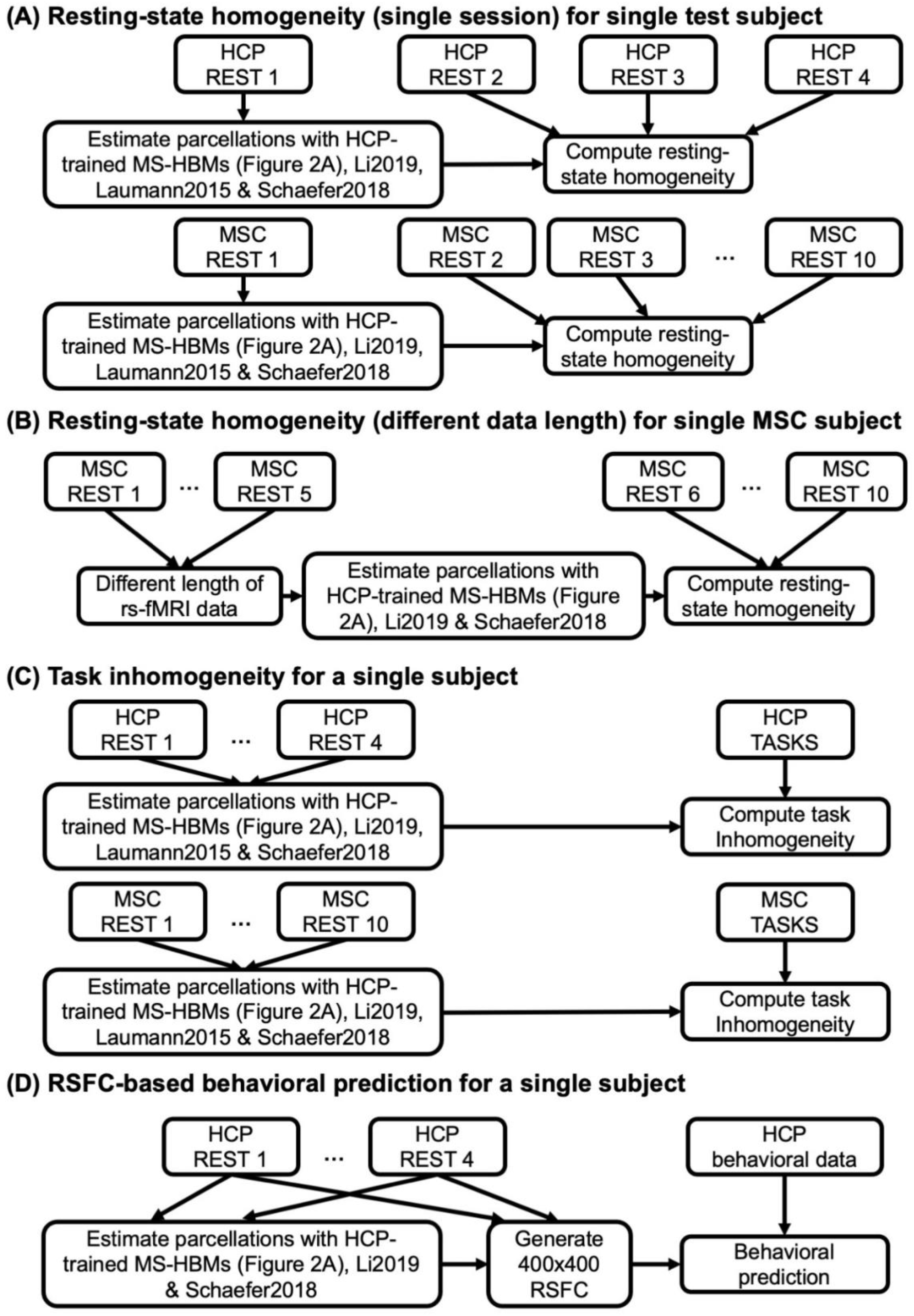
Flowcharts of comparisons with other algorithms. (A) Comparing out-of-sample resting-state homogeneity across different parcellation approaches applied to a single rs-fMRI session. (B) Comparing out-of-sample resting-state homogeneity across different parcellation approaches applied to different lengths of rs-fMRI data. (C) Comparing task inhomogeneity across different approaches. (D) Comparing RSFC-based behavioral prediction accuracies across different approaches. Across all analyses, MS-HBM parcellaions were estimated using the trained models from Figure 2A. We remind the reader that the trained MS-HBMs were estimated using the HCP training and validation sets (Figure 2A), which did not overlap with the HCP test set utilized in the current set of analyses. In the case of analyses (A) and (B), only a portion of rs-fMRI data was used to estimate the parcellations. The remaining rs-fMRI data was used to compute out-of-sample resting-state homogeneity. For analyses (C) and (D), all available rs-fMRI data was used to estimate the parcellations. Finally, we note that the local gradient approach (Laumann2015) does not yield a fixed number of parcels. Thus, the number of parcels is variable within an individual with different lengths of rs-fMRI data, so Laumann2015 was not considered for analysis B. Similarly, the number of parcels is different across participants, so the sizes of the RSFC matrices are different across participants. Therefore, Laumann2015 was also not utilized for analysis D.

**Figure 4.**
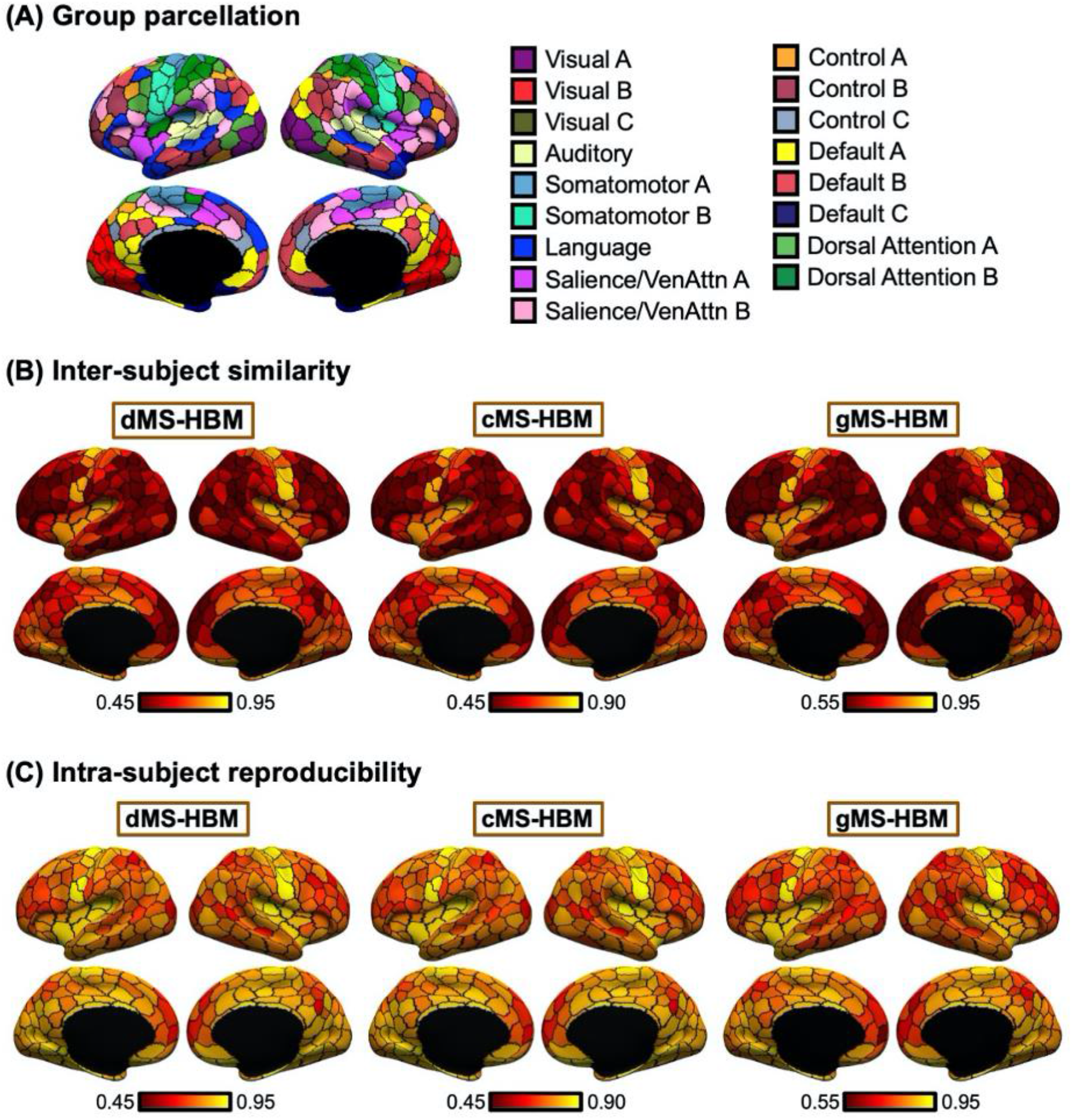
Individual-specific MS-HBM parcellations show high within-subject reproducibility and low across-subject similarity in the HCP test set. (A) 400-region Schaefer2018 group-level parcellation. (B) Inter-subject spatial similarity for different parcels. (C) Intra-subject reproducibility for different parcels. Yellow color indicates higher overlap. Red color indicates lower overlap. Individual-specific MS-HBM parcellations were generated by using day 1 (first two runs) and day 2 (last two runs) separately for each participant. Sensory-motor parcels exhibited higher intra-subject reproducibility and inter-subject similarity than association parcels.

Because MSC participants have large amount of rs-fMRI data (300 min), we also parcellated each MSC participant using different length of rs-fMRI data (10 min to 150 min) and evaluated the resting-state homogeneity with the remaining 5 sessions. This allowed us to estimate how much the algorithms would improve with more data (Figure 3B).

When comparing resting-state homogeneity between parcellations, the effect size (Cohen’s *d*) of differences and a two-sided paired-sample t-test (dof = 754 for HCP, dof = 8 for MSC) were computed.

### Task functional inhomogeneity

Task functional inhomogeneity was defined as the standard deviation of (activation) z-values within each parcel for each task contrast, adjusted for parcel size and summed across parcels (Gordon et al., 2017b; Schaefer et al., 2018). Lower task inhomogeneity means that activation within each parcel is more uniform. Therefore, lower task inhomogeneity indicates better parcellation quality. The HCP task-fMRI data consisted of 7 task domains: social cognition, motor, gambling, working memory, language processing, emotional processing, and relational processing (Barch et al., 2013). The MSC task-fMRI data consisted of three task domains: motor, mixed, and memory (Gordon et al., 2017b). Each task domain contained multiple task contrasts. All available task contrasts were utilized.

For each participant from the HCP test set (N = 755) and MSC dataset (N = 9), all rs-fMRI sessions were used to infer the individual-specific parcellation (Figure 3C). The individual-specific parcellation was then used to evaluate task inhomogeneity for each task contrast and then averaged across all available contrasts within a task domain, resulting in a single task inhomogeneity measure per task domain. When comparing between parcellations, we averaged the task inhomogeneity metric across all contrasts within a task domain before the effect size (Cohen’s *d*) of differences and a two-sided paired-sample t-test (dof = 754 for HCP, dof = 8 for MSC) were computed for each domain.

### Methodological considerations

It is important to note that a parcellation with more parcels tends to have smaller parcel size, leading to higher resting-state homogeneity and lower task inhomogeneity. For example, if a parcel comprised 2 vertices, then the parcel would be highly homogeneous. In our experiments, the MS-HBM algorithms and Li2019 were initialized with the 400-region Schaefer2018 group-level parcellation, resulting in the same number of parcels as Schaefer2018, i.e., 400 parcels. This allowed for a fair comparison among MS-HBMs, Li2019 and Schaefer2018.

However, parcellations estimated by Laumann2015 had a variable number of parcels across participants. Furthermore, Laumann2015 parcellations also had a significant number of vertices between parcels that were not assigned to any parcel, which has the effect of artificially increasing resting homogeneity and decreasing task inhomogeneity. Therefore, when comparing MS-HBM with Lauman2015 using resting-state homogeneity (Figure 3A) and task inhomogeneity (Figure 3C), we performed a post-hoc processing of MS-HBM parcellations to match the number of parcels and unlabeled vertices of Laumann2015 parcellations (Supplementary Methods S3).

In addition, the Laumann2015 approach yielded different numbers of parcels within an individual with different lengths of rs-fMRI data. Therefore, Laumann2015 was also excluded from the analysis of out-of-sample resting-state homogeneity with different lengths of rs-fMRI data (Figure 3B).

### RSFC-based behavioral prediction

Most studies utilized a group-level parcellation to derive RSFC for behavioral prediction (Dosenbach et al., 2010; Finn et al., 2015; Dubois et al., 2018; Li et al., 2019a; Weis et al., 2019). Here, we investigated if RSFC derived from individual-specific parcellations can improve behavioral prediction performance. As before (Kong et al., 2019; Li et al., 2019a; He et al., 2019), we considered 58 behavioral phenotypes measuring cognition, personality and emotion from the HCP dataset. Three participants were excluded from further analyses because they did not have all behavioral phenotypes, resulting in a final set of 752 test participants.

The different parcellation approaches were applied to each HCP test participant using all four rs-fMRI runs (Figure 3D). The Laumann2015 approach yielded parcellations with different numbers of parcels across participants, so there was a lack of inter-subject parcel correspondence. Therefore, we were unable to perform behavioral prediction with the Laumann2015 approach, so Laumann2015 was excluded from this analysis.

Given 400-region parcellations from different approaches (Schaefer2018; Li2019; dMS-HBM, cMS-HBM, gMS-HBM), functional connectivity was computed by correlating averaged time courses of each pair of parcels, resulting in a 400 × 400 RSFC matrix for each HCP test participant (Figure 3D). Consistent with our previous work (Kong et al., 2019; Li et al., 2019b; He et al., 2019), kernel regression was utilized to predict each behavioral measure in individual participants. Suppose *y* is the behavioral measure (e.g., fluid intelligence) and *FC* is the functional connectivity matrix of a test participant. In addition, suppose *y_i_* is the behavioral measure (e.g., fluid intelligence) and *FC_i_* is the individual-specific functional connectivity matrix of the *i*-th training participant. Then kernel regression would predict the behavior of the test participant as the weighted average of the behaviors of the training participants: *y* ≈ ∑_*i*∈training set_Similarity(*FC_i_*, *FC*)*y_i_*. Here, Similarity(*FC_i_*, *FC*) is the Pearson’s correlation between the functional connectivity matrices of the *i*-th training participant and the test participant. Because the functional connectivity matrices were symmetric, only the lower triangular portions of the matrices were considered when computing the correlation. Therefore, kernel regression encodes the intuitive idea that participants with more similar RSFC patterns exhibited similar behavioral measures.

In practice, an *l*_2_-regularization term (i.e., kernel ridge regression) was included to reduce overfitting (Supplementary Methods S4; Murphy, 2012). We performed 20-fold cross-validation for each behavioral phenotype. Family structure within the HCP dataset was taken into account by ensuring participants from the same family (i.e., with either the same mother ID or father ID) were kept within the same fold and not split across folds. For each test fold, an inner-loop 20-fold cross-validation was repeatedly applied to the remaining 19 folds with different regularization parameters. The optimal regularization parameter from the inner-loop cross-validation was then used to predict the behavioral phenotype in the test fold. Accuracy was measured by correlating the predicted and actual behavioral measure across all participants within the test fold (Finn et al., 2015; Kong et al., 2019; Li et al., 2019b). By repeating the procedure for each test fold, each behavior yielded 20 correlation accuracies, which were then averaged across the 20 folds. Because a single 20-fold cross-validation might be sensitive to the particular split of the data into folds (Varoquaux et al., 2017), the above 20-fold cross-validation was repeated 100 times. The mean accuracy and standard deviation across the 100 cross-validations will be reported. When comparing between parcellations, a corrected resampled t-test for repeated k-fold cross-validation was performed (Bouckaert and Frank, 2004). We also repeated the analyses using coefficient of determination (COD) as a metric of prediction performance.

As certain behavioral measures are known to correlate with motion (Siegel et al., 2016), we regressed out age, sex, framewise displacement (FD), DVARS, body mass index and total brain volume from the behavioral data before kernel ridge regression. To prevent any information leak from the training data to test data, the nuisance regression coefficients were estimated from the training folds and applied to the test fold.

### Code and data availability

Code for this work is freely available at the GitHub repository maintained by the Computational Brain Imaging Group (https://github.com/ThomasYeoLab/CBIG). The Schaefer2018 group-level parcellation and code are available here (https://github.com/ThomasYeoLab/CBIG/tree/master/stable_projects/brain_parcellation/Schaefer2018_LocalGlobal), while the areal-level MS-HBM parcellation code is available here (https://github.com/ThomasYeoLab/CBIG/tree/master/stable_projects/brain_parcellation/Kong2022_ArealMSHBM). We have also provided trained MS-HBM parameters at different spatial resolutions, ranging from 100 to 1000 parcels.

We note that the computational bottleneck for gMS-HBM is the computation of the local gradients (Laumann et al., 2015). We implemented a faster and less memory-intensive version of the local gradient computation by subsampling the functional connectivity matrices (Supplementary Methods S1.3). Computing the gradient map of a single HCP run requires 15 minutes and 3 GB of RAM, compared with 4 hours and 40 GB of RAM in the original version. The resulting gradient maps were highly similar to the original gradient maps (r = 0.97). The faster gradient code can be found here (https://github.com/ThomasYeoLab/CBIG/tree/master/utilities/matlab/speedup_gradients).

The individual-specific parcellations for the HCP and MSC, together with the associated RSFC matrices, are available here (https://balsa.wustl.edu/study/show/Pr8jD and https://github.com/ThomasYeoLab/CBIG/tree/master/stable_projects/brain_parcellation/Kong2022_ArealMSHBM).

## Results

### Overview

Three variations of the MS-HBM with different contiguity constraints (Figure 1) were applied to two multi-session rs-fMRI datasets to ensure that the approaches were generalizable across datasets with significant acquisition and processing differences. After confirming previous literature (Mueller et al., 2013; Laumann et al., 2015; Kong et al., 2019) that inter-subject and intra-subject RSFC variabilities were different across the cortex, we then established that the MS-HBM algorithms produced individual-specific areal-level parcellations with better quality than other approaches. Finally, we investigated whether RSFC derived from MS-HBM parcellations could be used to improve behavioral prediction.

### Sensory-motor cortex exhibits lower inter-subject but higher intra-subject functional connectivity variability than association cortex

The parameters of gMS-HBM, dMS-HBM and cMS-HBM were estimated using the HCP training set. Figure S1 shows the inter-subject RSFC variability (*ϵ_l_*) and intra-subject RSFC variability (*σ_l_*) overlaid on corresponding Schaefer2018 group-level parcels. The pattern of inter-subject and intra-subject RSFC variability were consistent with previous work (Mueller et al., 2013; Laumann et al., 2015; Kong et al., 2019). More specifically, sensory-motor parcels exhibited lower inter-subject RSFC variability than association cortical parcels. On the other hand, association cortical parcels exhibited lower intra-subject RSFC variability than sensory-motor parcels.

### Individual-specific MS-HBM parcellations exhibit high intra-subject reproducibility and low inter-subject similarity

To assess intra-subject reproducibility and inter-subject similarity, the three MS-HBM variants were tuned on the HCP training and validation sets, and then applied to the HCP test set. Individual-specific parcellations were generated by using resting-state fMRI data from day 1 (first 2 runs) and day 2 (last 2 runs) separately for each participant. All 400 parcels were present in 99% of the participants.

Figure 4 shows the inter-subject and intra-subject spatial similarity (Dice coefficient) of parcels from the three MS-HBM variants in the HCP test set. Intra-subject reproducibility was greater than inter-subject similarity across all parcels. Consistent with our previous work on individual-specific cortical networks (Kong et al., 2019), sensory-motor parcels were more spatially similar across participants than association cortical parcels. Sensory-motor parcels also exhibited greater within-subject reproducibility than association cortical parcels.

Overall, gMS-HBM, dMS-HBM and cMS-HBM achieved intra-subject reproducibility of 81.0%, 80.4% and 76.1% respectively, and inter-subject similarity of 68.2%, 68.1% and 63.9% respectively. We note that these metrics cannot be easily used to judge the quality of the parcellations. For example, gMS-HBM has higher intra-subject reproducibility and higher inter-subject similarity than cMS-HBM, so we cannot simply conclude that one is better than the other.

Figures 5A and S2 show the gMS-HBM parcellations of 4 representative HCP participants. Figures S3 and S4 show the dMS-HBM and cMS-HBM parcellations of the same HCP participants. Consistent with previous studies of individual-specific parcellations (Glasser et al., 2016; Chong et al., 2017; Gordon et al., 2017b; Salehi et al., 2018; Li et al., 2019b; Seitzman et al., 2019), parcel shape, size, location and topology were variable across participants. Parcellations were highly similar within each participant with individual-specific parcel features highly preserved across sessions (Figure 5B). Similar results were obtained with dMS-HBM and cMS-HBM (Figure 5B).

**Figure 5.**
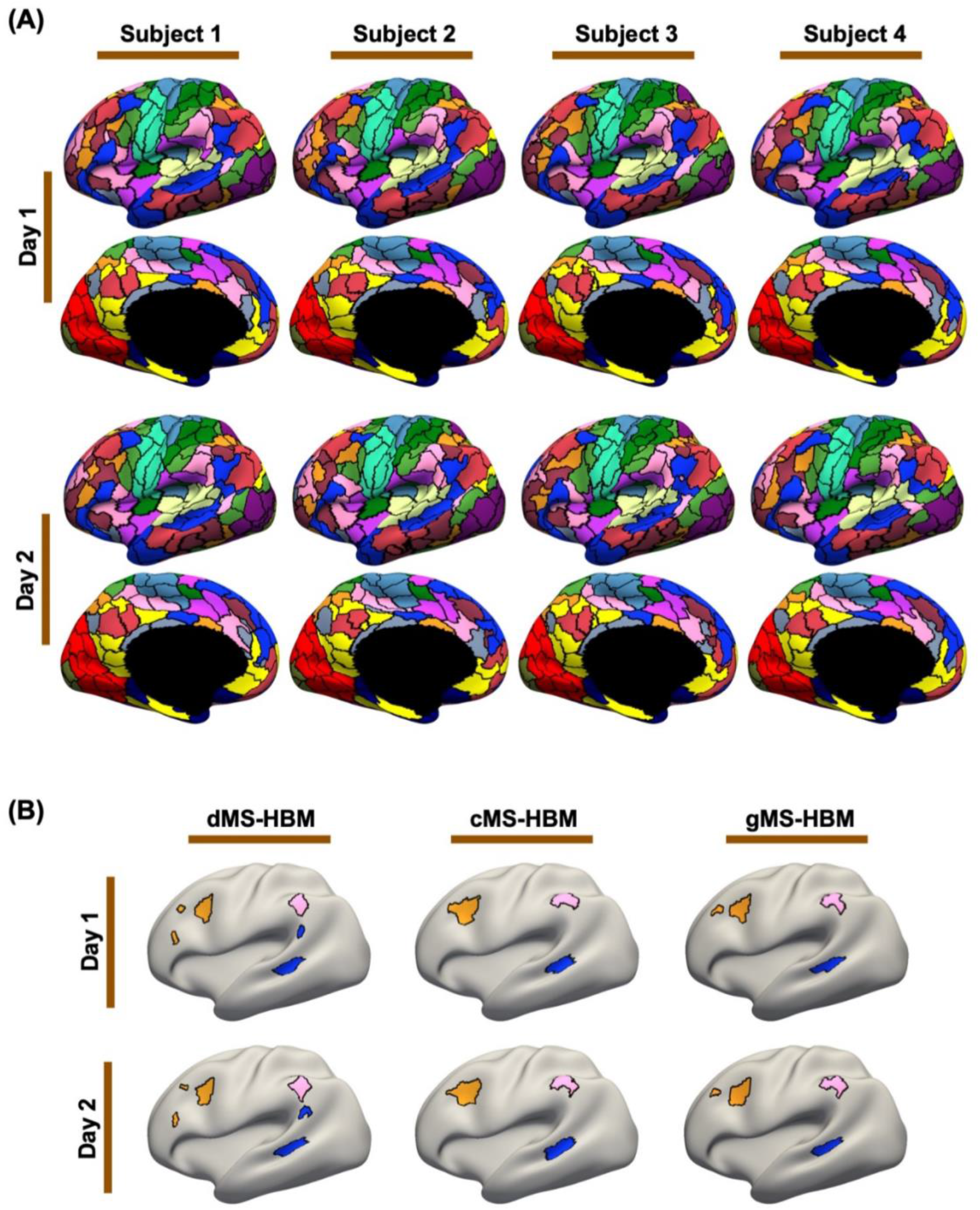
MS-HBM parcellations exhibit individual-specific features that are replicable across sessions. (A) 400-region individual-specific gMS-HBM parcellations were estimated using rs-fMRI data from day 1 and day 2 separately for each HCP test participant. Right hemisphere parcellations are shown in Figure S2. See Figure S3 and S4 for dMS-HBM and cMS-HBM. (B) Replicable individual-specific parcellation features in a single HCP test participant for dMS-HBM, cMS-HBM and gMS-HBM.

The trained MS-HBM from the HCP dataset was also applied to the MSC dataset. The MS-HBM parcellations of 4 representative MSC participants are shown in Figures S5, S6 and S7. Similar to the HCP dataset, the parcellations also captured unique features that were replicable across the first five sessions and the last five sessions. Overall, gMS-HBM, dMS-HBM and cMS-HBM achieved intra-subject reproducibility of 75.5%, 73.9% and 67.8% respectively, and inter-subject similarity of 50.6%, 47.1% and 42.9% respectively.

### Geometric properties of MS-HBM parcellations

In the HCP test set, the average number of spatially disconnected components per parcel was 1.95 ± 0.29 (mean ± std), 1 ± 0 and 1.06 ± 0.07 for dMS-HBM, cMS-HBM and gMS-HBM respectively. In the case of dMS-HBM, the maximum number of spatially disconnected components (across all participants and parcels) was 11 (Figure S8). In the case of gMS-HBM, the maximum number of spatially disconnected components (across all participants and parcels) was 3 (Figure S8). On the other hand, the average roundness of the parcellations was 0.56 ± 0.02 (mean ± std), 0.60 ± 0.01 and 0.58 ± 0.02 for dMS-HBM, cMS-HBM and gMS-HBM respectively. Overall, gMS-HBM parcels have much fewer spatially disconnected components than dMS-HBM, while achieving intermediate roundness between dMS-HBM and cMS-HBM.

### Individual-specific MS-HBM parcels exhibit higher resting-state homogeneity than other approaches

Individual-specific areal-level parcellations were estimated using a single rs-fMRI session for each HCP test participant and each MSC participant. Resting-state homogeneity was evaluated using leave-out sessions in the HCP (Figures 6A and 6B) and MSC (Figures 6C, 6D and S9) datasets. We note that comparisons with Laumann2015 are shown on separate plots (Figures 6B and 6D) because Laumann2015 yielded different number of parcels across participants. Therefore, we matched the number of MS-HBM parcels to Laumann2015 for each participant for fair comparison (see Methods).

**Figure 6.**
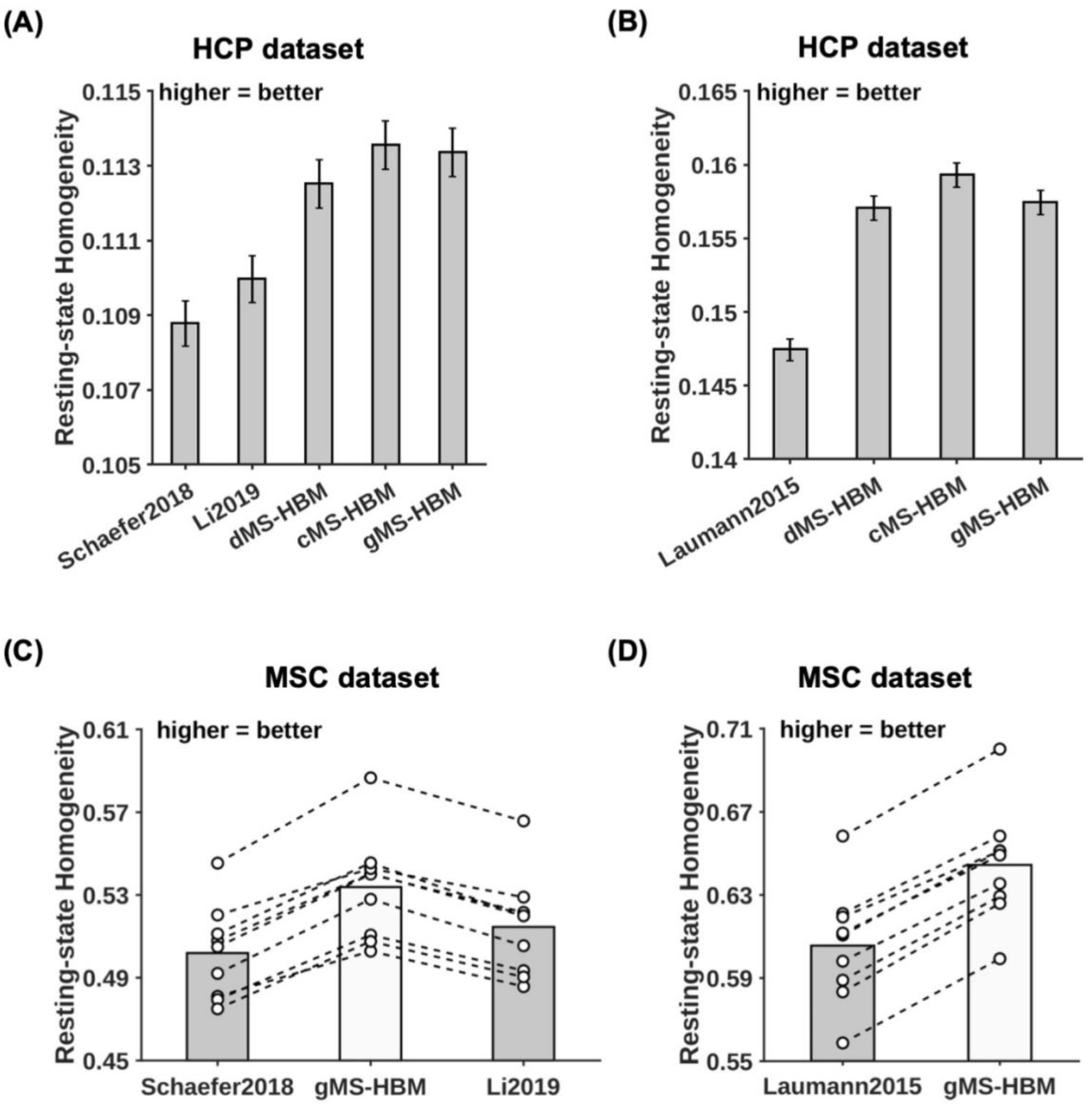
MS-HBM parcellations achieved better out-of-sample resting-state homogeneity than other approaches. (A) 400-region individual-specific parcellations were estimated using a single rs-fMRI session and resting-state homogeneity was computed on the remaining sessions for each HCP test participant. Error bars correspond to standard errors. (B) Same as (A) except that Laumann2015 allowed different number of parcels across participants, so we matched the number of MS-HBM parcels to Laumann2015 for each participant. Therefore, the numbers for (A) and (B) were not comparable. (C) 400-region individual-specific parcellations were estimated using a single rs-fMRI session and resting-state homogeneity was computed on the remaining sessions for each MSC participant. Each circle represents one MSC participant. Dash lines connect the same participants. (D) Same as (C) except that Laumann2015 allowed different number of parcels across participants, so we matched the number of MS-HBM parcels to Laumann2015 for each participant. Results for dMS-HBM and cMS-HBM in the MSC dataset are shown in Figure S9.

Across both HCP and MSC datasets, the MS-HBM algorithms achieved better homogeneity than the group-level parcellation (Schaefer2018) and two individual-specific areal-level parcellation approaches (Laumann2015 and Li2019). Compared with Schaefer2019, the three MS-HBM variants achieved an improvement ranging from 3.4% to 7.5% across the two datasets (average improvement = 5.2%, average Cohen’s d = 3.8, largest p = 1.9e-6). Compared with Li2019, the three MS-HBM variants achieved an improvement ranging from 2.2% to 4.9% across the two datasets (average improvement = 3.4%, average Cohen’s d = 3.9, largest p = 5.5e-6). Compared with Laumann2015, the three MS-HBM variants achieved an improvement ranging from 6.3% to 7.8% across the two datasets (average improvement = 6.7%, average Cohen’s d = 7.5, largest p = 1.2e-9). All reported p values were significant after correcting for multiple comparisons with false discovery rate (FDR) q < 0.05.

Among the three MS-HBM variants, cMS-HBM achieved the highest homogeneity, while dMS-HBM was the least homogeneous. In the HCP dataset, cMS-HBM achieved an improvement of 0.19% (Cohen’s d = 0.5, p = 3.5e-38) over gMS-HBM, and gMS-HBM achieved an improvement of 0.76% (Cohen’s d = 2.5, p = 3.5e-38) over dMS-HBM. In the MSC dataset, cMS-HBM achieved an improvement of 1.1% (Cohen’s d = 3.5, p = 6.3e-6) over gMS-HBM, and gMS-HBM achieved an improvement of 0.7% (Cohen’s d = 2.6, p = 6.1e-5) over dMS-HBM. All reported p values were significant after correcting for multiple comparisons with FDR q < 0.05.

Individual-specific parcellations were estimated with increasing length of rs-fMRI data in the MSC dataset. Resting-state homogeneity was evaluated using leave-out sessions (Figures 7A and S10). We note that Laumann2015 parcellations had different number of parcels with different length of rs-fMRI data. Therefore, the resting-state homogeneity of Laumann2015 parcellations was not comparable across different length of rs-fMRI data, so the results were not shown. Because Schaefer2018 is a group-level parcellation, the parcellation stays the same regardless of the amount of data. Therefore, the performance of the Schaefer2018 group-level parcellation remained constant regardless of the amount of data. Surprisingly, the performance of the Li2019 individual-specific parcellation approach also remained almost constant regardless of the amount of data. One possible reason is that Li2019 constrained individual-specific parcels to overlap with group-level parcels. This might be an overly strong constraint, which could not be overcome with more rs-fMRI data. By contrast, the MS-HBM algorithms (dMS-HBM, cMS-HBM, and gMS-HBM) exhibited higher homogeneity with increased length of rs-fMRI data, suggesting that MS-HBM models were able to improve with more rs-fMRI data.

**Figure 7.**
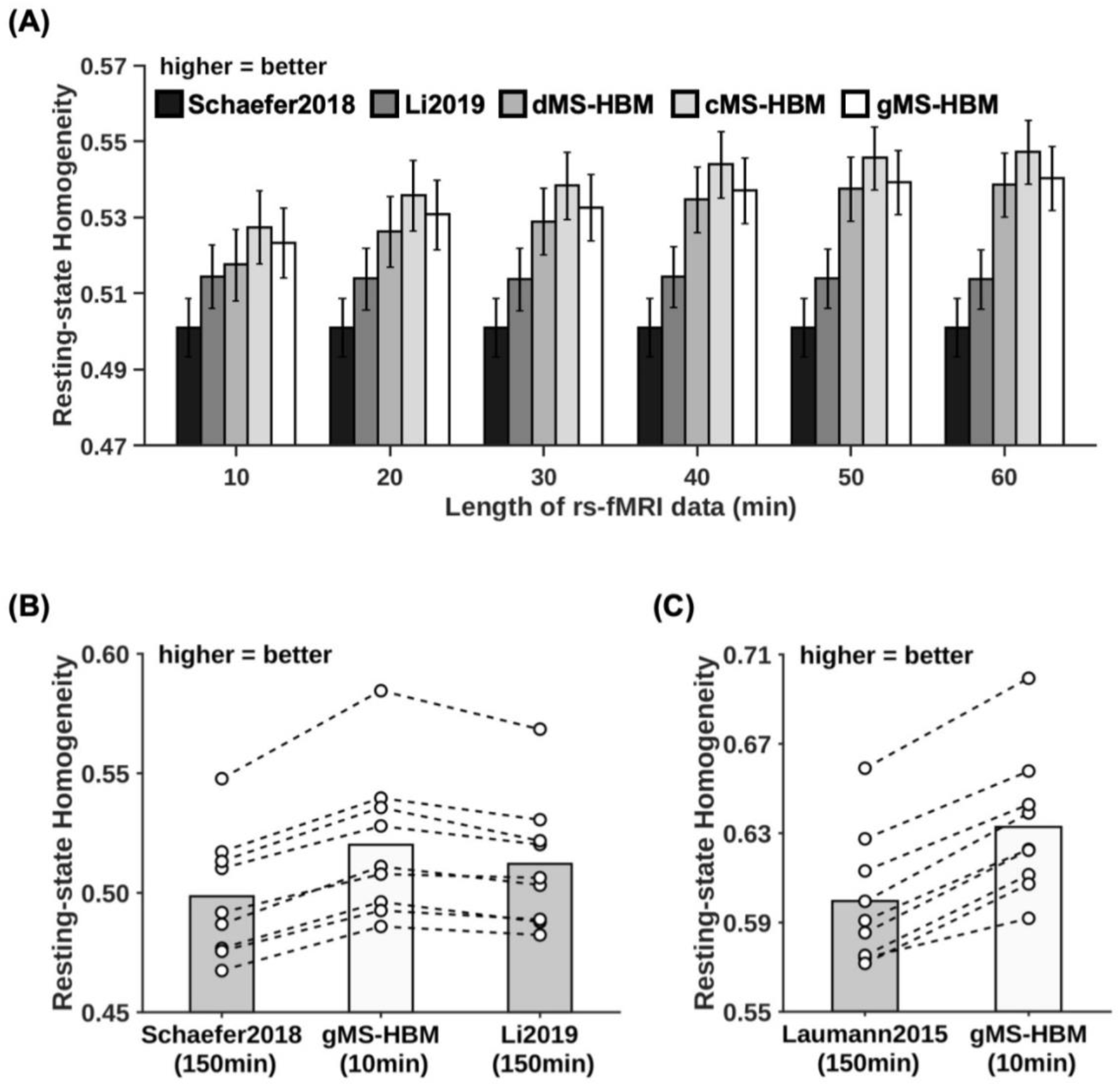
MS-HBM parcellations achieved better out-of-sample resting-state homogeneity with less amount of data. (A) 400-region individual-specific parcellations were estimated using different lengths of rs-fMRI data for each MSC participant. Resting-state homogeneity was evaluated using leave-out sessions. Error bars correspond to standard errors. (B) 400-region individual-specific parcellations were estimated for each MSC participant using 10 min of rs-fMRI data for gMS-HBM and 150 min of rs-fMRI data for Li2019. Each circle represents one MSC participant. Dash lines connect the same participants. (C) Same as (B) except that Laumann2015 yielded different number of parcels for each participant, so we matched the number of MS-HBM parcels accordingly for each participant. Results for dMS-HBM and cMS-HBM are shown in Figure S10.

Furthermore, using just 10 min of rs-fMRI data, the MS-HBM algorithms achieved better homogeneity than Laumann2015 and Li2019 using 150 min of rs-fMRI data (Figures 7B and S10). More specifically, compared with Laumann2015 using 150 min of rs-fMRI data, dMS-HBM, cMS-HBM, and gMS-HBM using 10 min of rs-fMRI data achieved an improvement of 2.6% (Cohen’s d = 2.7, p = 3.6e-5), 6.2% (Cohen’s d = 5.5, p = 1.9e-7) and 5.6% (Cohen’s d = 6.1, p = 2.3e-7) respectively. Compared with Li2019 using 150 min of rs-fMRI data, dMS-HBM, cMS-HBM, and gMS-HBM using 10 min of rs-fMRI data achieved an improvement of 0.4% (Cohen’s d = 0.4, not significant), 2.4% (Cohen’s d = 1.9, p = 4.3e-4) and 1.5% (Cohen’s d = 1.7, p = 1.0e-3) respectively. All reported p values were significant after correcting for multiple comparisons with FDR q < 0.05.

### Individual-specific MS-HBM parcels exhibit lower task inhomogeneity than other approaches

Individual-specific parcellations were estimated using all rs-fMRI sessions from the HCP test set and MSC dataset. Task inhomogeneity was evaluated using task fMRI. Figures 8 and S11 show the task inhomogeneity of all approaches for all task domains in the MSC and HCP datasets respectively. Compared with Schaefer2019, the three MS-HBM variants achieved an improvement ranging from 0.9% to 5.9% across all task domains and datasets (average improvement = 3.2%, average Cohen’s d = 2.4, largest p = 2.0e-3). Compared with Li2019, the three MS-HBM variants achieved an improvement ranging from 0.8% to 5.0% across all task domains and datasets (average improvement = 2.7%, average Cohen’s d = 2.2, largest p = 1.8e-3). Compared with Laumann2015, the three MS-HBM variants achieved an improvement ranging from 1.9% to 28.1% across all task domains and datasets (average improvement = 6.7%, average Cohen’s d = 2.3, largest p = 0.017). All reported p values were significant after correcting for multiple comparisons with false discovery rate (FDR) q < 0.05. In the case of MSC, these improvements were observed in almost every single participant across all tasks (Figure 8).

**Figure 8.**
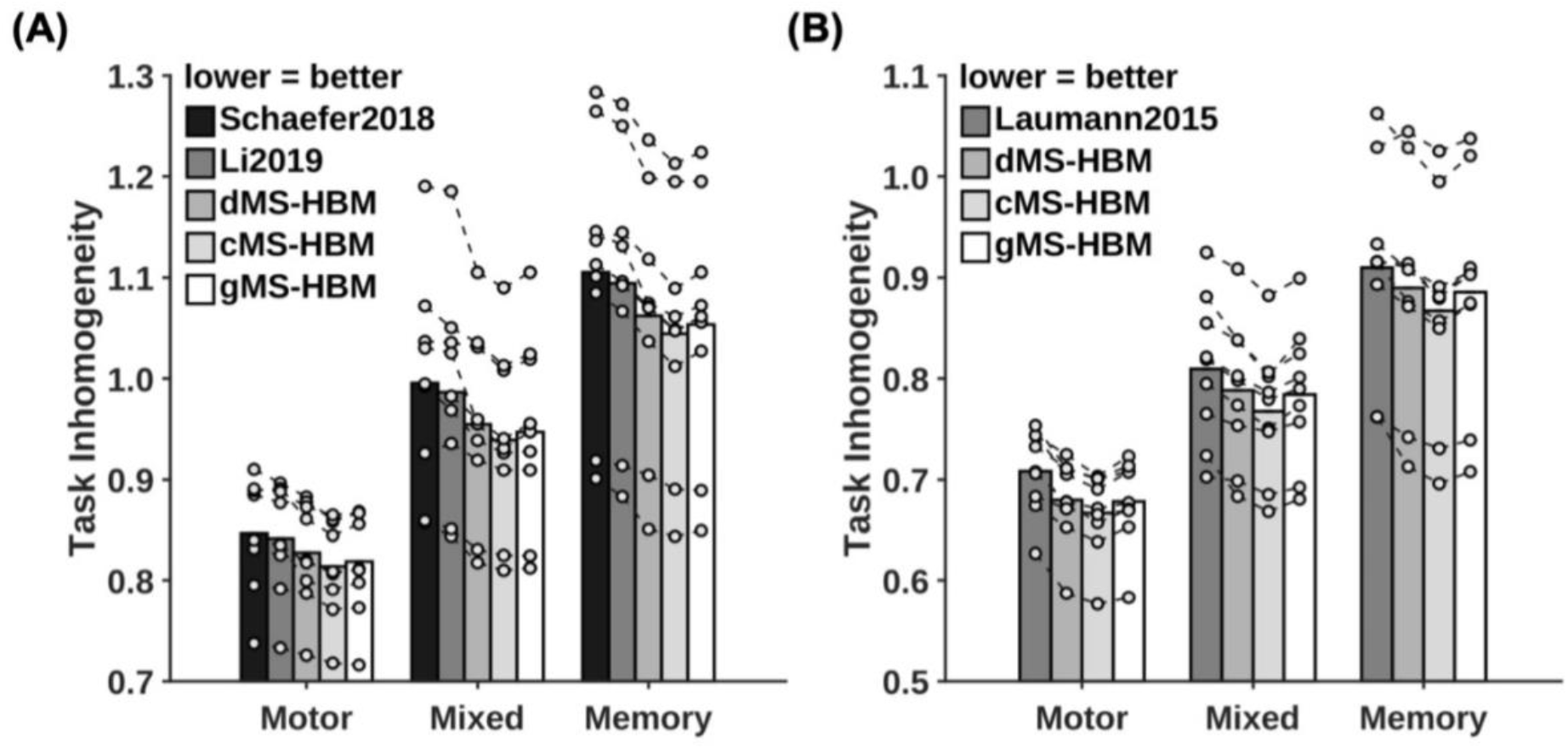
MS-HBM parcellations achieved better task inhomogeneity in the MSC dataset. (A) 400-region individual-specific parcellations were estimated using all resting-state fMRI sessions. Task inhomogeneity was evaluated using task fMRI. Task inhomogeneity was then defined as the standard deviation of task activation within each parcel, and then averaged across all parcels and contrasts within each behavioral domain. Lower value indicates better task inhomogeneity. Each circle represents one MSC participant. Dash lines connect the same participants. (B) Same as (A) except that Laumann2015 yielded different number of parcels for each participant, so we matched the number of MS-HBM parcels accordingly for each participant. HCP results are shown in Figure S11.

Among the three MS-HBM variants, cMS-HBM achieved the best task inhomogeneity, while dMS-HBM achieved the worst task inhomogeneity. Compared with gMS-HBM, cMS-HBM achieved an improvement ranging from 0.03% to 0.92% across all task domains and datasets (average improvement = 0.3%, average Cohen’s d = 0.6, largest p = 0.013). Compared with dMS-HBM, gMS-HBM achieved an improvement ranging from 0.06% to 1.1% across all task domains and datasets (average improvement = 0.5%, average Cohen’s d = 1.1, largest p = 1.2e-3). All reported p values were significant after correcting for multiple comparisons with FDR q < 0.05.

### Functional connectivity of individual-specific MS-HBM parcels improves behavioral prediction

Individual-specific parcellations were estimated using all rs-fMRI sessions from the HCP test set. The RSFC of the individual-specific parcellations was used for predicting 58 behavioral measures. We note that the number of parcels was different across participants for Laumann2015, so Laumann2015 could not be included for this analysis.

Tables S2 and S3 summarize the average prediction accuracies (Pearson’s correlation) for different sets of behavioral measures, including cognitive, personality and emotion measures. Overall, individual-specific functional connectivity strength from MS-HBM parcellations achieved better prediction performance than other approaches. In general, gMS-HBM achieved better prediction performance than dMS-HBM and cMS-HBM, but differences were not significant.

Figure 9A shows the average prediction accuracies of all 58 behaviors across different parcellation approaches. Compared with Schaefer2018 and Li2019, gMS-HBM achieved improvements of 16% (p = 5.0e-4) and 18% (p = 5.4e-4) respectively. Both p values remained significant after correcting for multiple comparisons with FDR q < 0.05. Compared with cMS-HBM and dMS-HBM, gMS-HBM achieved an improvement of 5.5% and 3.4% respectively. However, differences among MS-HBM variants were not significant.

**Figure 9.**
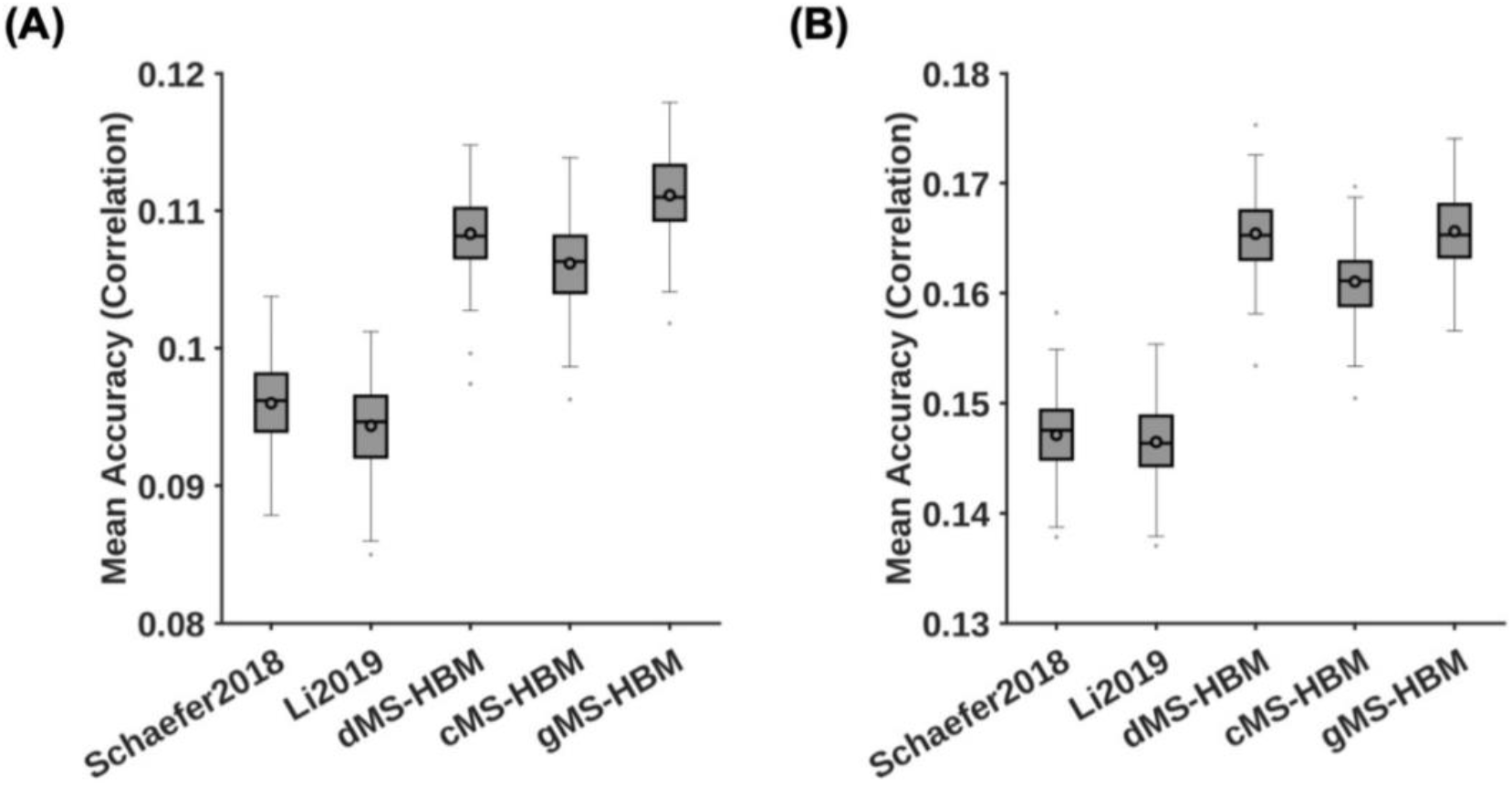
MS-HBM achieves the best behavioral prediction performance as measured by Pearson’s correlation. (A) Average prediction accuracies (Pearson’s correlation) of all 58 behavioral measures. Boxplots utilized default Matlab parameters, i.e., box shows median and inter-quartile range (IQR). Whiskers indicate 1.5 IQR (not standard deviation). Circle indicates mean. dMS-HBM, cMS-HBM and gMS-HBM achieved average prediction accuracies of r = 0.1083 ± 0.0031 (mean ± std), 0.1062 ± 0.0031 and 0.1111 ± 0.0031 respectively. On the other hand, Schaefer2018 and Li2019 achieved average prediction accuracies of r = 0.0960 ± 0.0031 and 0.0944 ± 0.0031 respectively. (B) Average prediction accuracies (Pearson’s correlation) of 36 behavioral measures with accuracies (Pearson’s correlation) higher than 0.1 for at least one approach (“36 behaviors > 0.1”). dMS-HBM, cMS-HBM and gMS-HBM achieved average prediction accuracies of r = 0.1630 ± 0.0034 (mean ± std), 0.1590 ± 0.0035 and 0.1656 ± 0.0036 respectively. On the other hand, Schaefer2018 and Li2019 achieved average prediction accuracies of r = 0.1442 ± 0.0036 and 0.1444 ± 0.0035 respectively.

We note that some behavioral measures were predicted poorly by all approaches. This is not unexpected because we do not expect all behavioral measures to be predictable with RSFC. Therefore, we further consider a subset of behavioral measures that could be predicted well by at least one approach. Figure 9B shows the average prediction accuracies of 36 behaviors with accuracies higher than 0.1 for at least one approach (“36 behaviors > 0.1”). Compared with Schaefer2018 and Li2019, gMS-HBM achieved improvements of 13% (p = 2.2e-4) and 13% (p = 4.5e-4) respectively. All p values remained significant after correcting for multiple comparisons with FDR q < 0.05. Differences among MS-HBM variants were again not significant. Similar conclusions were obtained with COD instead of correlations (Figure 10; Tables S4 and S5).

**Figure 10.**
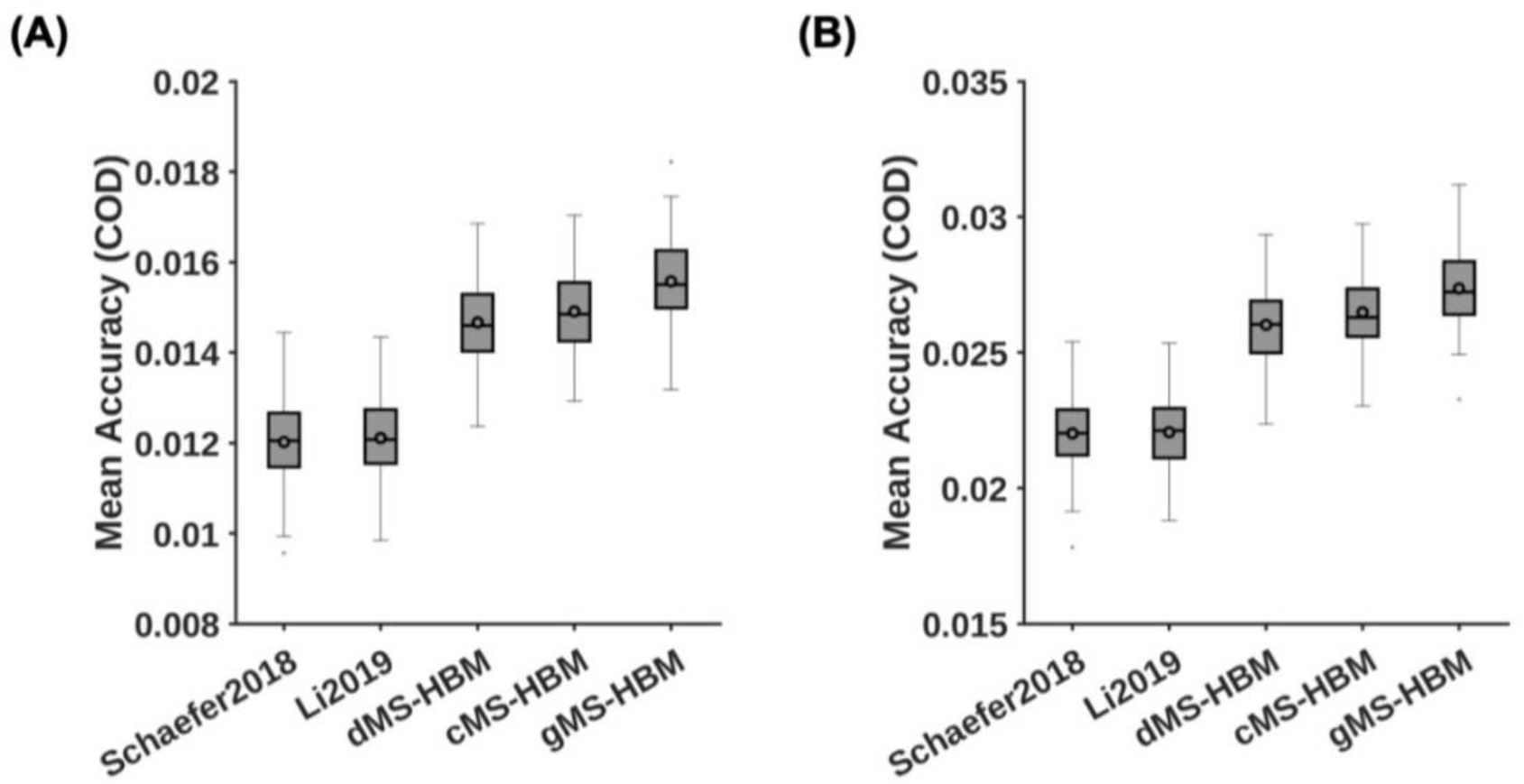
MS-HBM achieves the best behavioral prediction performance as measured by coefficient of determination (COD). (A) Average prediction accuracies (COD) of all 58 behavioral measures. Boxplots utilized default Matlab parameters, i.e., box shows median and inter-quartile range (IQR). Whiskers indicate 1.5 IQR (not standard deviation). Circle indicates mean. dMS-HBM, cMS-HBM and gMS-HBM achieved average prediction accuracies (COD) = 0.0147 ± 0.0009 (mean ± std), 0.0149 ± 0.0009 and 0.0156 ± 0.0010 respectively. On the other hand, Schaefer2018 and Li2019 achieved average prediction accuracies (COD) = 0.0120 ± 0.0009 and 0.0121 ± 0.0009 respectively. (B) Average prediction accuracies (COD) of 36 behavioral measures with accuracies (Pearson’s correlation) greater than 0.1 for at least one approach (“36 behaviors > 0.1”). dMS-HBM, cMS-HBM and gMS-HBM achieved average prediction accuracies (COD) = 0.0252 ± 0.0014 (mean ± std), 0.0257 ± 0.0014 and 0.0266 ± 0.0014 respectively. On the other hand, Schaefer2018 and Li2019 achieved average prediction accuracies (COD) = 0.0212 ± 0.0014 and 0.0213 ± 0.0014 respectively.

### Task performance measures are more predictable than self-reported measures

To explore which behavioral measures can be consistently predicted well regardless of parcellations, we ordered the 58 behavioral measures based on averaged prediction accuracies (Pearson’s correlation) across Schaefer2018, Li2019 and the three MS-HBM variants (Figure 11B). Our previous studies (Li et al., 2019a; Liégeois et al., 2019) have suggested that “self-reported” and “task performance” measures might be differentially predicted under different conditions. Using the same classification of behavioral measures (Li et al., 2019a; Liégeois et al., 2019), we found that the average prediction accuracies of self-reported measures and task performance measures were r = 0.0890 ± 0.0048 and r = 0.1181 ± 0.0033 respectively (Figure 11A), suggesting that on average, task performance measures were more predictable than self-reported measures (p = 0.042).

**Figure 11.**
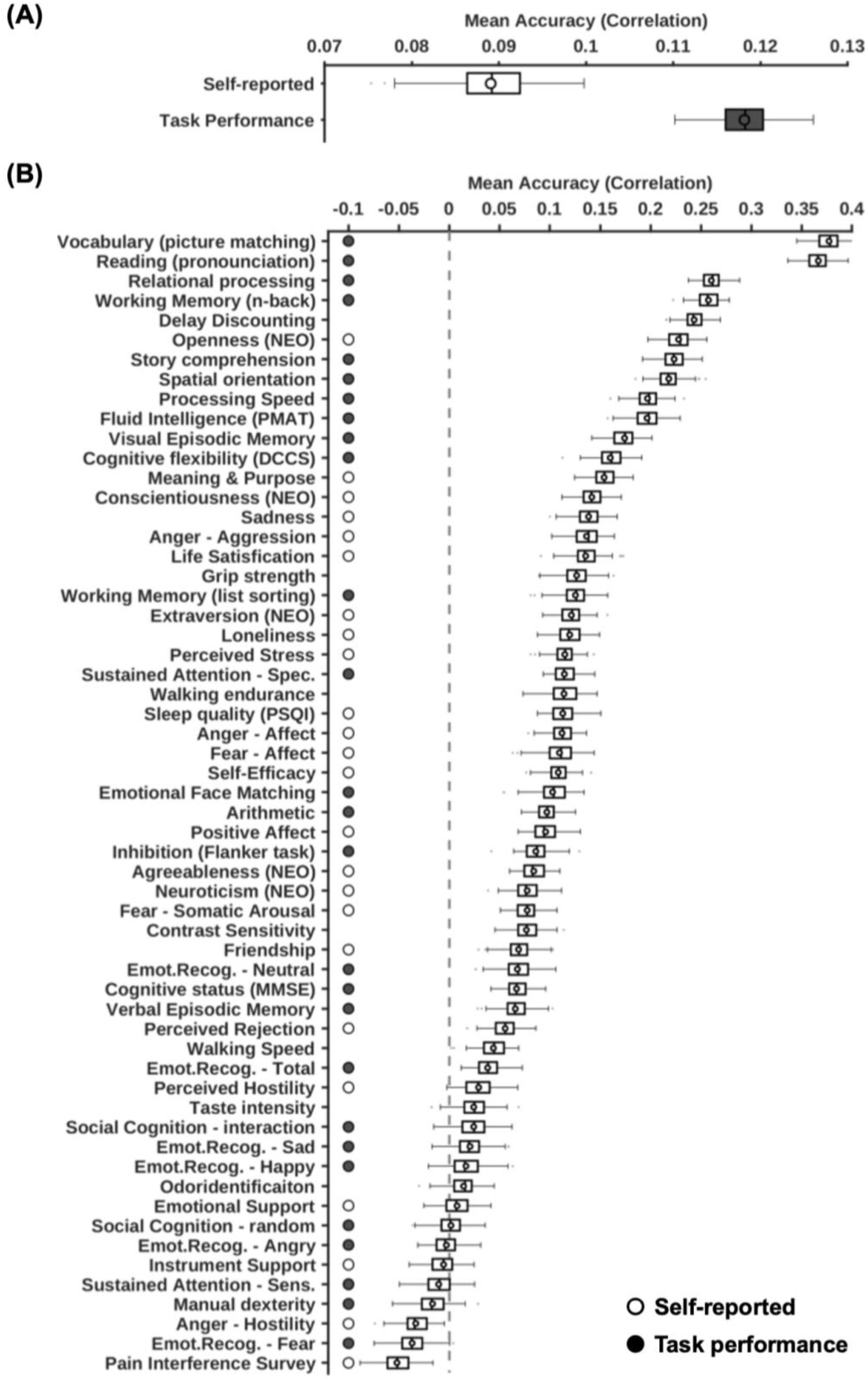
Task performance measures were predicted better than self-reported measures across different parcellation approaches. Prediction accuracies were averaged across all parcellation approaches (three MS-HBM variants, Schaefer2018, and Li2019). (A) Prediction accuracies averaged across HCP task-performance measures (gray) and HCP self-reported measures (white). (B) Behavioral measures were ordered based on average prediction accuracies. Gray color indicates task performance measures. White color indicates self-reported measures. Boxplots utilized default Matlab parameters, i.e., box shows median and inter-quartile range (IQR). Whiskers indicate 1.5 IQR (not standard deviation). Circle indicates mean. Designation of behavioral measures into “self-reported” and “task-performance) measures followed previous studies (Li et al., 2019a; Liégeois et al., 2019).

## Discussion

In this manuscript, we demonstrated the robustness of the MS-HBM areal-level parcellation approach. Compared with a group-level parcellation and two state-of-the-art individual-specific areal-level parcellation approaches, we found that MS-HBM parcels were more homogeneous during resting-state while also exhibiting more uniform task activation patterns (i.e., lower task inhomgeneity). Furthermore, RSFC derived from individual-specific MS-HBM parcellations achieved better behavioral prediction performance than other approaches. Among the three MS-HBM variants, the contiguous MS-HBM (cMS-HBM) exhibited the best resting homogeneity and task inhomogeneity, while the gradient-infused MS-HBM (gMS-HBM) exhibited the best behavioral prediction performance.

### Interpretation of the MS-HBM areal-level parcellations

Previous studies have estimated around 300 to 400 classically defined cortical areas in the human cerebral cortex (Van Essen et al. 2012b). Therefore, various groups (including ours) have most frequently utilized the 400-region Schaefer group-level parcellation (Varikuti et al., 2018; Franzmeier et al., 2019; Kebets et al., 2019; Murphy et al., 2020; Orban et al., 2020). Other studies have opted to utilize different resolutions of the Schaefer group-level parcellation, e.g., 100 regions (Chin Fatt et al., 2019), 200 regions (Anderson et al., 2020; Faskowitz et al., 2020) and 800 regions (Valk et al., 2020). Despite our focus on the 400-region areal-level parcellations in the current study, we do not believe that there is an optimal number of cortical parcels because of the multi-resolution organization of the cerebral cortex (Churchland and Sejnowski, 1988; van den Heuvel and Yeo, 2017). Indeed, given the heterogeneity of cortical areas (Kaas, 1987; Amunts and Zilles, 2015), cortical areas might be further subdivided into meaningful computational sub-units.

More specifically and consistent with other studies, our areal-level parcels likely captured sub-areal features such as somatotopy and visual eccentricity (Gordon et al., 2016; Schaefer et al., 2018). Ultimately, the choice of parcellation resolution might depend on the specific application. For example, a recent study suggested that brain-behavior relationships are scale-dependent (Betzel et al., 2019). Furthermore, a higher resolution parcellation might be computationally infeasible for certain analysis, such as edge-centric network analysis (Faskowitz et al., 2020). Therefore, we have provided trained MS-HBM at different spatial resolutions, ranging from 100 to 1000 parcels. It is worth noting that because our parcels do not correspond to traditional cortical areas (Kaas, 1987; Amunts and Zilles, 2015), we have been careful to avoid the term “areas”. Instead we use the term “areal-level parcellation” when referring to the entire parcellation and “parcels” when referring to individual regions throughout the manuscript.

Several studies have shown that brain networks reconfigure during tasks (Cole et al., 2014; Krienen et al., 2014; Salehi et al., 2019). Consequently, some have questioned the existence of a single individual-specific areal-level parcellation that generalizes across resting and task states (Salehi et al., 2019). While we do not contest the results of Salehi and colleagues, we have a very different interpretation. Cortical areas (e.g., V1) are conceptualized as representing stable computational units (Felleman and Van Essen, 1991). Consequently, their boundaries should remain the same regardless of transient task states across the span of a few days, even if long-term experiences can potentially shape the development and formation of cortical areas (Arcaro et al., 2017; Gomez et al., 2019). Thus, the results of Salehi and colleagues do not rule out the plausibility of estimating a stable individual-level areal-level parcellation with rs-fMRI data alone. Rather, Salehi and colleagues motivate the need to estimate areal-level parcellations jointly from resting-fMRI, task-fMRI and other modalities (Glasser et al., 2016; Eickhoff et al., 2018a) in order to achieve invariance across brain states. We leave this for future work.

### MS-HBM areal-level parcellations are more homogeneous than other approaches in out-of-sample resting and task-fMRI

Dealing with RSFC matrices at the original voxel or vertex resolution is difficult because of the high-dimensionality. Thus, areal-level brain parcellations have been widely utilized as a dimensionality reduction tool (Eickhoff et al., 2018a), e.g., averaged time course of a parcel is used to represent the entire parcel (Varoquaux and Craddock, 2013; Finn et al., 2015; Rosenberg et al., 2016). For the dimensionality reduction to be valid, vertices within each areal-level parcel should have similar time courses, i.e., high resting-state homogeneity (Gordon et al., 2016; Schaefer et al., 2018). Across two datasets (HCP and MSC), we found that MS-HBM areal-level parcellations exhibited higher resting-state homogeneity than three other approaches, suggesting that rs-fMRI time courses are more similar within MS-HBM parcels (Figures 6 and 7).

Furthermore, if an individual-specific areal-level parcellation accurately captures the functional brain organization of a participant, one might expect task activation to be uniform within parcels, i.e., low task inhomogeneity (Gordon et al., 2017b; Schaefer et al., 2018). We found that MS-HBM parcellations achieved better task inhomogeneity than other approaches in both HCP and MSC datasets (Figure 8 and S11). Given the strong link between task-fMRI and rs-fMRI (Smith et al., 2009; Mennes et al., 2010; Cole et al., 2014; Krienen et al., 2014; Bertolero et al., 2015; Yeo et al., 2015; Tavor et al., 2016), this is perhaps not surprising.

Intriguingly, the improvement in task inhomogeneity varied significantly across task domains (Figures 8 and S11) with the motor task exhibiting the least task inhomogeneity improvement for both HCP and MSC datasets. Given that the motor domain exhibited one of the lowest task inhomogeneity across behavioral domains, there might not be much room for improvement. Furthermore, sensory-motor parcels exhibited low inter-subject variation in terms of location and spatial topography (Figure 4B), so different approaches might perform similarly well. Other possible reasons might include variation in task design and duration.

It is worth pointing out that even though MSC dataset only contained 9 participants, MS-HBM parcellations exhibited better resting homogeneity and task inhomogeneity in every single participant (Figures 6–8, S9–S11). This suggests that MS-HBM parameters estimated from HCP were effective in MSC despite significant acquisition and preprocessing differences.

### MS-HBM works well even with only 10 min of rs-fMRI data

It is well-known that longer scan durations can improve the reliability of RSFC measures (Van Dijk et al., 2010; Xu et al., 2016; Kong et al., 2019). Recent studies have suggested that at least 20-30 min of data is needed to obtain reliable measurements (Laumann et al., 2015; O’Connor et al., 2017; Gordon et al., 2017b). Consistent with previous work, we found that resting-state homogeneity of individual-specific areal-level parcellations continued to improve with more data (Figure 7 and S10). The improvements started to plateau around 40-50 min of data.

Although MS-HBM required multi-session rs-fMRI data for training, the models could be applied to a single rs-fMRI session from a new dataset. More specifically, in the MSC dataset, we showed that MS-HBM areal-level parcellations estimated with only 10 min of rs-fMRI data exhibited better resting-state homogeneity than two other approaches using 150 min of data (Gordon et al., 2017b; Li et al., 2019b).

### RSFC of individual-specific MS-HBM parcellations improves behavioral prediction

A vast body of literature has shown that functional connectivity derived from group-level parcellations can be utilized for behavioral prediction (Hampson et al., 2006; Finn et al., 2015; Smith et al., 2015; Yeo et al., 2015; Rosenberg et al., 2016; He et al., 2019). However, there is a preponderance of evidence that group-level parcellations obscure individual-specific topographic features (Harrison et al., 2015; Laumann et al., 2015; Langs et al., 2016; Braga and Buckner, 2017; Chong et al., 2017; Gordon et al., 2017a, 2017b), which are behaviorally meaningful (Bijsterbosch et al., 2018, 2019; Kong et al., 2019; Seitzman et al., 2019). Recent studies have also suggested that functional connectivity strength derived from individual-specific parcellations might also improve behavioral prediction (Li et al., 2019b; Pervaiz et al., 2019).

We found that MS-HBM parcellations captured individual-specific features that were replicable across sessions (Figures 5, S2–S7). Furthermore, RSFC derived from individual-specific MS-HBM areal-level parcellations achieved better behavioral prediction performance compared with a group-level parcellation (Schaefer et al., 2018) and a recently published individual-specific parcellation approach (Li et al., 2019b). Overall, our results suggest that individual differences in functional connectivity strength of MS-HBM parcels were more behaviorally meaningful than of other parcellation approaches.

It is worth noting that the absolute improvement in prediction performance was modest on average, although some behavioral measures appeared to benefit more than others. For example, when comparing Li2019 and gMS-HBM for behavioral prediction, the prediction accuracy (Pearson’s correlation) of “openness (NEO)” improved from 0.19 to 0.26, while the accuracy (Pearson’s correlation) of “vocabulary (picture matching)” improved from 0.36 to 0.39. Thus, gMS-HBM might be more helpful for predicting certain behavioral measures than others.

Further analysis suggested that task performance measures were on average predicted with higher accuracy than self-reported measures (Figure 11). This differentiation between task performance and self-reported measures was consistent with previous investigations of RSFC-behavior relationships. For example, RSFC has been shown to predict cognition better than personality and mental health (Dubois et al., 2018; Chen et al., 2020). Dynamic functional connectivity is also more strongly associated with cognition and task performance than self-reported measures (Vidaurre et al., 2017; Liégeois et al., 2019). Finally, regressing the global signal has been shown to improve the prediction of task performance measures more than self-reported measures (Li et al., 2019).

### Spatially-localized individual-specific areal-level parcels

Postmortem studies have generally identified cortical areas that are spatially contiguous (Kaas, 1987; Felleman and Van Essen, 1991; Amunts and Zilles, 2015). This has motivated most resting-state areal-level parcellations to estimate spatially contiguous parcels (Shen et al., 2013; Honnorat et al., 2015; Gordon et al., 2016; Chong et al., 2017). One approach to achieve spatially contiguous parcels is to introduce a spatial connectedness term into the optimization objective so that distributed parcels would have large penalty (Honnorat et al., 2015; Schaefer et al., 2018). Another approach is to start with initial spatially contiguous parcels and to iteratively adjust the boundaries to maintain spatial contiguity (Blumensath et al., 2012; Chong et al., 2017; Salehi et al., 2019). Yet another method is to utilize the local-gradient approach, which detects sharp transitions in RSFC profiles, followed by a post-processing procedure (Cohen et al., 2008; Gordon et al., 2016). However, work from Glasser and colleagues suggested that some individual-specific areal-level parcels might comprise multiple spatially close components in some individuals (Glasser et al., 2016).

Given the lack of consensus, we explored three MS-HBM variants in this study. We found that strictly contiguous cMS-HBM parcels achieved the best out-of-sample resting-state homogeneity and task inhomogeneity (Figures 6–8, S8–S10). One possible reason is that cMS-HBM parcellation boundaries were smoother than dMS-HBM and gMS-HBM parcellations. Since fMRI data is spatially smooth, parcellations with smoother boundaries might have an inherent homogeneity advantage, without necessarily being better at capturing true areal boundaries. Another potential artefact of smooth data is the appearance of excessively round parcels that are at odds with histological studies which show that cortical areas express diverse spatial configurations.

Based on our geometric analyses, we found gMS-HBM to be most anatomically plausible among the three parcellations, having both fewer spatially disconnected components than dMS-HBM, and intermediate levels of roundness between dMS-HBM and cMS-HBM. Furthermore, RSFC derived from gMS-HBM parcels achieved the best behavioral prediction performance, albeit not reaching statistical significance (Figures 9, S11; Tables S2–S5). As elaborated in previous studies (Gordon et al., 2016; Schaefer et al., 2018; Kong et al., 2019), assessment of parcellations should integrate and weigh performance across multiple metrics. For the reasons outlined above, we prefer individual-specific gMS-HBM areal-level parcellations among the three MS-HBM variants.

Overall, our findings suggest that the brain’s large-scale organization might potentially comprise certain functional regions that are spatially disconnected. Neuronal migration, guided by cell-to-cell interactions and gradients of diffusible cues, plays an important role in establishing the brain’s complex cytoarchitectonic organization during embryogenesis (Silva et al., 2019). Spatially disconnected parcels might reflect functionally analogous neuronal populations from the same cellular lineage that separate due to natural variation in migration patterns in early development.

That said, we are aware that one cannot establish with certainty the existence of spatially disconnected cortical areas based on resting-fMRI data alone. It is possible that disconnected components of a non-contiguous parcel are inseparable by resting-fMRI measurements, but are separable by other neural properties, such as microstructure or task activations. Given that fMRI is an indirect measurement of neuronal signals, the functional coupling among disconnected components could also be driven by non-neural mechanisms (e.g., vasculature).

Nevertheless, our individual-level areal parcellation provides an explicit model that can be further validated using prospectively acquired rs-fMRI paired with other approaches, e.g., post-mortem histological analyses (Xu et al., 2018; Hayashi et al., 2020) or with spatially-targeted intracranial recording (Wang et al., 2015; Fox et al., 2018).

## Conclusions

We proposed a multi-session hierarchical Bayesian model (MS-HBM) that accounted for both inter-subject and intra-subject functional connectivity variability when estimating individual-specific areal-level parcellations. Three MS-HBM variants with different spatial localization priors were explored. Using 10 min of rs-fMRI data, individual-specific MS-HBM areal-level parcellations generalized better to out-of-sample rs-fMRI data from the same participants than a group-level parcellation approach and two prominent individual-specific areal-level parcellation approaches using 150 min of rs-fMRI data. Furthermore, RSFC derived from MS-HBM parcellations exhibited better behavioral prediction performance than alternative parcellation approaches.

## Acknowledgment

We like to thank Danilo Bzdok, John Murray, Michael Breakspear, Peter Fox, and Jack Lancaster for insightful comments and feedback on this work. This work was supported by the Singapore National Research Foundation (NRF) Fellowship (Class of 2017), the Singapore Ministry of Defense (Project CURATE) and the National University of Singapore Yong Loo Lin School of Medicine (NUHSRO/2020/124/TMR/LOA). Any opinions, findings and conclusions or recommendations expressed in this material are those of the authors and do not reflect the views of the Singapore Ministry of Defense and National Research Foundation, Singapore. Computational work for this article was partially performed on resources of the National Supercomputing Centre, Singapore (https://www.nscc.sg). Our research also utilized resources provided by the Center for Functional Neuroimaging Technologies, P41EB015896 and instruments supported by 1S10RR023401, 1S10RR019307, and 1S10RR023043 from the Athinoula A. Martinos Center for Biomedical Imaging at the Massachusetts General Hospital. Data were in part provided by the Human Connectome Project, WU-Minn Consortium (Principal Investigators: David Van Essen and Kamil Ugurbil; 1U54MH091657) funded by the 16 NIH Institutes and Centers that support the NIH Blueprint for Neuroscience Research; and by the McDonnell Center for Systems Neuroscience at Washington University.

## Supplemental Material

This supplemental material is divided into *Supplemental Methods* and *Supplemental Results* to complement the Methods and Results sections in the main text, respectively.

## Supplementary Methods

This section provides additional mathematical and implementation details of the multi-session hierarchical Bayesian model (MS-HBM). Section S1 provides mathematical details about the generative model. Section S2 describes the algorithms for estimating group-level priors and deriving the individual-specific parcellations and how “free” parameters of the model are set. Section S3 describes the matching algorithms for comparing MS-HBM parcellations with another parcellation approach. Section S4 describes the kernel ridge regression model for behavioral prediction.

### S1. Mathematical model

In this section, we describe our model for individual-specific areal-level parcellation of the cerebral cortex. We assume a common surface coordinate system, where the cerebral cortex is represented by left and right hemisphere spherical meshes such as fs_LR32k surface meshes. Each mesh consists of a collection of vertices and edges connecting neighboring vertices into triangles (https://en.wikipedia.org/wiki/Triangle_mesh).

Let *N* denote the total number of vertices, *T* denote the number of resting-state fMRI (rs-fMRI) sessions, *S* denote the number of subjects, *L* denote the number of parcels, and 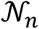 denote the neighboring vertices of vertex *n* (as defined by the cortical mesh). For each subject s and session *t*, there is a preprocessed rs-fMRI time course associated with each vertex *n*. For each subject *s*, there is an unknown parcellation label 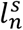 at vertex *n*. Note that the parcellation label is assumed to be the same across sessions (hence there is no index on the session). In this work, we use 1:*S* to denote a set of subjects {1, 2,…, *S*}, 1: *T* to denote a set of sessions {1, 2,…, *T*}, 1: *N* to denote a set of vertices {1, 2,…, *N*}, 1: *L* to denote a set of parcellation labels {1, 2,…, *L*}.

For each subject s at a particular session t, we computed the functional connectivity profile of each vertex (of the cortical mesh) by correlating the vertex’s fMRI time course with the time courses of uniformly distributed cortical regions of interests (ROIs). For the HCP and MSC datasets, the preprocessed data were in fs_LR32k surface space. The ROIs consisted of 1483 vertices spaced approximately uniformly distributed across the two hemispheres. Each vertex’s connectivity profile was binarized (see Methods in main manuscript) and normalized to unit length. Let 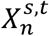 denote the binarized, normalized functional connectivity profile of subject *s* at vertex *n* during session *t*. Let *D* denote the total number of ROIs and hence the length of 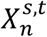. We denote the connectivity profiles from all sessions of all subjects at all cortical vertices as 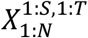.

Figure 1 (main text) illustrates the schematic of the areal-level multi-session hierarchical Bayesian model (MS-HBM). Following previous work (Yeo et al., 2011; Kong et al., 2019), the functional connectivity profile 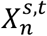 of subject s from a session t at vertex n is assumed to be generated from a von Mises-Fisher distribution,

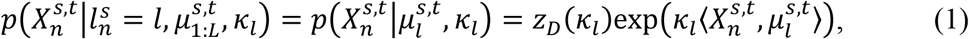

where 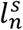 is the parcellation label at vertex *n* of subject *s*, and 〈,〉 denote inner product. 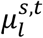 and *κ_l_* are the mean direction and concentration parameter of the von Mises-Fisher distribution for parcel label *l* of subject s during session *t*. 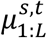 are the mean directions for networks 1 to *L*. We can think of 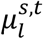 as the mean connectivity profile of network label *l* normalized to unit length. If functional connectivity profile 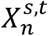 is similar to mean connectivity profile 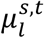 (i.e., 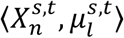 is big), then vertex *n* is more likely to be assigned to parcel *l*. The concentration parameter *κ_l_* controls the variability of the functional connectivity profiles within parcel *l*. A higher *κ_l_* results in a lower dispersion (i.e., lower variance), which means that vertices belonging to the same parcel are more likely to possess functional connectivity profiles that are close to the mean connectivity profile of the parcel. *κ_l_* is assumed to be the same for all parcels, subjects and sessions. Thus, we will use *κ* to indicate *κ_l_* henceforth. Finally, *z_D_*(*κ*) is a normalization constant to ensure a valid probability distribution (Banerjee et al., 2005):

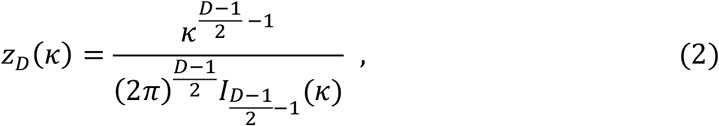

where 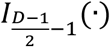 is the modified Bessel function of the first kind with order 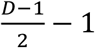.

Similar to our previous work (Kong et al., 2019), we modeled both inter- and intra-subject variability. To model intra-subject functional connectivity variability, we assume a conjugate prior on the subject-specific and session-specific mean connectivity profiles 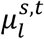, which turns out to also be a von Mises-Fisher distribution:

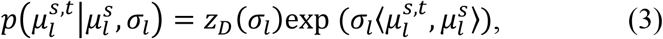

where 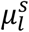 and *σ_l_* are the mean direction and concentration parameter of the von Mises-Fisher distribution for parcel label *l* of subject *s*. We can think of 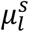 as the individual-specific functional connectivity profile of parcel *l* of subject *s*. The concentration parameter *σ_l_* controls how much the session-specific mean direction 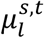 of subject *s* during session *t* can deviate from the subject-specific mean direction 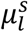. A higher *σ_l_* would imply lower intra-subject functional connectivity variability across sessions. *σ_l_* is network-specific but is assumed to be the same for all subjects.

To model inter-subject functional connectivity variability, we assume a conjugate prior on the subject-specific mean connectivity profiles 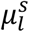, which is again a von Mises-Fisher distribution whose mean direction corresponded to the group-level mean direction *μ^g^*:

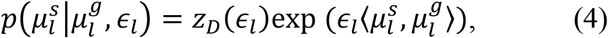

where *μ^g^* and *ϵ_l_* are the mean direction and concentration parameter of the von Mises-Fisher distribution for parcel label *l*. We can think of 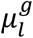 as the group-level functional connectivity profile of parcel *l*. The concentration parameter *ϵ_l_* controls how much the individual-specific connectivity profile 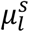 can deviate from the group-level connectivity profile 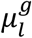. A higher *ϵ_l_* would imply lower inter-subject functional connectivity variability across subjects.

Because the functional connectivity profiles of individual subjects are generally very noisy, we impose a MRF prior on the hidden parcellation labels 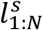

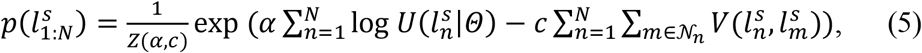

where *Z*(*α, c*) is a normalization term (partition function) to ensure 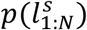 is a valid probability distribution. log 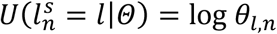 is a singleton potential encouraging certain vertices to be associated with certain labels. 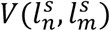 is a pairwise potential (Potts model) encouraging neighboring vertices to have the same parcellation labels:

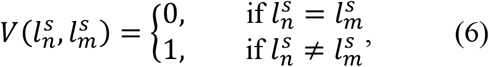

The parameters *α* and *c* are tunable parameters greater than zero, and control the tradeoffs between the various terms in the generative model. Assuming that *α* = 1 and *c* = 0, then *θ_l,n_* can be interpreted as the probability of label *l* occurring at vertex *n* of subject *s*.

However, many parcels will be spatially distributed due to strong long-range RSFC. Requiring parcels in a MRF framework to be spatially connected (with minimal other assumptions) is non-trivial (Nowozin and Lampert 2010; Honnorat et al. 2015). Here, spatial localization prior is imposed by Φ:

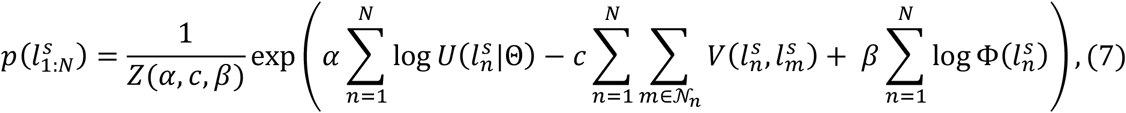

where 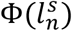 is a singleton potential encoding the spatial localization constraint. The parameters *β* is a tunable parameter greater than zero. We consider three different spatial localization priors which will be explained in S1.1, S1.2, and S1.3.

In addition, it is well-known that early sensory and late motor architectonic areas (areas 3 and 4) are on either side of the central sulcus. Given low inter-subject variability of these areas (Fischl et al., 2008), we do not expect this property to be violated in individuals. Therefore, we additionally constrained the Schaefer2018 group-level parcels on either side of the central sulcus not to extend across the central sulcus in the individual-specific areal-level parcellations. Removal of this constraint resulted in parcels extending across the central sulcus, while not yielding any quantitative improvements, e.g., in behavioral prediction accuracies. Consequently, we decided to keep this constraint because the resulting parcellations were biologically more plausible.

#### S1.1 Distributed MS-HBM (dMS-HBM)

We considered a spatial localization prior that constrained each individual-specific parcel to be within 30mm of the group-level Schaefer2018 parcel boundaries (similar to Glasser et al., 2016). The spatial localization prior Φ = Φ^*d*^ is therefore defined as a *N*×*L* binary mask, where 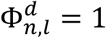 if the shortest geodesic distance between vertex *n* and group-level parcel *l* is less than (or equal to) 30 mm, else 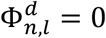. Thus, when 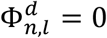, then it is impossible for the vertex *n* to be assigned to parcel *l* (Eq. (7)).

#### S1.2 Contiguous MS-HBM (cMS-HBM)

We considered a spatial localization prior that encouraged brain locations with similar 3-dimensional spherical coordinates to be grouped into the same parcel (similar to Schaefer et al., 2018). Here, we denote the 3 × 1 spherical coordinates of vertex *n* as *Y_n_*. The spatial localization prior 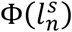 is then defined as

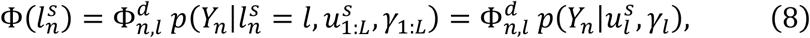

where 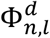 is the spatial localization constraint from dMS-HBM (S1.1) and 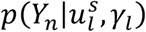 follows a von Mises-Fisher distribution:

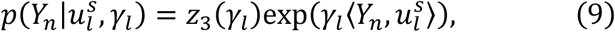

where 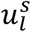 and *γ_l_* are the mean direction and concentration parameter of the von Mises-Fisher distribution for parcel label *l* of subject *s*. We can think of 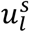 as the mean spatial coordinates of parcel *l* normalized to unit norm (i.e., sphere). Therefore, if vertex *n* is spatially close to the mean spatial location of parcel *l* (i.e., 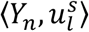 is big), then vertex *n* is more likely to be assigned to parcel *l*. Consequently, for large enough values of *γ_l_*, the parcels will be spatially connected. In the optimization procedure, we start with a smaller value for *γ_l_* and then iteratively increase *γ_l_* for parcel *l* to ensure spatial contiguity.

#### S1.3 Gradient-infused MS-HBM (gMS-HBM)

##### S1.3.1 Gradient-infused spatial localization prior

A well-known areal-level parcellation approach is the local gradient approach, which detects local abrupt changes (i.e., gradients) in RSFC across the cortex (Cohen et al., 2008). Our previous study (Schaefer et al., 2018) has suggested the utility of combining local gradient (Cohen et al., 2008; Gordon et al., 2016) and global clustering (Yeo et al., 2011) approaches for estimating areal-level parcellations. Therefore, in the case of gradient-infused MS-HBM (gMS-HBM), the spatial contiguity prior in cMS-HBM is complemented with a prior based on RSFC gradient maps, which encourages brain locations with gentle changes in functional connectivity to be grouped into the same parcel.

For each subject *s*, the RSFC gradient map *Gmap^s^* is an *N* × 1 vector, where *N* is the number of vertices. High *Gmap^s^* values indicate abrupt change in RSFC; low *Gmap^s^* values indicate gentle change in RSFC. We define a graph where the vertices and edges correspond to the mesh structure of the surface mesh. For a particular subject *s*, the distance between two adjacent vertices *a* and *b* is defined as 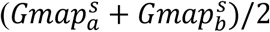. Based on this graph, we constructed an *N* × *N* gradient distance matrix *Gmat^s^*, where each entry represents the shortest geodesic distance between each pair of vertices. If there is an abrupt change in RSFC (high *Gmap^s^*) somewhere along all paths linking two vertices *i* and *m*, then the shortest geodesic distance between *i* and *m* will be large, even though *i* and *m* are spatially close. By contrast, if there is a path linking vertex *i* and vertex *m*, such that RSFC changes are gentle (low *Gmap^s^*) along the entire path, then the shortest geodesic distance between *i* and *m* will be relatively small, except if vertices *i* and *m* are far apart.

Ideally, we would like to incorporate *Gmat^s^* into our model: two vertices with high *Gmat^s^* should be encouraged to be in different parcels. This will generally encourage spatially contiguous parcels since two vertices spatially far apart will generally have higher *Gmat^s^* and thus will be encouraged to be in separate parcels. Because gradient-based parcellations tend to have irregular-shaped parcels (Gordon et al., 2016), we will also be encouraging spatial contiguity without relatively round parcels like in cMS-HBM. However, incorporating *Gmat^s^* into our model is not easy because *Gmat^s^* is an *N* × *N* distance matrix, which is not easily incorporated into our mixture-type model. Furthermore, *N* is also quite large (e.g., more than 30K), so dealing with an *N* × *N* matrix would slow down our algorithm.

Therefore, we applied the diffusion embedding algorithm(Coifman et al., 2005) to the *N × N* geodesic gradient distance matrix *Gmat^s^*, thus embedding the *N* vertices into a *K* dimensional space. Roughly speaking, the Euclidean distance between two vertices (based on their *K* diffusion coordinates) should be similarly to the geodesic distance. The larger *K* would lead to a better approximation, but with more computation costs. We found that *K* = 100 leads to a sufficiently good approximation. More specifically, the correlation between *Gmat^s^* and the Euclidean distance in the *K* dimensional space was 0.84 when averaged across 40 random HCP subjects. In this way, for a particular subject *s*, the information of the geodesic gradient distance matrix *Gmat^s^* is embedded as a 1 × 100 diffusion coordinates for each vertex *n*, which we denote as 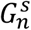. If there is an abrupt change in RSFC (high *Gmap^s^*) somewhere along all paths linking two vertices *i* and *m*, then the Euclidean distance between their diffusion coordinates 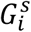 and 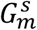 will be large. By contrast, if there is a path linking vertex *i* and vertex *m*, such that RSFC changes are gentle (low *Gmap^s^*) along the entire path, then the Euclidean distance between their diffusion coordinates 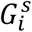 and 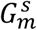 will be relatively small, except if vertices *i* and *m* are far apart. The spatial localization prior 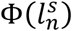 is thus defined as

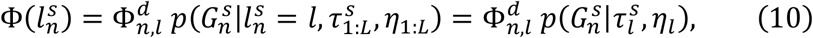

where 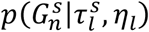 follows a Gaussian distribution:

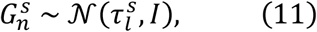

where 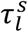 is the mean and *I* is the identity matrix. We can think of 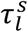 as the mean diffusion coordinates of parcel *l*. Therefore, if vertex *n* is spatially close to the mean diffusion coordinates of parcel *l* (i.e., 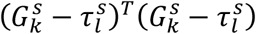 is small), then vertex *n* is more likely to be assigned to parcel *l*.

We note that the goal of the diffusion embedding was not to compute principal gradients (Margulies et al., 2016), but simply for dimensionality reduction to reduce computational costs and memory requirements when training and applying gMS-HBM. In addition, the computation of the local gradient (Gordon et al., 2016), the computation of the gradient distance matrix *Gmat^s^* and the diffusion embedding of *Gmat^s^* also required significant computation costs and memory. Thus, in the following sections (Supplementary Methods S1.3.2 and Supplementary Methods S1.3.3), we discussed how to speed up these computations and reduce memory requirements.

##### S1.3.2 Reducing computational costs of gradient maps

We mentioned in Section S1.3.1 that the spatial localization prior in gMS-HBM was constructed based on the RSFC gradient maps. Figure SM1 (left panel) illustrates the original approach for generating gradient map for a particular subject (Laumann et al., 2015; Gordon et al., 2016). Given the resting-state fMRI data of a subject, the *N* × *N* RSFC matrix was first computed. Each row and each column of the RSFC matrix were then correlated to obtain an *N* × *N* RSFC similarity matrix, where the *n*-th column represented the spatial correlation of functional connectivity patterns between vertex *n* and all other vertices. By computing the first order spatial gradient for each column of the RSFC similarity matrix, we obtained the gradients of the RSFC similarity matrix, where high gradient values indicated rapid changes in RSFC, and low gradient values indicated gentle changes in RSFC. After that, the watershed algorithm^1^ was applied to each column of the gradient matrix to generate one binarized gradient map for each vertex. The *N* binarized gradient maps were then averaged to obtain a single gradient map, where high values indicate high probability of a rapid change in RSFC patterns occurring at that location. Because this original version involved *N* × *N* matrices, it was computationally very expensive to compute.

**Figure SM1.**
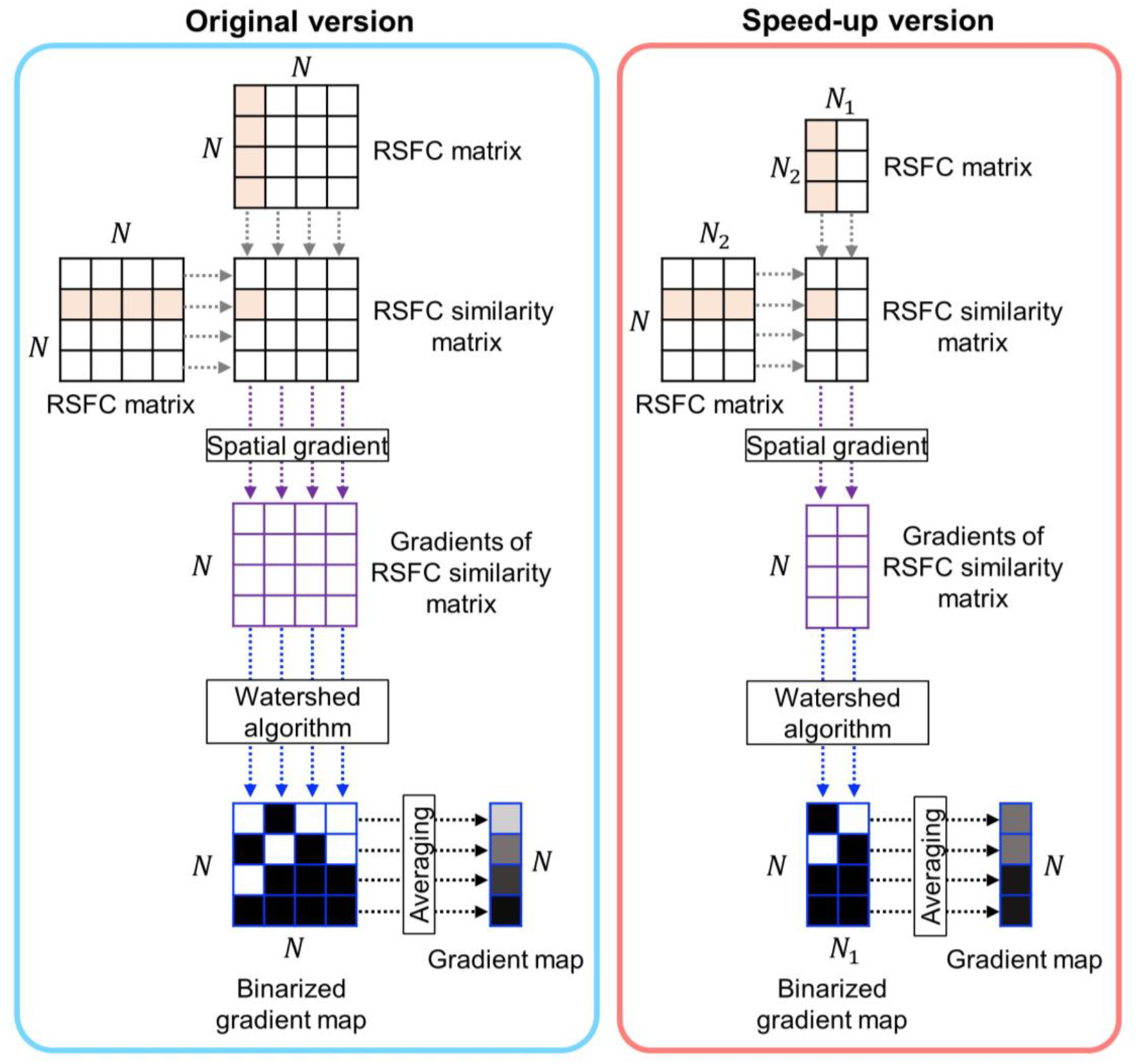
Illustration of the procedure to generate a RSFC gradient map. Left panel shows the original version, right panel shows our speed-up version. In the case of fs_LR32k, note that *N* = 59412. In the case of fsaverage6, note that *N* = 74947.

Therefore, we implemented a faster and less memory-intensive version by subsampling the functional connectivity matrices. Figure SM1 right panel illustrates the speed-up version for generating the gradient map for each participant. Instead of computing the *N* × *N* RSFC matrix to generate the RSFC similarity matrix, we randomly selected *N*_1_ and *N*_2_ vertices from *N* vertices to compute the subsampled *N* × *N*_2_ and *N*_2_ × *N*_1_ RSFC matrices. We correlated each row of the *N* × *N*_2_ RSFC matrix and each column of the *N*_2_ × *N*_1_ RSFC matrix to obtain the subsampled *N* × *N*_1_ RSFC similarity matrix. We then computed the spatial gradients of the subsampled RSFC similarity matrix and utilized the watershed algorithm to obtain *N*_1_ binarized gradient maps. The *N*_1_ binarized gradient maps were then averaged to obtain a single gradient map. Furthermore, to reduce the memory usage, we split the *N* × *N*_1_ RSFC similarity matrix into multiple small blocks (without doing any subsampling), where each block of the RSFC similarity matrix was generated sequentially.

In the case of fs_LR32k, on our machine, the original algorithm (working with the full 59412 × 59412 RSFC matrix) required 40GB of RAM and 4 hours using 1 CPU core to compute the gradient map for a single rs-fMRI run. On the same machine, the speed-up version (*N*_1_, = 297 and *N*_2_ = 5941) only required 3GB of RAM and 15 minutes with 1 core. The resulting gradient maps were highly similar to the original gradient maps (r = 0.97 averaged across 40 random participants). In the case of fsaverage6, on our machine, the original algorithm (working with the full 74947 × 74947 RSFC matrix) required 60GB of RAM and 6 hours using 1 CPU core to compute the gradient map for a single rs-fMRI run. On the same machine, the speed-up version (*N*^1^ = 375 and *N*_2_ = 7495) only required 4GB of RAM and 20 minutes with 1 core. The resulting gradient maps were highly similar to the original gradient maps (r = 0.97 averaged across 40 random participants).

##### S1.3.3 Reducing computational costs of computing the diffusion embedding matrix

In Section S1.3.1, we mentioned that the RSFC gradient map was utilized to generate an *N* × *N* geodesic gradient distance matrices. We then applied the diffusion embedding algorithm (Margulies et al., 2016) to reduce the dimensionality to 100 diffusion coordinates. The resulting *N* × 100 diffusion embedding matrix was utilized for the spatial localization prior in Eq. (10). However, this approach involved *N* × *N* geodesic gradient distance matrices, so it was also computationally expensive.

To overcome this issue, we downsampled the *N* × 1 RSFC gradient map from S1.3.2 to a lower resolution mesh with Af vertices. The resulting *N*_3_ × 1 downsampled gradient map was then used to generate *N*_3_ × *N*_3_ geodesic gradient distance matrices. The diffusion embedding algorithm was performed on the *N*_3_ × *N*_3_ gradient distance matrix to obtain an *N*_3_ × 100 diffusion embedding matrix. After that, we upsampled the *N*_3_ × 100 diffusion embedding matrix back to the original space to generate an *N* × 100 diffusion embedding matrix.

In the case of fs_LR32k, on our machine, the original diffusion embedding algorithm (working with the full 59412 × 59412 geodesic gradient distance matrices) required 6GB of RAM and 2 hours using 1 core to convert the *N* × 1 RSFC gradient map (Section S1.3.2) to an 59412 × 100 diffusion embedding matrix. On the same machine, the sped-up diffusion embedding algorithm (*N*_3_ = 20484) only required 3 GB of RAM and 10 minutes with 1 core. The resulting diffusion embedding matrices were highly similar to the original diffusion embedding matrices computed using the original gradient algorithm and original diffusion embedding algorithm (r = 0.98 averaged across 40 random participants). In the case of fsaverage6, on our machine, the original diffusion embedding algorithm (working with the full 74947 × 74947 geodesic gradient distance matrices) required 8GB of RAM and 2.5 hours using 1 core to convert the 74947 × 1 RSFC gradient map (Section S1.3.2) to an 74947 × 100 diffusion embedding matrix. On the same machine, the sped-up diffusion embedding algorithm (*N*_3_ = 25924) only required 3 GB of RAM and 10 minutes with 1 core. The resulting diffusion embedding matrices were highly similar to the original diffusion embedding matrices computed using the original gradient algorithm and original diffusion embedding algorithm (r = 0.97 averaged across 40 random participants).

### S2. Model estimation

In this section, we describe how model parameters are estimated from a training set and a validation set (Section S2.1), and how the parameters can be used to parcellate a new subject (Section S2.2). Throughout the entire section, we assume that the number of parcels *L* = 400 without loss of generality.

#### S2.1 Learning model parameters

Our goal is to estimate the model parameters 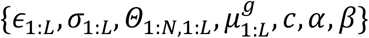 from a training set and a validation set of binarized and normalized functional connectivity profiles, which can then be utilized for estimating individual-specific parcellations in unseen data of new subjects (Section S2.2). As a reminder, *ϵ*_1:*L*_ is a group prior representing inter-subject functional connectivity variability, *σ*_1:*L*_ is a group prior corresponding to intra-subject functional connectivity variability, *Θ*_1:*N*,1:*L*_ is a group prior representing inter-subject spatial variability and reflects the probability of a parcel occurring at particular spatial location, and 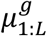 is the group-level connectivity profile for each parcel. The parameters *α, c* and *β* tradeoff between various terms in the generative model. Because the partition function *Z*(*α, c, β*) (Eq. (7)) is NP-hard to compute, for computational efficiency, we first assume *α* = 1, *c* = 0 and for a particular value of *β* in order to estimate 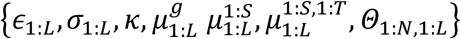 and spatial localization prior parameters 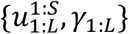 (for cMS-HBM) or 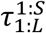 (for gMS-HBM) from the training dataset. Under this scenario, *Z*(*α, c, β*) is a constant. The tunable parameters *α, c* and *β* are then estimated in the validation set using a grid search.

##### S2.1.1 Estimating 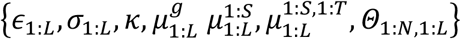 and spatial localization prior parameters from training set

###### S2.1.1.1 dMS-HBM

Given observed binarized, normalized functional connectivity profiles 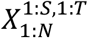 from the training set and spatial localization prior 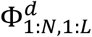, we seek to estimate 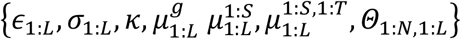 using Expectation-Maximization (EM). As previously explained, we assume *α* = 1, *c* = 0 and a particular value of *β*. Since 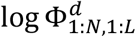 is either –∞ or 0 (i.e., hard constraint), we can simply assume *β* = 1.

Let 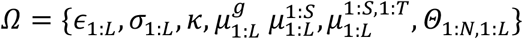. We consider the following maximum-a-posterior (MAP) estimation problem:

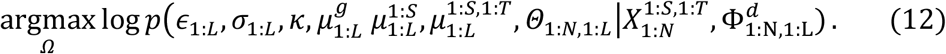

Assuming a uniform (improper) prior on {*Θ*_1:*N*, 1:*L*_, *κ, σ*_1:*L*_, *ϵ*_1:*L*_}, the MAP problem can be written as

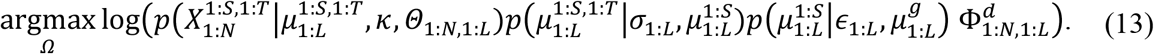

We then introduce the parcellation labels 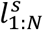 for each subject s as latent variables, and use Jensen’s inequality to define a lower bound 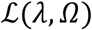, where 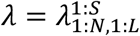 are the parameters of the *q* functions 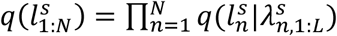:

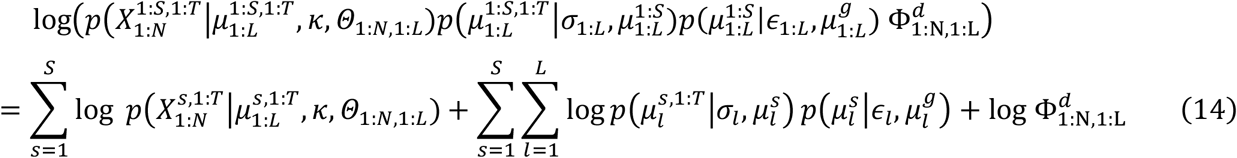

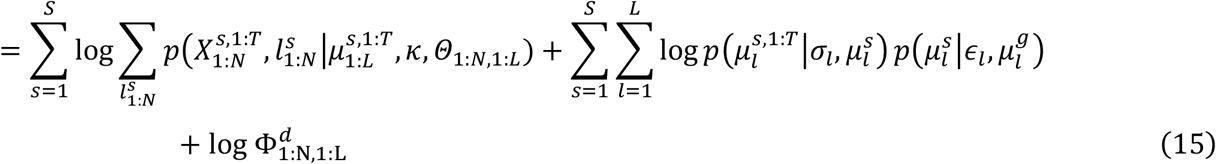

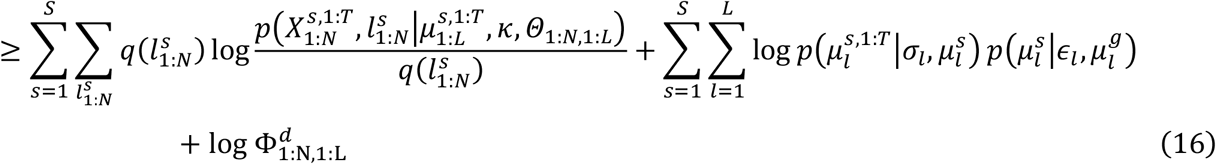

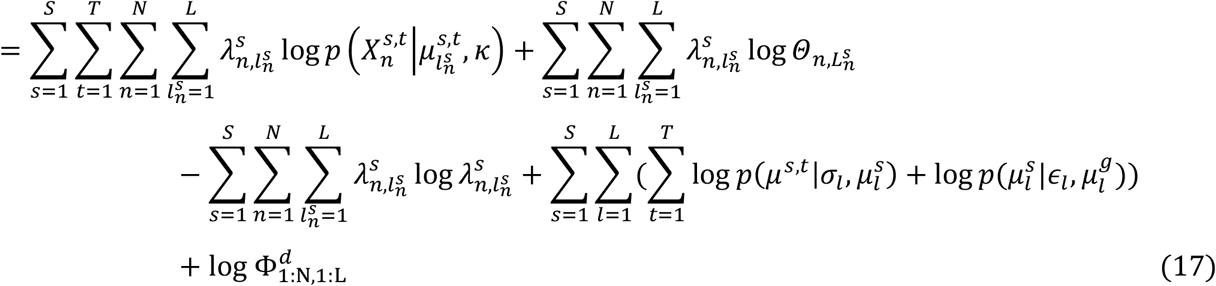

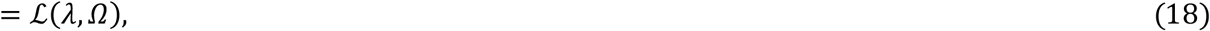

where equality is achieved when 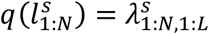 are the posterior probability of the individual-specific parcellation of subject s given the parameters *Ω*. Therefore, instead of maximizing the original MAP problem (Eq. (12)), we instead maximize the lower bound:

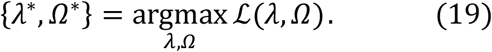

In the E-step, we fix 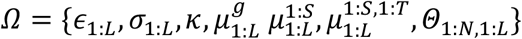, and estimate *λ*:

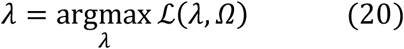

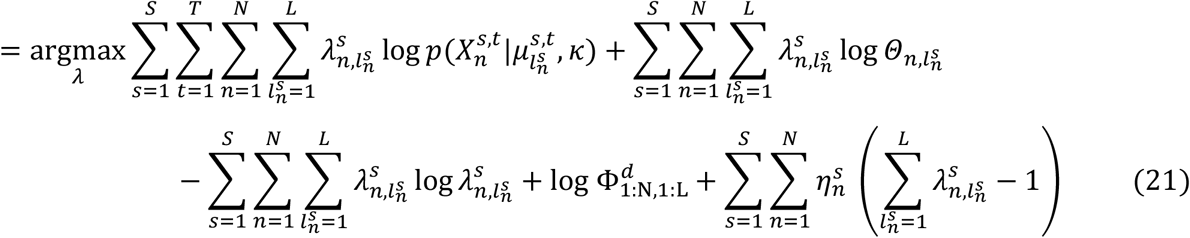

where 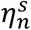 are the Lagrange multipliers enforcing the constraint 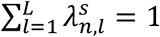. Optimizing Eq. (21) by differentiating with respect to 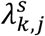 (where *k* and *j* are dummy variables indexing location and parcel label respectively) and setting to 0, we get:

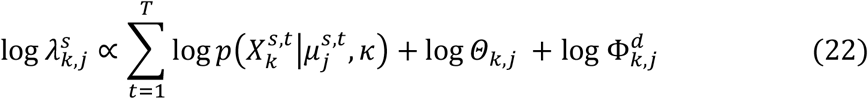

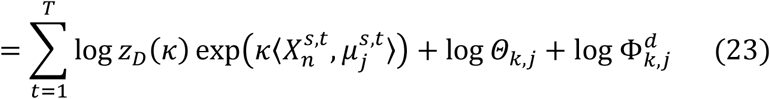

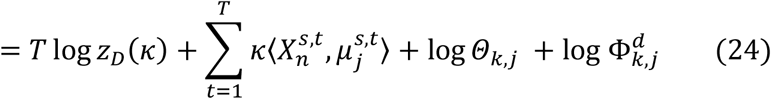

In the M-step, we fix *λ* and estimate *Ω*.

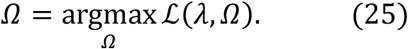

By using the constraints that 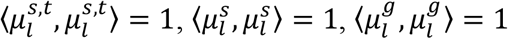, *κ* > 0, *σ_l_* > 0, *ϵ_l_* > 0, and differentiating 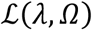 with respect to 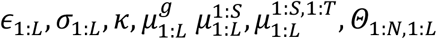, and setting the derivatives to zero, we get the following update equations:

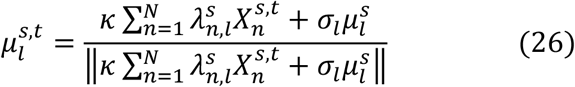

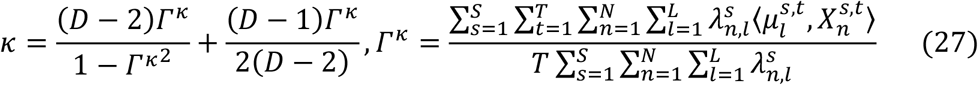

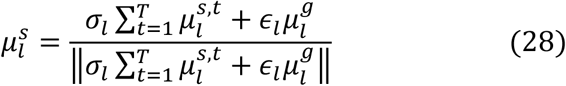

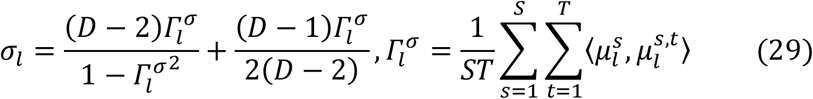

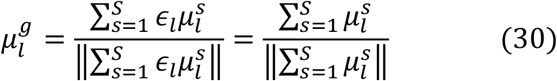

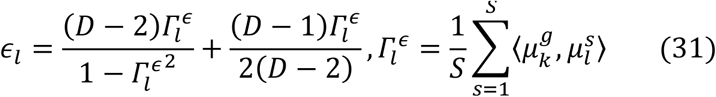

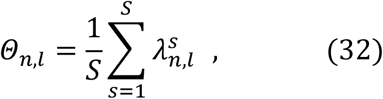

where *D* is the length of 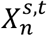 (i.e., number of ROIs in each functional connectivity profile), *S* is the number of subjects, *T* is the number of sessions, and ║·║ corresponds to the *l*_2_-norm. Therefore, the estimate of the functional connectivity profile 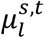 (Eq. (26)) of parcel *l* of subject *s* during session *t* is the weighted sum of the average time course of vertices constituting parcel *l* of subject s during session 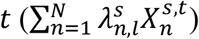 and the subject-specific mean direction 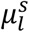, with weights *κ* and *σ_l_* for each term, normalized to be unit norm. If *σ_l_* is much greater than *κ*, then 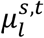 is more likely to be dominated by subject-specific mean direction 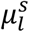, which means that the functional connectivity profile of parcel *l* is highly stable across sessions. Similarly, the estimate of the functional connectivity profile 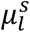 (Eq. (28)) of parcel *l* of subject s is the weighted sum of the average session-specific mean directions across all sessions for parcel *l* of subject 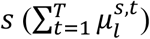 and the group-level mean direction 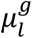, with weights *σ_l_* and *ϵ_l_* for each term, normalized to be unit norm. If *ϵ_l_* is much greater than *σ_l_*, then 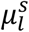 is more likely to be dominated by group-level mean direction 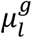, which means that the functional connectivity profile of parcel *l* is highly stable between subjects. Finally, the estimate of the group-level functional connectivity profile 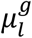 (Eq. (30)) of parcel *l* is the sum of the subject-specific mean directions across all subjects for parcel 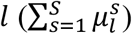, normalized to be unit norm. The estimate of *Θ_n,l_* (Eq. (32)) is the posterior probability of parcel *I* being assigned to vertex *n*, averaged across all the subjects.

Given the training set, the algorithm first utilizes the 400-region Schaefer2018 group-level parcellation (Schaefer et al., 2018) to initialize the EM algorithm. The EM algorithm iterates E-step (Eq. (24)) and M-step (Eqs. (26–32)) till convergence. We note that the update equations (Eqs. (26–32)) in the M-step are dependent on each other. Therefore, within the M-step, the update equations (Eqs. (26–32) are iterated till convergence.

###### S2.1.1.2 cMS-HBM

Similar to dMS-HBM in S2.1.1.1, given observed binarized, normalized functional connectivity profiles 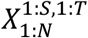 from the training set and the spherical coordinate *Y*_1:*N*_, we seek to estimate 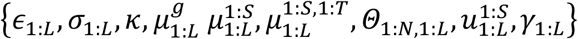 using Expectation-Maximization (EM). As previously explained, we assume *α* = 1, *c* = 0 and a particular value of *β*. We will have:

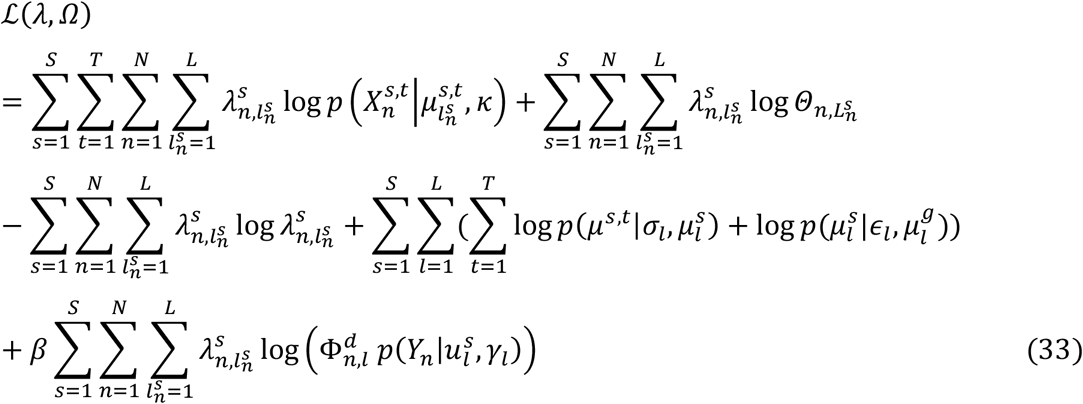

By applying EM algorithm explained in S2.1.1.1, we will have E-step:

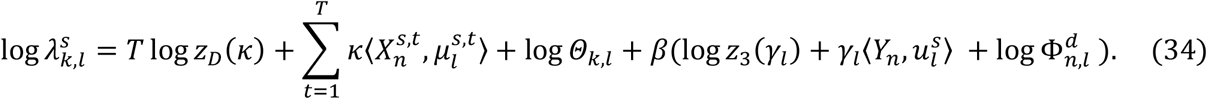

In M-step, 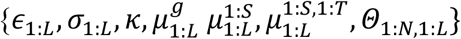 can be updated using the same formulas as Eqs. (26–32). The spatial localization prior parameters 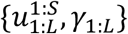 can be updated by the following equations:

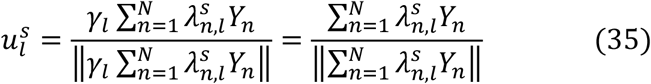

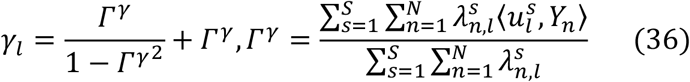

###### S2.1.1.3 gMS-HBM

Similar to dMS-HBM in S2.1.1.1, given observed binarized, normalized functional connectivity profiles 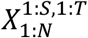 from the training set and gradient matrices 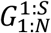, we seek to estimate 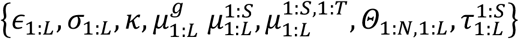 using Expectation-Maximization (EM). As previously explained, we assume *α* = 1, *c* = 0 and a particular value of *β*. We will have:

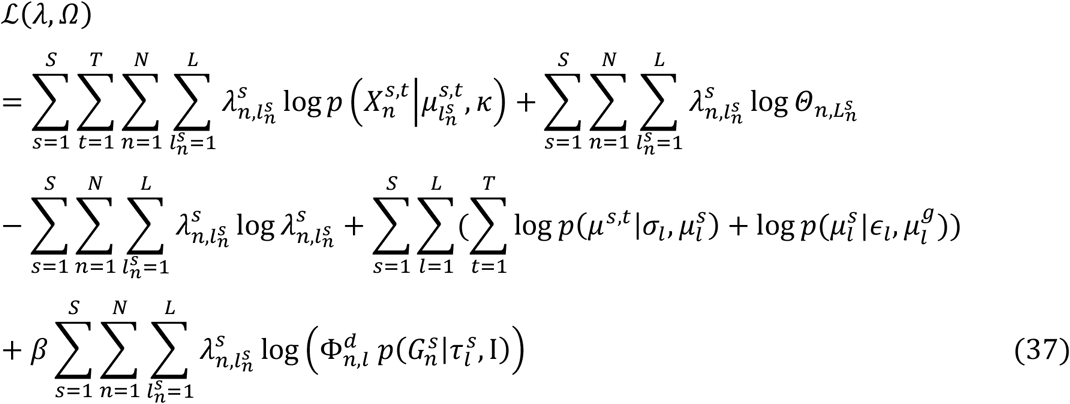

By applying EM algorithm explained in S2.1.1.1, we will have E-step:

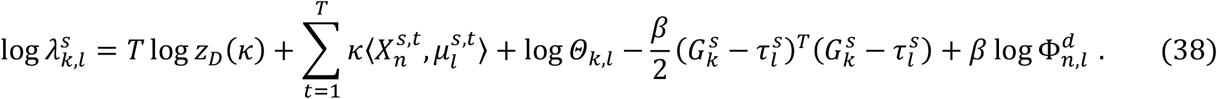

In M-step, 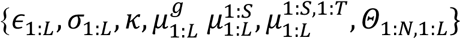 can be updated using the same formulas as Eqs. (26–32). The spatial localization prior parameters 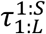 can be updated by the following equations:

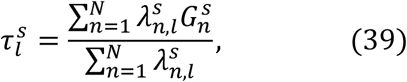

##### S2.1.2 Estimating tunable parameters c, α and β

###### S2.1.2.1 dMS-HBM

In the previous subsection (Section S2.1.1), the training set was used to estimate 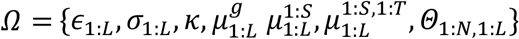, assuming *α* = 1, *c* = 0 and *β* = 1. To tune the parameters *c* and *α*, we assume access to a validation set. Recall that each subject in the validation set has multiple rs-fMRI sessions. We consider *c* ∈ {0.1,1,5,10,20,30,40,50,60} and *α* ∈ {0.1,1, 5,10,20,30,40,50,60}. For a given pair of (*c, α*), and given 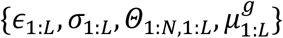 estimated from the training set, we estimate for each subject in the validation set, the individual-specific parcellation based on a subset of rs-fMRI sessions (see Section S2.2 for algorithm). Restingstate homogeneity (main text) is then computed in the remaining rs-fMRI sessions of the validation subjects. The pair of (*c, α*) with the highest homogeneity in the unseen rs-fMRI sessions of the validation subjects is then utilized for parcellating new subjects. For the HCP dataset, the optimal pair of parameters was *c* = 50 and *α* = 20.

###### S2.1.2.2 cMS-HBM and gMS-HBM

Similar to dMS-HBM in S2.1.2.1, the training set was used to estimate 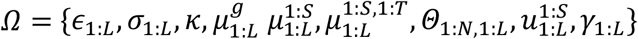 for cMS-HBM and 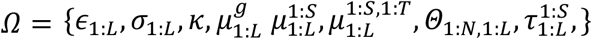 for gMS-HBM with *β* ∈ {10, 30, 50,100,150}, assuming *α* = 1, *c* = 0. To tune the parameters *c, α* and *β*, we assume access to a validation set. We consider, *c* ∈ {0.1,1, 5,10,20, 30,40, 50,60} and *α* ∈ {0.1,1,5,10,20, 30,40, 50,60}. For a given triplet of (*c, α, β*), and given 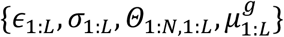 estimated from the training set with *β*, we estimate for each subject in the validation set, the individual-specific parcellation based on a subset of rs-fMRI sessions (see Section S2.2 for algorithm). Resting-state homogeneity (main text) is then computed in the remaining rs-fMRI sessions of the validation subjects. The triplet (*c, α, β*) with the highest homogeneity in the unseen rs-fMRI sessions of the validation subjects is then utilized for parcellating new subjects. For cMS-HBM, the optimal triplet of parameters was *c* = 1, *α* = 1 and *β* = 100. For gMS-HBM, the optimal triplet of parameters was *c* = 30, *α* = 30 and *β* = 100.

Throughout the paper (main text), the reported quality (Figures 6–11) of the individual-specific parcellations was evaluated using subjects not used to tune the parameters. For example, in the case of the HCP data (Figures 6–11), model parameters were estimated the HCP training and validation sets, while the reported quality of the individual-specific parcellations was evaluated using the HCP test set. In the case of the MSC subjects (Figures 6–8), the model parameters were estimated from the HCP training and validation sets.

#### 52.2 Individual-levelparcellation estimation

##### S2.2.1 dMS-HBM

Using parameters 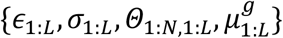 estimated from the training set (Section S2.1.1), and for a particular triplet (*c, α, β*), we can estimate the individual-specific parcellation 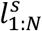 of a new subject *s* with *T* sessions by employing the variational Bayes expectation maximization (VBEM) algorithm.

Let 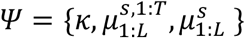. We consider the following maximum-a-posterior (MAP) estimation problem:

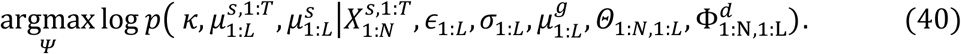

Assuming a uniform (improper) prior on *κ*, and by introducing the parcellation labels 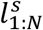 of the new subject *s* as latent variables, the lower bound 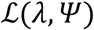 of the MAP problem (Eq. (40)) can be written as:

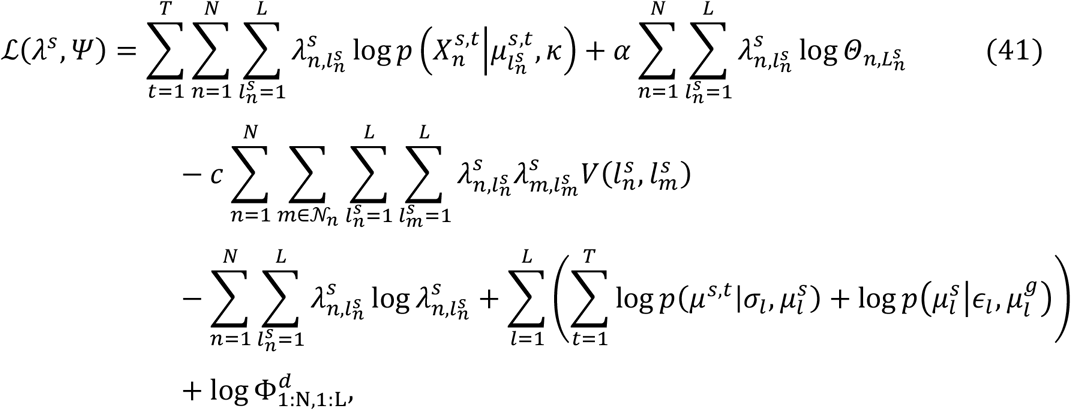

where equality is achieved when *λ^s^* is the posterior probability of the individual-specific parcellation of subject *s* given the parameters *Ψ*. Similar to Section S2.1.1, we can maximize the lower bound (Eq. (41)) by iteratively updating *λ^s^* and *Ψ*. Unlike Section S2.1.1, we cannot compute the exact posterior probability *λ^s^* because of the pairwise potentials in the Markov random field (Wainwright and Jordan, 2008). Using the mean-field approximation (Wainwright and Jordan, 2008), an approximate posterior probability *λ^s^* is estimated in the variational E-step, while *Ψ* is updated in the variational M-step.

More specifically, in the variational E-step, *Ψ* is fixed and *λ^s^* is estimated as follows:

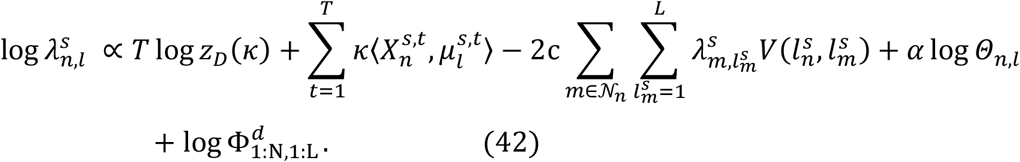

In the variational M-step, *λ^s^* is fixed and 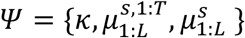 is estimated as follows:

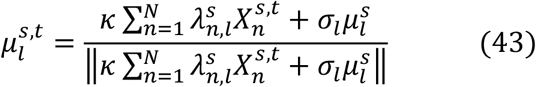

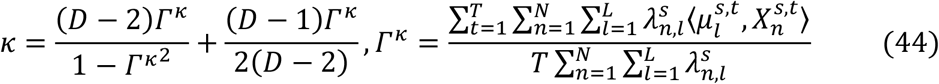

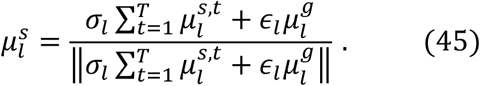

Once the VBEM algorithm converges, vertex *n* of subject *s* will be assigned to label *l* with the highest (approximate) posterior probability.

##### S2.2.1 cMS-HBM

Similar to dMS-HBM in S2.2.1, let 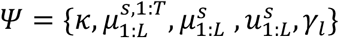, we consider the following maximum-a-posterior (MAP) estimation problem:

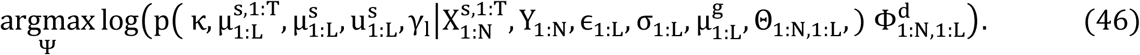

Assuming a uniform (improper) prior on *κ* and *γ_l_*, by introducing the parcellation labels 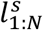 of the new subject *s* as latent variables, the lower bound 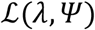 of the MAP problem (Eq. (46)) can be written as:

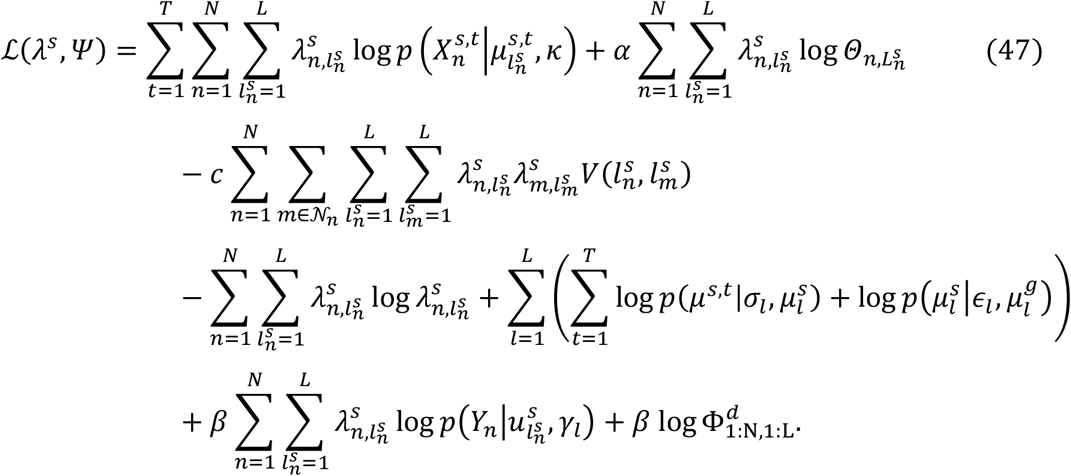

Similarly, in the variational E-step, *Ψ* is fixed and *λ^s^* is estimated as follows:

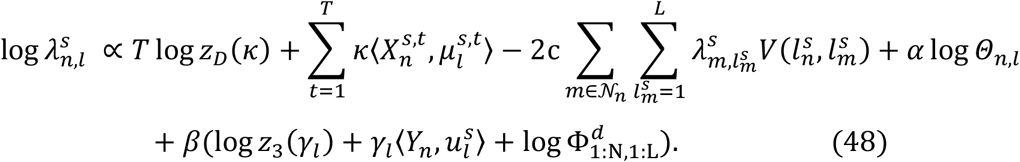

In the variational M-step, *λ^s^* is fixed and 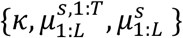 is estimated as Eqs. (43–45). 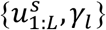 can be estimated as follows:

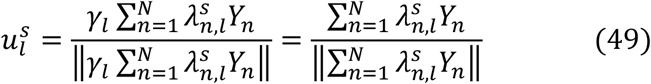

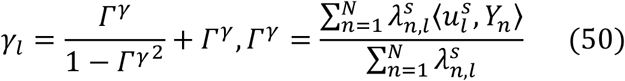

##### S2.2.1 gMS-HBM

Similar to dMS-HBM in S2.2.1, let 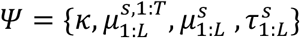, we consider the following maximum-a-posterior (MAP) estimation problem:

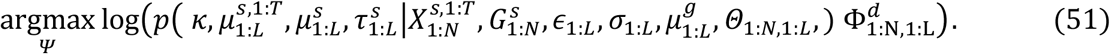

Assuming a uniform (improper) prior on *κ*, by introducing the parcellation labels 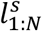 of the new subject *s* as latent variables, the lower bound 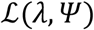 of the MAP problem (Eq. (51)) can be written as:

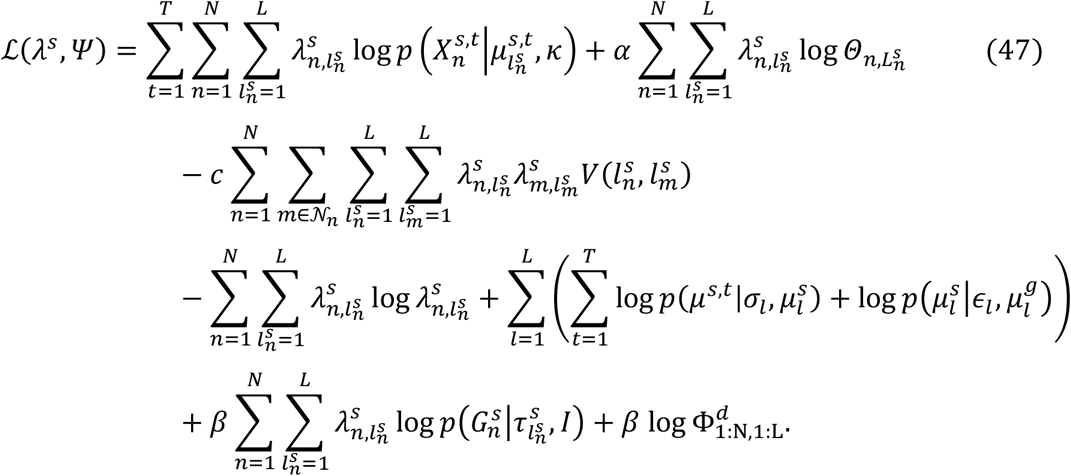

Similarly, in the variational E-step, *Ψ* is fixed and *λ^s^* is estimated as follows:

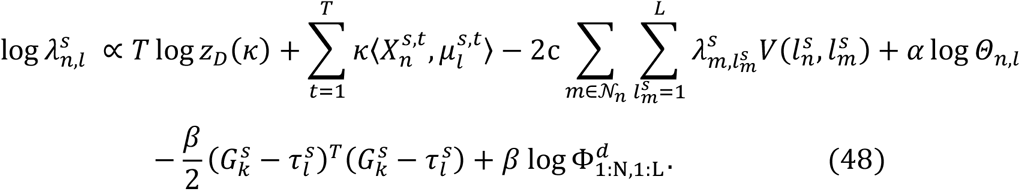

In the variational M-step, *λ^s^* is fixed and 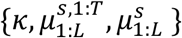 is estimated as Eqs. (43–45). 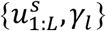 can be estimated as follows:

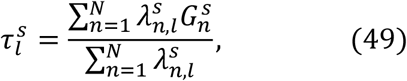

### S3. Matching algorithm for comparison with Laumann2015

We mentioned in the main text that parcellations estimated by Laumann2015 (Laumann et al., 2015; Gordon et al., 2017) had a variable number of parcels across subjects. Furthermore, Laumann2015 parcellations also had a significant number of vertices between parcels that were not assigned to any parcel, which had the effect of increasing resting homogeneity and decreasing task inhomogeneity. Therefore, we performed a post-hoc processing of MS-HBM parcellations to match the number of parcels and unlabeled vertices of Laumann2015 parcellations.

Since Laumann2015 individual-specific parcellations typically have around 500-600 parcels, we utilized 600-region Schaefer2018 group-level parcellation to initialize MS-HBM algorithms to generate 600-region MS-HBM parcellations. For each subject, we sort the 600 MS-HBM parcels based on parcel size and for parcels with the same size, we sort them based on resting-state homogeneity. Assume Laumann2015 parcellation has M parcels, we find the 600-M MS-HBM parcels with the smallest size and the lowest resting-state homogeneity. For each vertex *i* within these 600-M parcels, we re-assigned it to the neighboring parcel *p* with the highest similarity 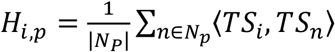, where *N_p_* was the set of vertices within parcel *p, TS_i_* was the time course of vertex *i*. After the re-assignment procedure, a 2-vertex thick parcellation boundary was created for MS-HBM. Let’s denote *N*_2*thick*_ as the number of 2-vertex thick parcellation boundary vertices. Suppose there were W vertices that were unlabeled in Lauman2015 parcellation. If W was less than *N*_2*thick*_, we removed the labels from the top W vertices within the 2-vertex thick boundary with the lowest resting-state homogeneity. If W was more than *N*_2*thick*_, we first removed the labels from all the vertices within the 2-vertex thick boundary, and then remove the labels from the remaining W-*N*_2*thick*_ vertices, which were the nearest neighbors of the 2-vertex thick boundary vertices with the lowest resting-state homogeneity. In this way, we can match the MS-HBM parcellations to the Laumann2015 parcellations with the same number of parcels and the same number of unlabeled vertices.

### S4. Behavioral prediction model

In this section, we describe our model for behavioral prediction based on individual differences in the functional connectivity. Kernel regression (Murphy et al., 2012) was utilized to predict each behavioral phenotype in individual subjects. The derivations of our behavioral prediction model has been shown in our previous work (Kong et al., 2019). Here, we include the derivation again for completeness.

Suppose we have *M* training subjects, *y_i_* is the behavioral measure (e.g., fluid intelligence) and *FC_i_* is the 400 × 400 RSFC matrix generated by individual-specific parcellation of the *i*-th training subject. Given {*y*_1_, *y*_2_,…, *y_M_*} and {*FC*_1_, *FC*_2_,…, *FC_M_*}, the kernel regression model will be:

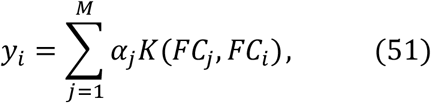

where *K*(*FC_j_, FC_i_*) is the Pearson correlation between the RSFC matrices of the *i*-th and *j*-th training subjects. The classical way to estimate *α* in Eq. (51) is to minimize the quadratic cost:

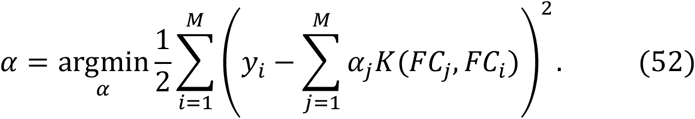

Defining ***y*** = [*y*_1_, *y*_2_,…, *y_M_*]^*T*^, ***α*** = [*α*_1_, *α*_2_,…, *α*_3_]^*T*^ and 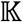 to be an *M* × *M* matrix, whose (*j, i*)-th element is *K*(*FC_j_*, *FC_i_*), Eq. (52) can be written as:

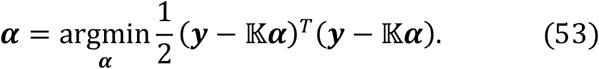

Differentiating Eq. (53) with respect to ***α***, we can get

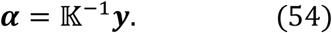

To predict the behavior measure *y_s_* (e.g., fluid intelligence) of a test subject *s* with its RSFC matrix *FC_S_*, we can compute *K*(*FC_i_, FC_s_*), which is the Pearson correlation between RSFC matrix of subject *s* and *i*-th training subject. The predicted behavior measure *y_s_* can be calculated as

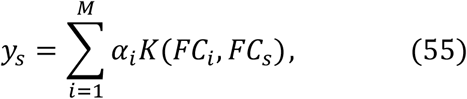

where *α_i_* is estimated by Eq. (53). If we denote ***K**_s_* = [*K*(*FC*_1_, *FC_s_*), *K*(*FC*_2_, *FC_s_*),…, *K*(*FC_M_, FC_s_*)], then Eq. (55) can be written as:

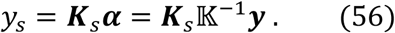

In practice, 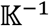 is a symmetric matrix whose diagonal elements are roughly the same and ~100 times larger than the off-diagonal elements. Therefore, the predicted behavior measure *y_s_* can be seen as the weighted average of the behaviors of the training subjects: 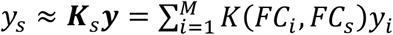. If the RSFC matrix *FC_S_* of test subject *s* is more similar to the RSFC matrix of training subject *i* than training subject *j*, then weight *K*(*FC_i_, FC_S_*) will be larger than *K*(*FC_j_, FC_S_*), and so *y_s_* will be more similar to *y_i_* than *y_j_*.

To reduce overfitting, an *l*_2_-regularization term (i.e., kernel ridge regression) is typically added to cost function (Eq. (53)), resulting in a new regularized cost function:

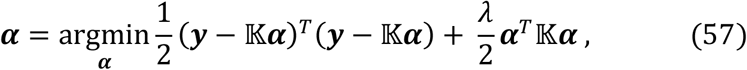

where *λ* is a tuning parameter, which controls the importance of the regularization term. Differentiating Eq. (57) with respect to ***α***, we get

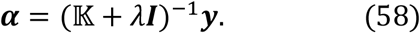

To predict the behavior measure *y_s_* (e.g., fluid intelligence) of a test subject *s*, Eq. (58) is substituted into Eq. (55), resulting in

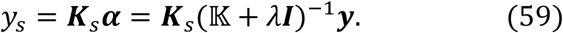

## Supplemental Results

**Figure S1.**
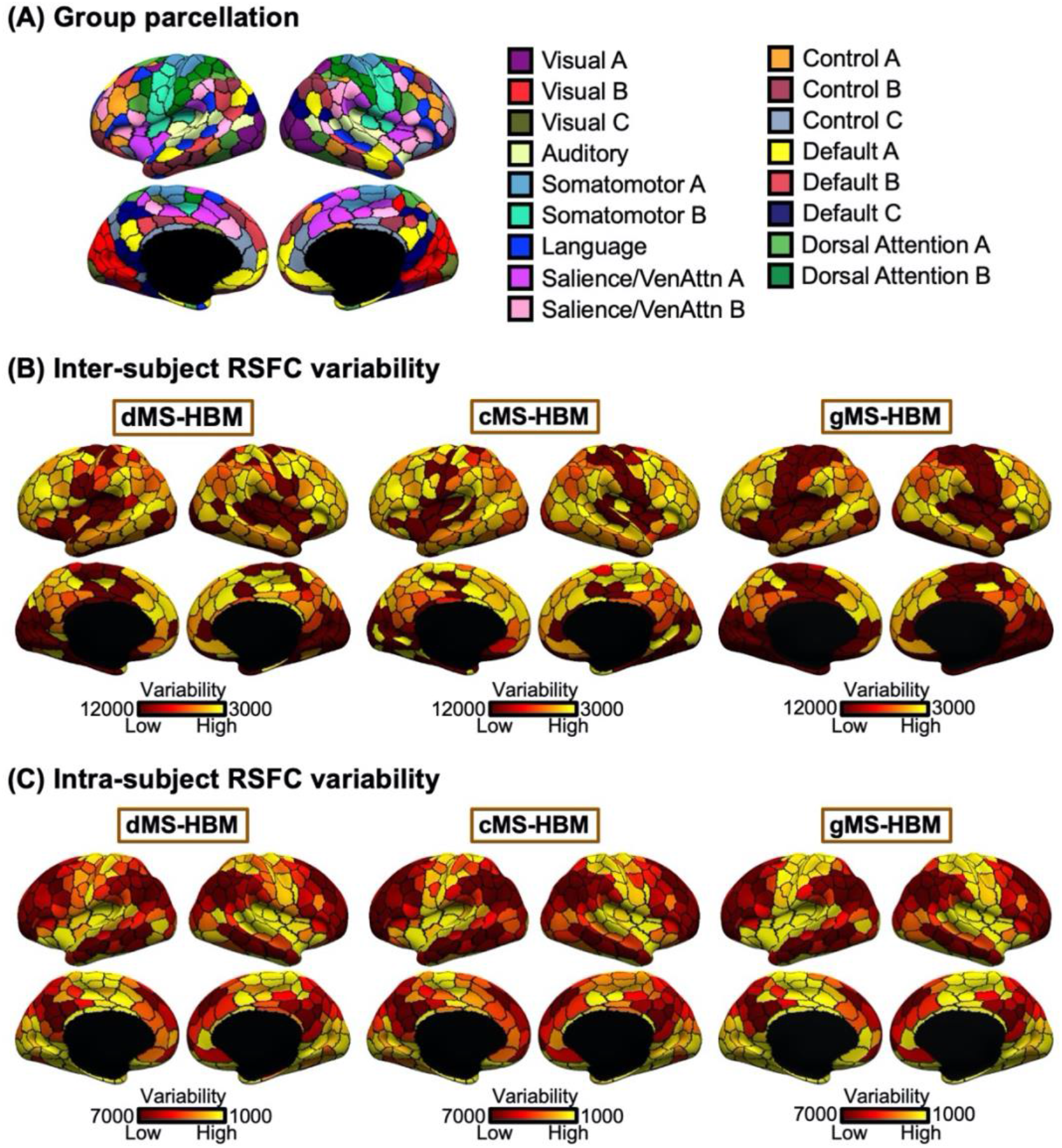
Sensory-motor parcels exhibit lower inter-subject, but higher intra-subject, functional connectivity variability than association cortical parcels for in the HCP training set. (A) 400-region Schaefer2018 group-level parcellation. (B) Inter-subject resting-state functional connectivity variability for different parcels. (C) Intra-subject resting-state functional connectivity variability for different parcels. Note that (B) and (C) correspond to the *ϵ_l_* and *σ_l_* parameters in Figure 1, where higher values indicate lower variability.

**Figure S2.**
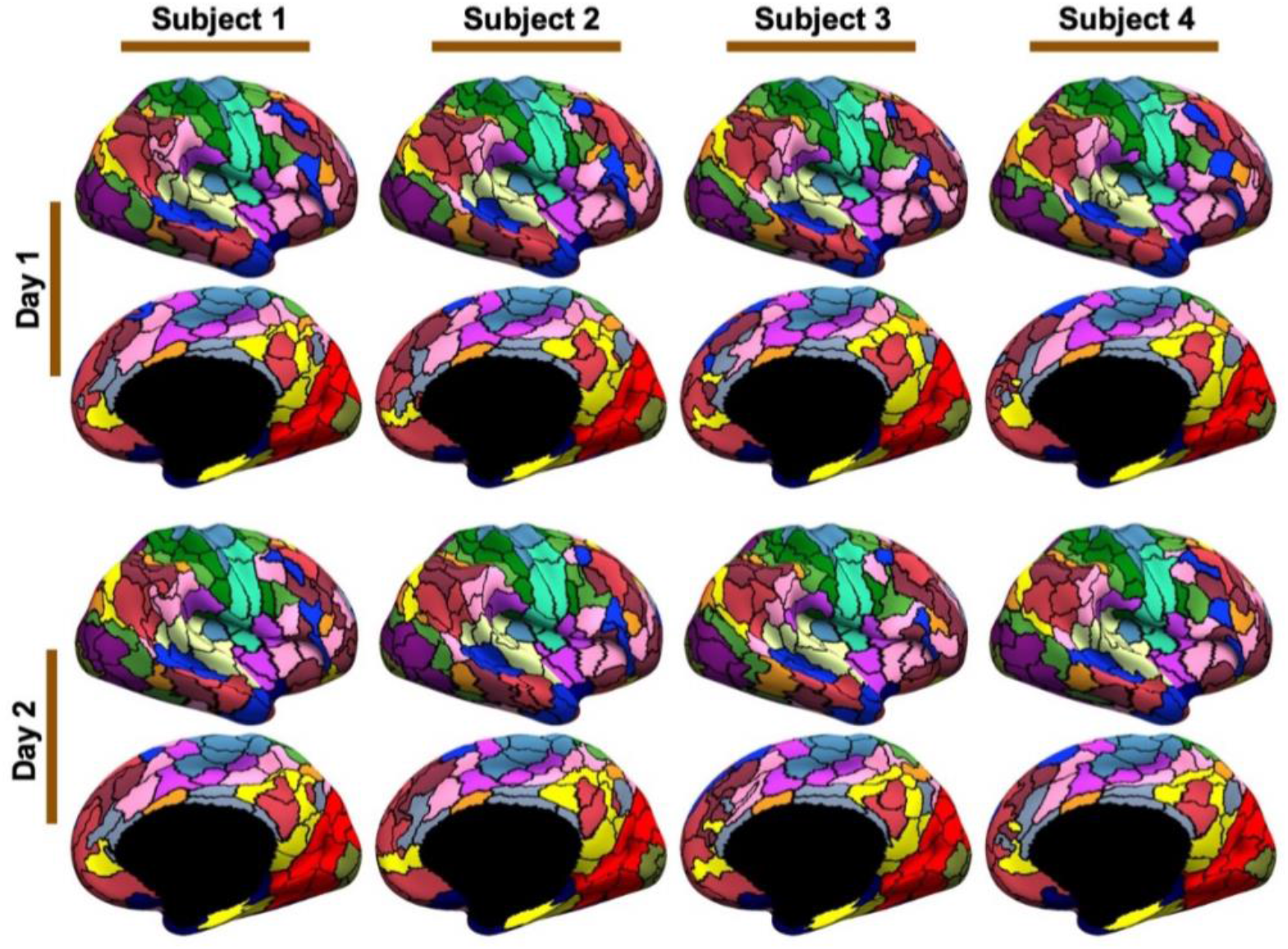
400-region individual-specific areal-level parcellations were estimated using rs-fMRI data from day 1 and day 2 separately for each participant from HCP test set. Right hemisphere parcellations of four representative participants are shown here. Left hemisphere parcellations are shown in Figure 2C.

**Figure S3.**
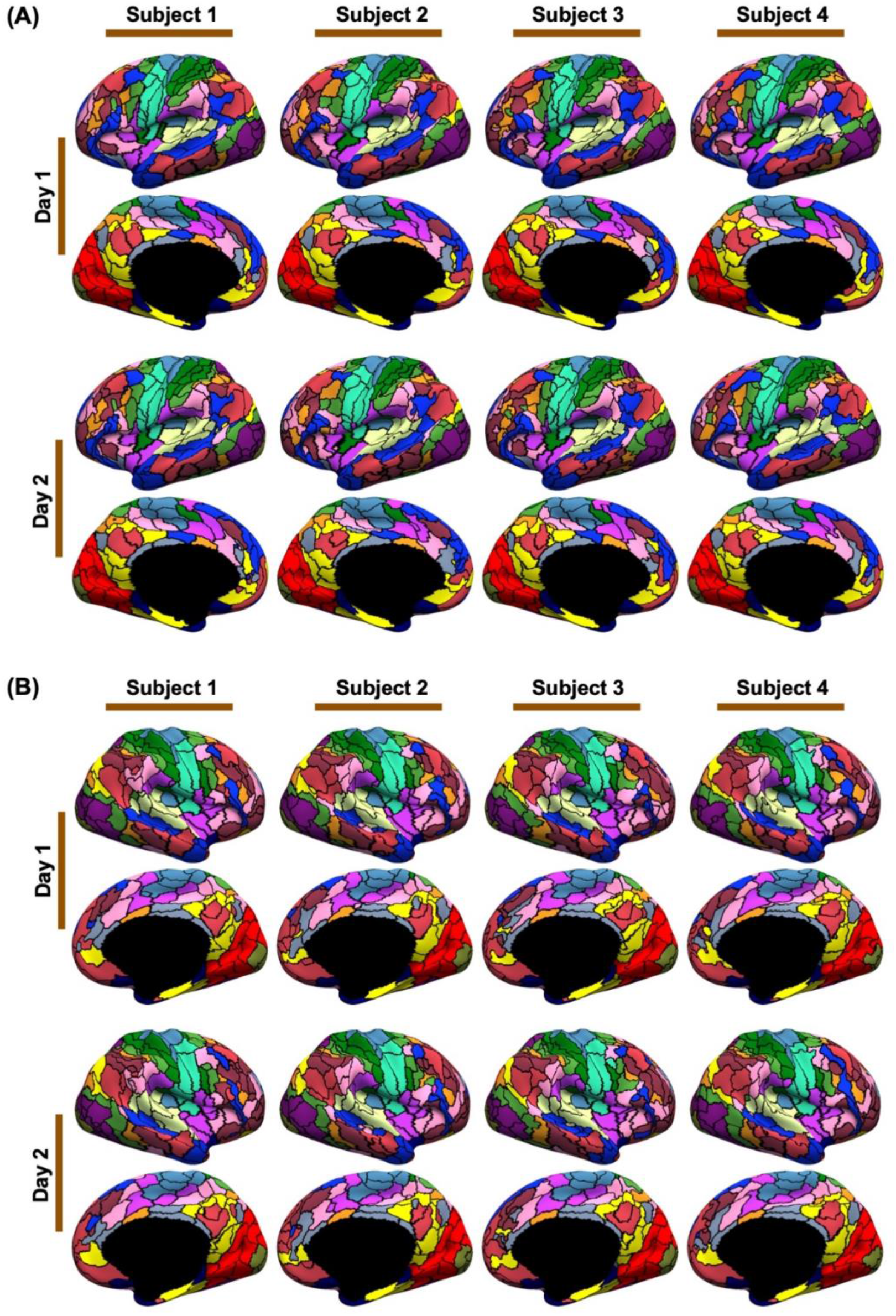
400-region individual-specific dMS-HBM parcellations were estimated using rs-fMRI data from day 1 and day 2 separately for each participant from HCP test set. (A) Left and (B) right hemisphere parcellations of four representative participants are shown here.

**Figure S4.**
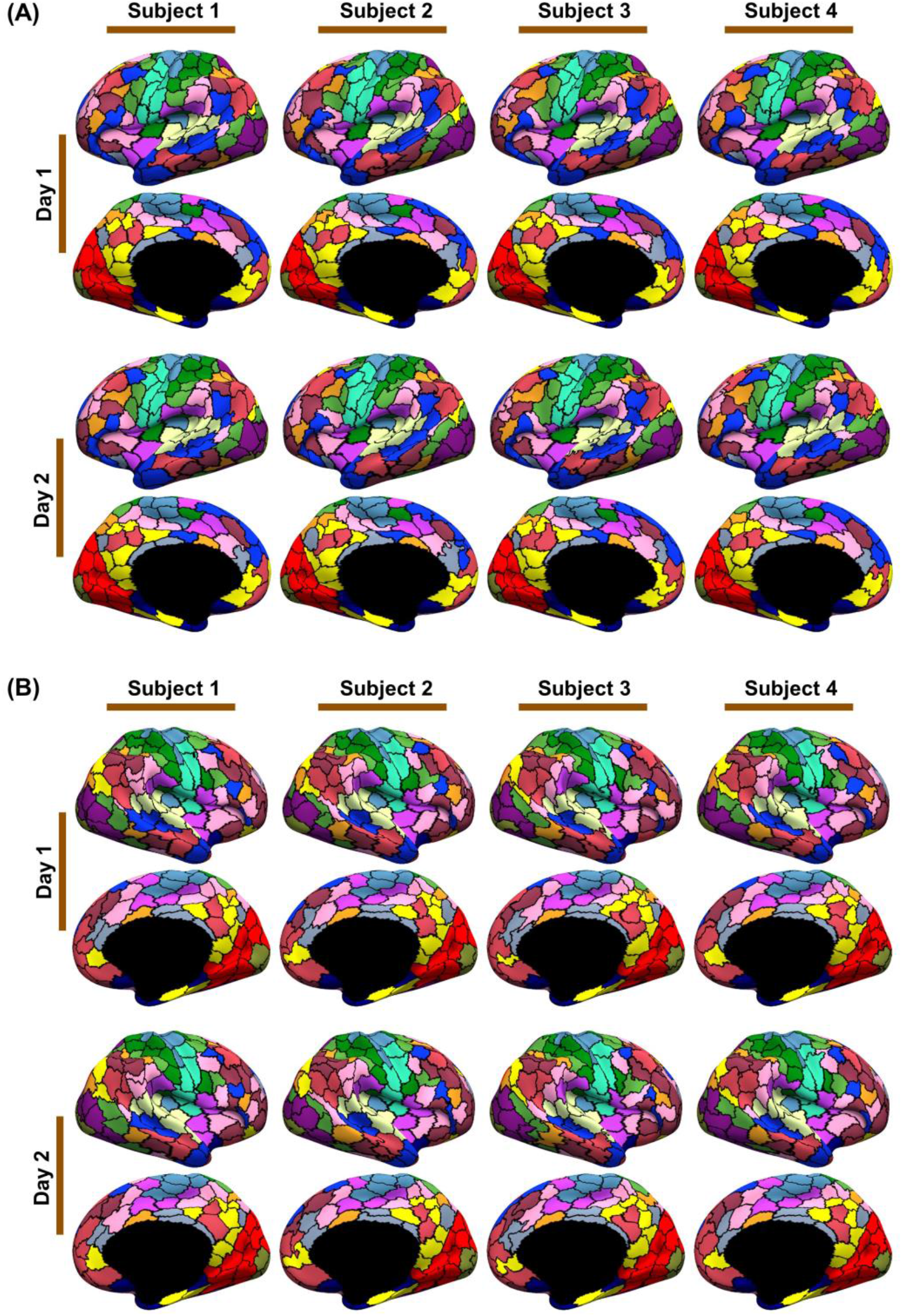
400-region individual-specific cMS-HBM parcellations were estimated using rs-fMRI data from day 1 and day 2 separately for each participant from HCP test set. (A) Left and (B) right hemisphere parcellations of four representative participants are shown here.

**Figure S5.**
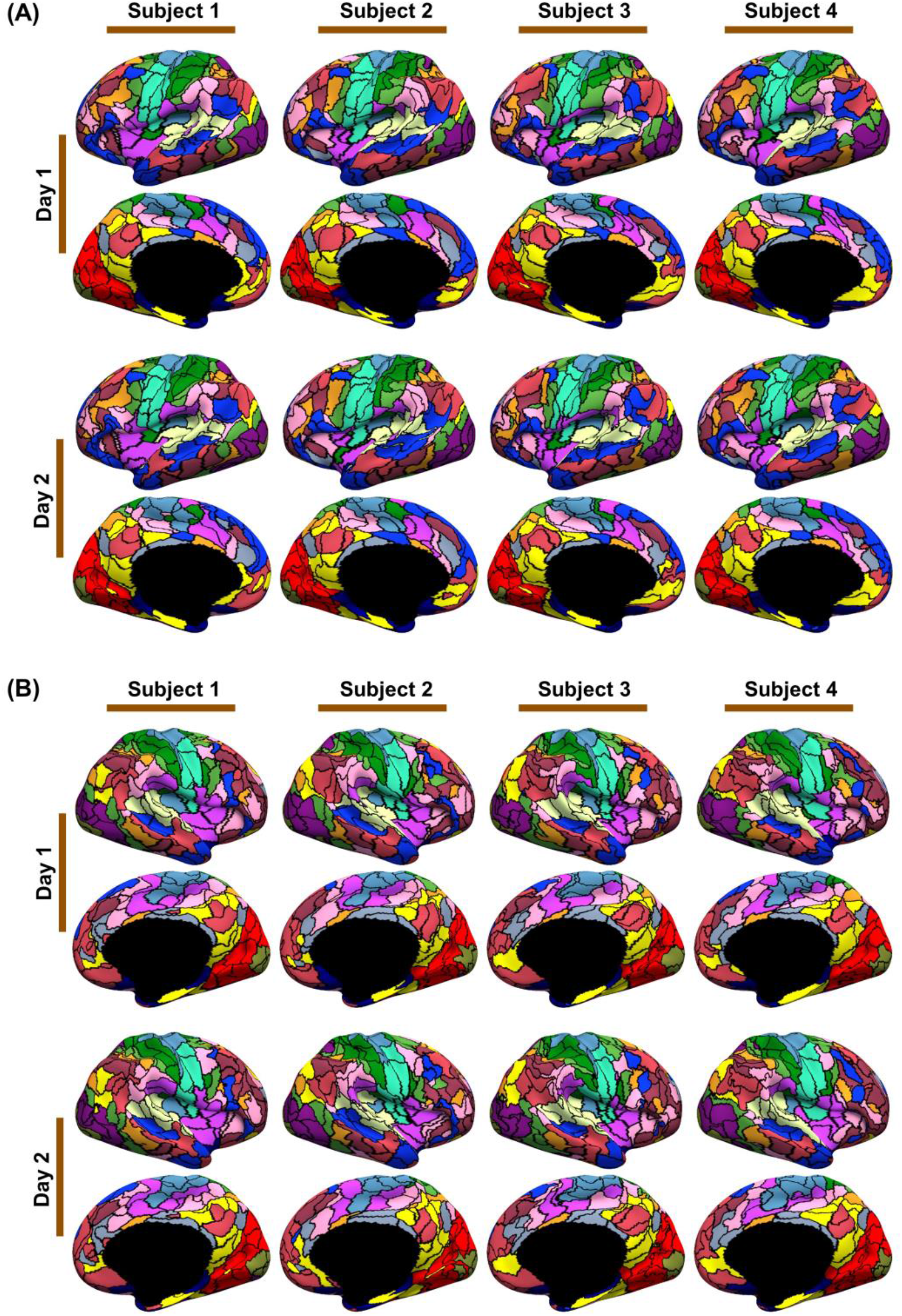
400-region individual-specific dMS-HBM parcellations were estimated using rs-fMRI data from sessions 1-5 and sessions 6-10 separately for each participant from MSC dataset. (A) Left and (B) right hemisphere parcellations of four representative participants are shown here.

**Figure S6.**
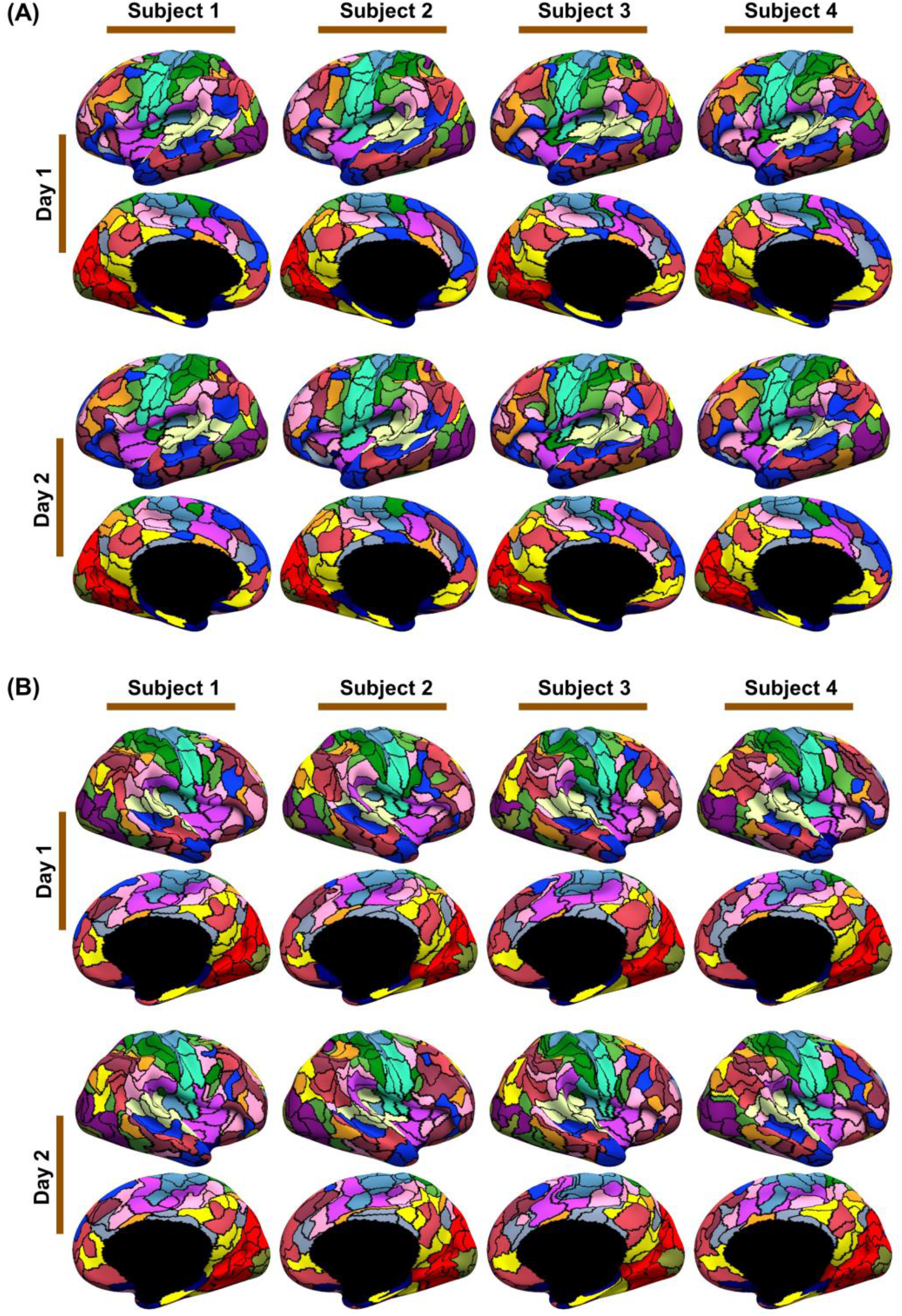
400-region individual-specific cMS-HBM parcellations were estimated using rs-fMRI data from sessions 1-5 and sessions 6-10 separately for each participant from MSC dataset. (A) Left and (B) right hemisphere parcellations of four representative participants are shown here.

**Figure S7.**
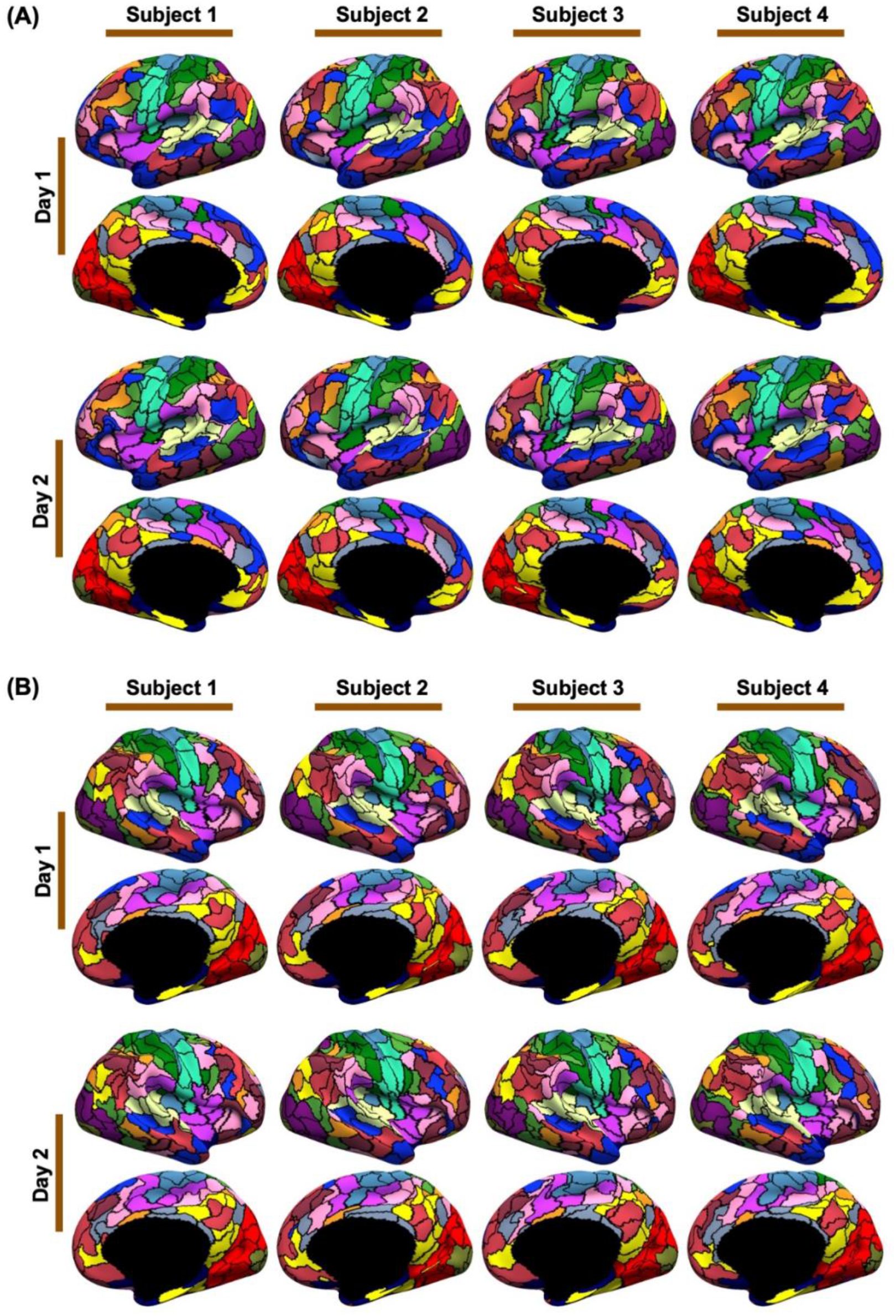
400-region individual-specific gMS-HBM parcellations were estimated using rs-fMRI data from sessions 1-5 and sessions 6-10 separately for each participant from MSC dataset. (A) Left and (B) right hemisphere parcellations of four representative participants are shown here.

**Figure S8.**
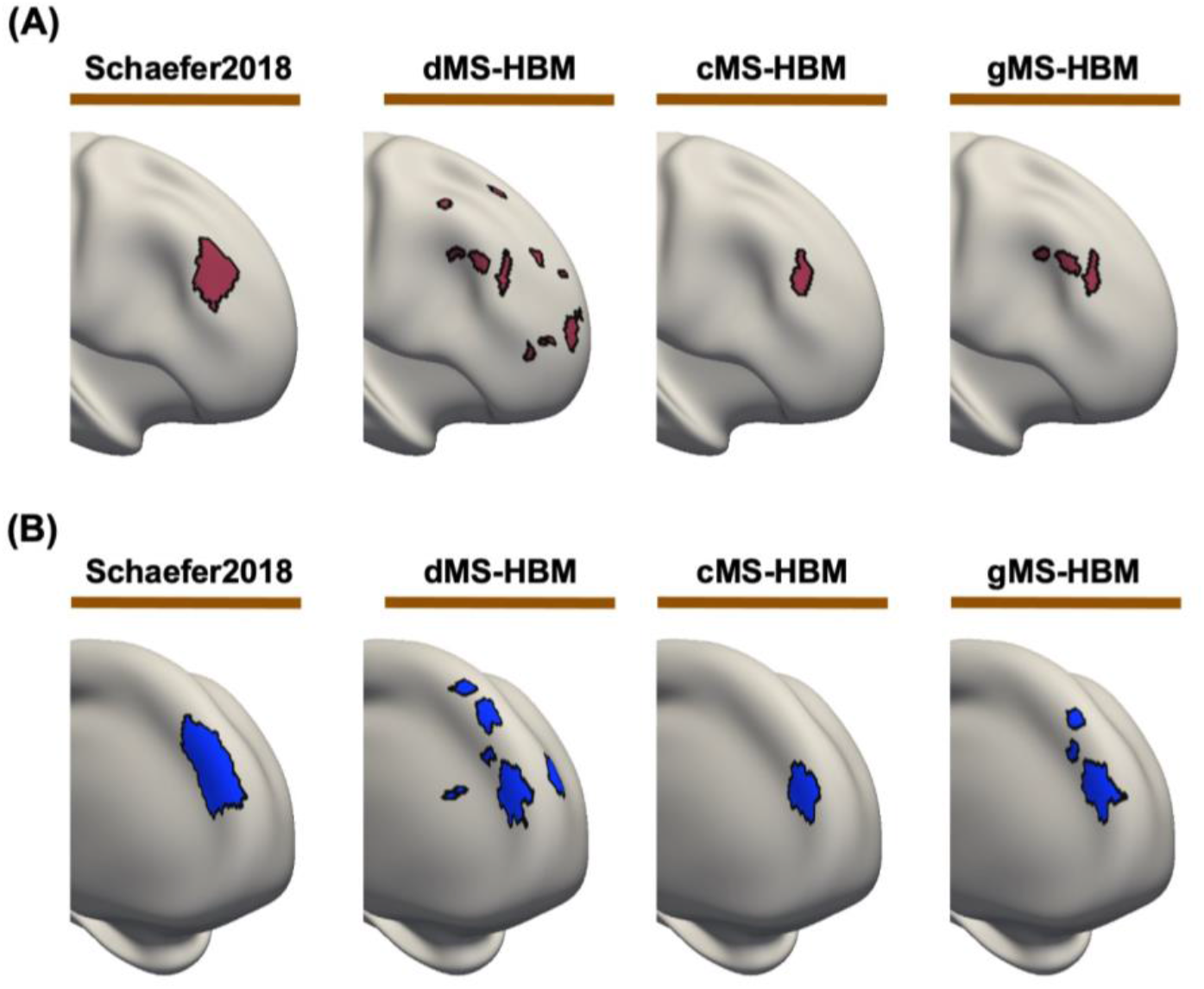
Parcels with the maximum number of spatially disconnected components for (A) dMS-HBM and (B) gMS-HBM. The maximum number of spatially disconnected components for dMS-HBM and gMS-HBM is 11 and 3.

**Figure S9.**
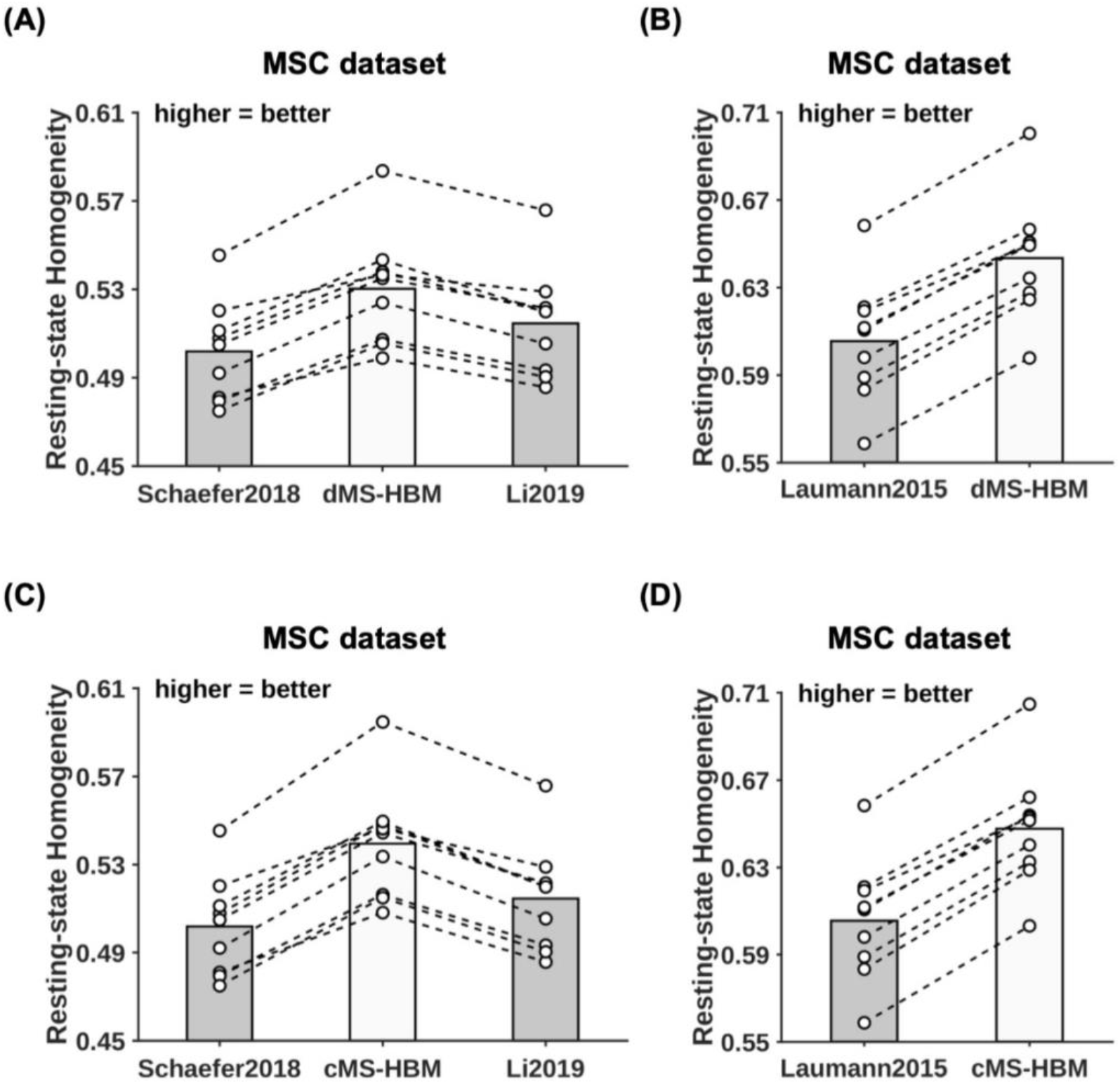
MS-HBM parcellations achieved better out-of-sample resting-state homogeneity than other approaches in the MSC dataset. (A, C) 400-region individual-specific parcellations were estimated using a single rs-fMRI session and resting-state homogeneity were computed with the remaining sessions for each MSC participant. Each circle represents one MSC participant. Dash lines connect the same participants. (B, D) Same as (A, C) except that Laumann2015 yielded different number of parcels for each participant, so we matched the number of MS-HBM parcels accordingly for each participant.

**Figure S10.**
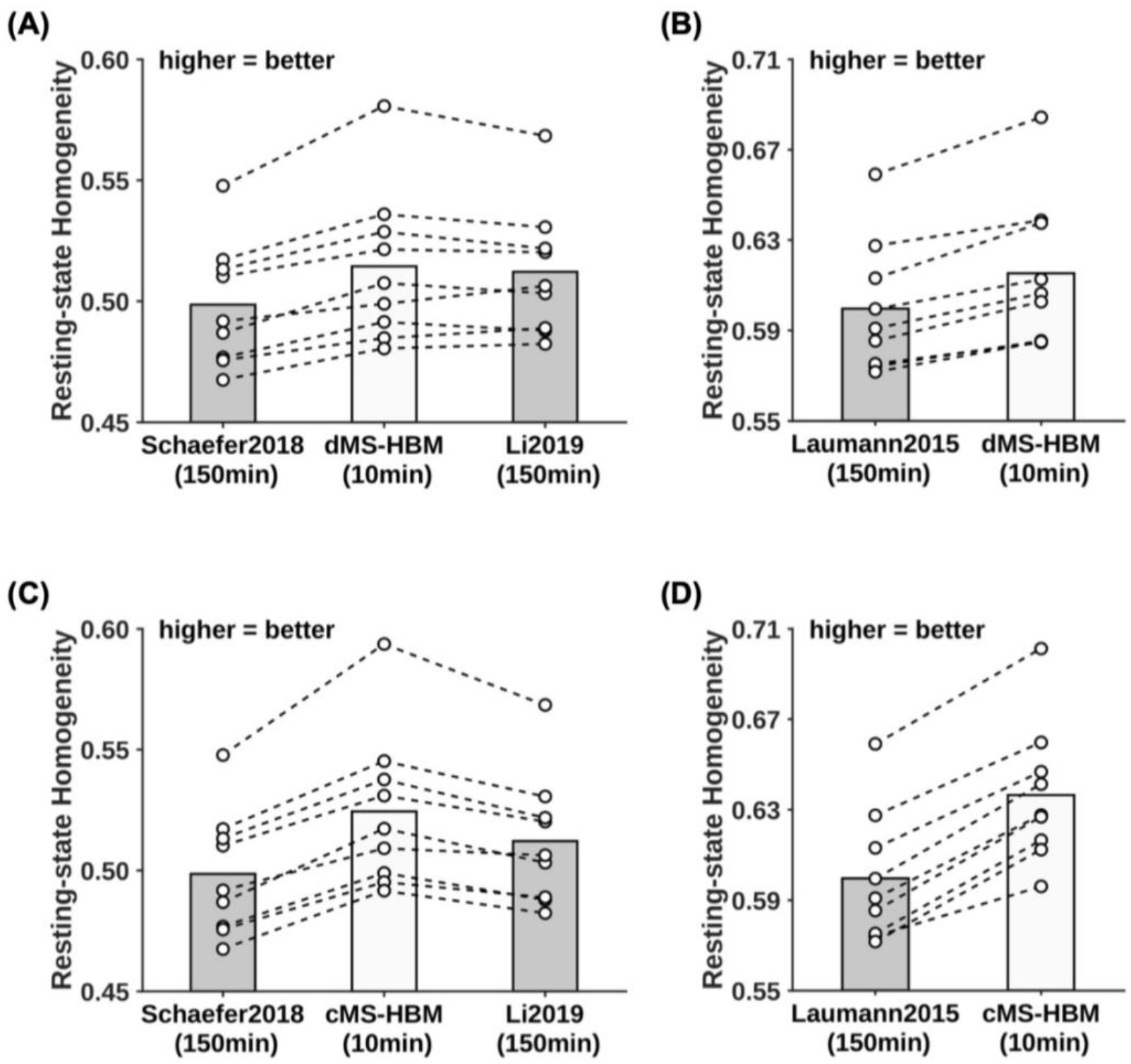
MS-HBM parcellations achieved better out-of-sample resting-state homogeneity with less amount of data. (A, C) 400-region individual-specific parcellations were estimated for each MSC participant using 10 min of rs-fMRI data for gMS-HBM and 150 min of rs-fMRI data for Li2019. Resting-state homogeneity was evaluated using leave-out sessions. (B, D) Same as (A, C) except that Laumann2015 yielded different number of parcels for each participant, so we matched the number of MS-HBM parcels accordingly for each participant.

**Figure S11.**
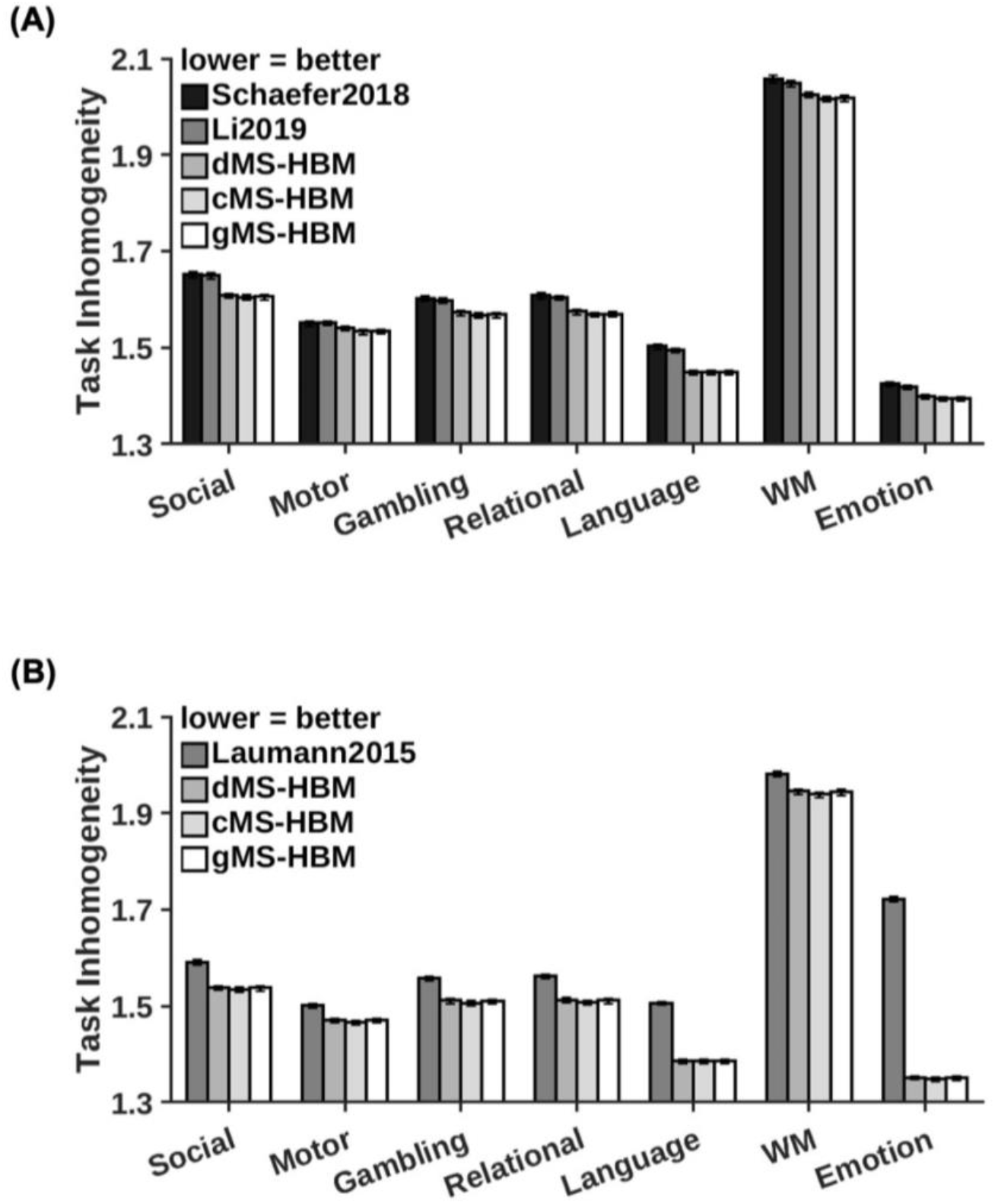
MS-HBM parcellations achieved better task inhomogeneity in the HCP dataset. (A) 400-region individual-specific parcellations were estimated using all resting-state fMRI data. Task inhomogeneity was evaluated in the task data. Task inhomogeneity was then defined as the standard deviation of task activation within each parcel, and then averaged across all parcels and contrasts within each behavioral domain. Lower value indicates better task inhomogeneity. Error bars correspond to standard errors. (B) Same as (A) except that Laumann2015 yielded different number of parcels for each participant, so we matched the number of MS-HBM parcels accordingly for each participant. Compared with Schaefer2018, Laumann2015 and Li2019, dMS-HBM achieved an improvement of 2.1% (Cohen’s d > 0.82 for all domains), 7.3% (Cohen’s d > 0.53 for all domains) and 1.8% (Cohen’s d > 1.0 for all domains) respectively; cMS-HBM achieved an improvement of 2.4% (Cohen’s d > 1.5 for all domains), 7.6% (Cohen’s d > 0.59 for all domains) and 2.1% (Cohen’s d > 1.8 for all domains) respectively; gMS-HBM achieved an improvement of 2.3% (Cohen’s d > 1.4 for all domains), 7.4% (Cohen’s d > 1.7 for all domains) and 2.0% (Cohen’s d > 1.4 for all domains) respectively.

**Table S1.**
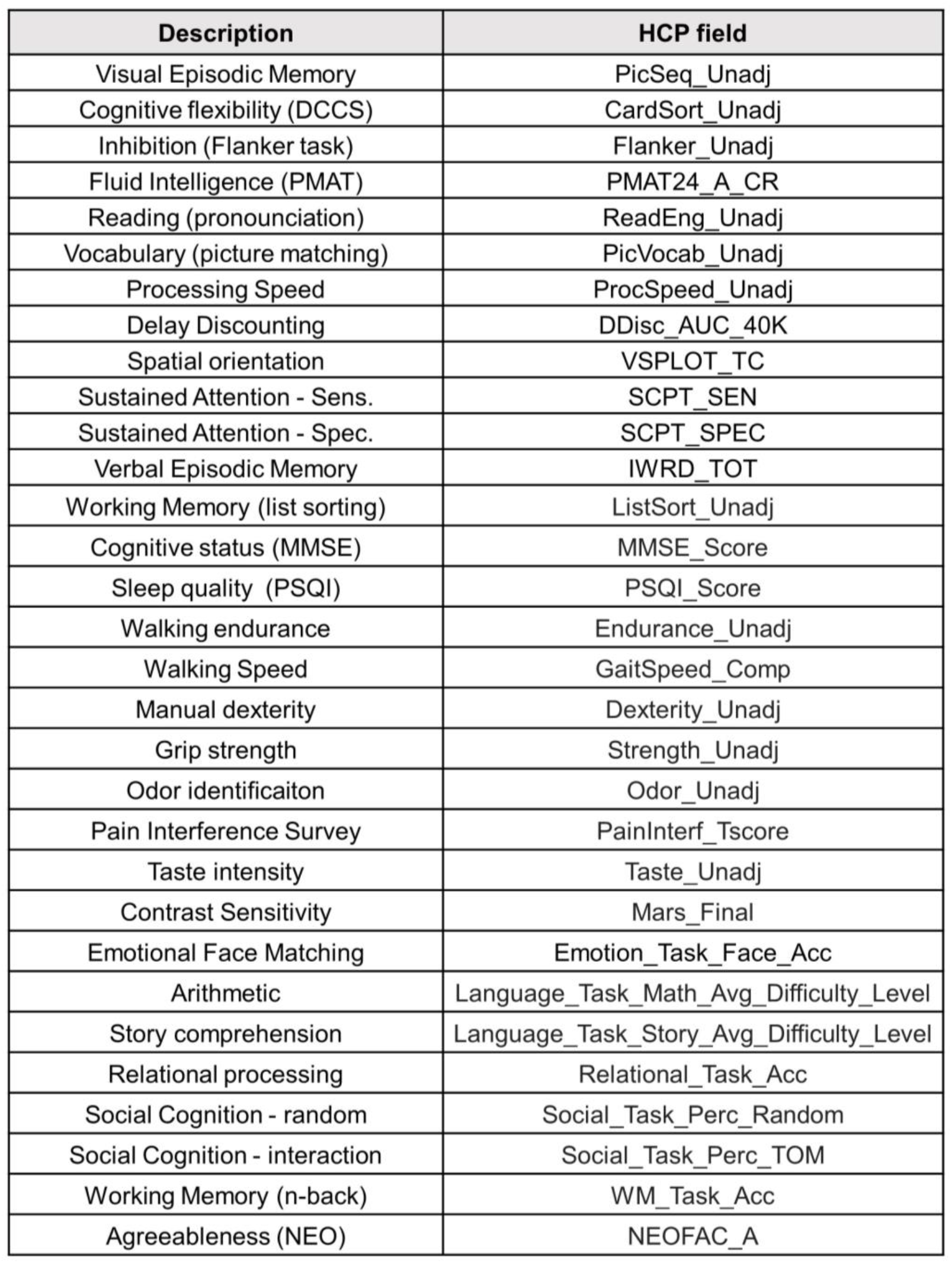
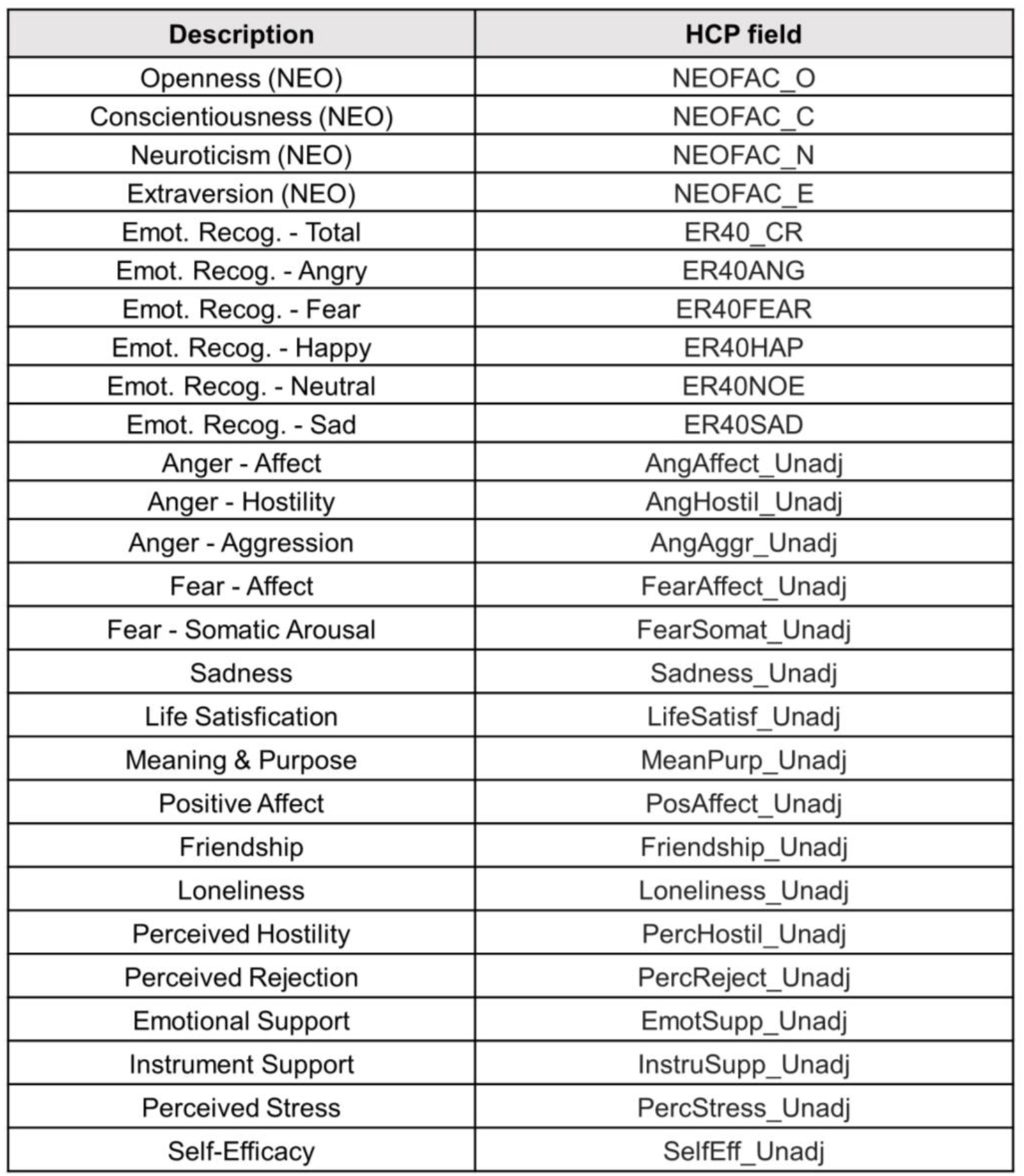
Lookup table showing the original HCP variable names with the corresponding descriptive labels used in the manuscript. More details of the behavioral measures can be found in the HCP data dictionary.

**Table S2.** Average prediction accuracies (Pearson’s correlation) for all 58 behavioral measures and 36 behaviors with accuracies higher than 0.1 for at least one approach (“36 behaviors > 0.1”) across different parcellation approaches. Prediction was based on individual-specific functional connectivity. The mean accuracy and standard deviation were calculated across 100 20-fold cross-validations. The percentage improvement and number of cross-validations that MS-HBM algorithms outperform Schaefer2018 and Li2019 across 100 20-fold cross-validations (shown in brackets) were reported. Red font indicates statistical significance after correcting for multiple comparisons with false discovery rate q < 0.05.

|  |  | All 58 behaviors | 36 behaviors >0.1 |
| --- | --- | --- | --- |
| Accuracy<br>(correlation)<br>across 100<br>20-fold CVs<br>(mean ± std) | gMS-HBM | 0.1111±0.0031 | 0.1656±0.0036 |
| | cMS-HBM | 0.1062±0.0031 | 0.1610±0.0036 |
| | dMS-HBM | 0.1083±0.0031 | 0.1654±0.0034 |
| | Schaefer2018 | 0.0960±0.0031 | 0.1471±0.0036 |
| | Li2019 | 0.0944±0.0031 | 0.1465±0.0034 |
| Percentage<br>improvement<br>(#CV splits<br>MS-HBM<br>outperforms<br>Schaefer2018) | gMS-HBM<br>vs<br>Schaefer2018 | 15.81%<br>(100/100)<br>(p=0.0005) | 12.59%<br>(100/100)<br>(p=0.0002) |
|  | cMS-HBM<br>vs<br>Schaefer2018 | 10.65%<br>(100/100)<br>(p=0.0251) | 9.47%<br>(100/100)<br>(p=0.0076) |
|  | dMS-HBM<br>vs<br>Schaefer2018 | 12.88%<br>(100/100)<br>(p=0.0057) | 12.44%<br>(100/100)<br>(p=0.0003) |
| Percentage<br>improvement<br>(#CV splits<br>MS-HBM<br>outperforms<br>Li2019) | gMS-HBM<br>vs<br>Li2019 | 17.80%<br>(100/100)<br>(p=0.0005) | 13.09%<br>(100/100)<br>(p=0.0004) |
|  | cMS-HBM<br>vs<br>Li2019 | 12.55%<br>(100/100)<br>(p=0.0171) | 9.96%<br>(100/100)<br>(p=0.0112) |
|  | dMS-HBM<br>vs<br>Li2019 | 14.83%<br>(100/100)<br>(p=0.0035) | 12.94%<br>(100/100)<br>(p=0.0006) |

**Table S3.** Average prediction accuracies (Pearson’s correlation) for 13 cognitive measures, NEO-5 personality measures and emotional measures across different parcellation approaches. Prediction was based on individual-specific functional connectivity. The mean accuracy and standard deviation were calculated across 100 20-fold cross-validations. The percentage improvement and number of cross-validations that MS-HBM algorithms outperform Schaefer2018 and Li2019 across 100 20-fold cross-validations (shown in brackets) were reported. Red font indicates statistical significance after correcting for multiple comparisons with false discovery rate q < 0.05.

|  |  | 13 Cognitive measures | NEO-5 | Emotional recognition (ER) |
| --- | --- | --- | --- | --- |
| Accuracy (correlation) across 100 20-fold CVs (mean ± std) | gMS-HBM | 0.1848±0.0050 | 0.1504±0.0072 | 0.0808±0.0050 |
| | cMS-HBM | 0.1738±0.0049 | 0.1472±0.0080 | 0.0708±0.0053 |
| | dMS-HBM | 0.1828±0.0049 | 0.1294±0.0070 | 0.0766±0.0049 |
| | Schaefer2018 | 0.1795±0.0042 | 0.1124±0.0085 | 0.0535±0.0050 |
| | Li2019 | 0.1750±0.0043 | 0.1135±0.0077 | 0.0499±0.0048 |
| Percentage improvement (#CV splits MS-HBM outperforms Schaefer2018) | gMS-HBM vs Schaefer2018 | 2.94%<br>(96/100)<br>(p=0.4415) | 34.15%<br>(100/100)<br>(p=0.0014) | 51.90%<br>(100/100)<br>(p=0.0007) |
|  | cMS-HBM vs Schaefer2018 | -3.20%<br>(5/100)<br>(p=0.4598) | 31.26%<br>(100/100)<br>(p=0.0054) | 33.02%<br>(100/100)<br>(p=0.0443) |
|  | dMS-HBM vs Schaefer2018 | 1.85%<br>(83/100)<br>(p=0.6364) | 15.39%<br>(100/100)<br>(p=0.1464) | 44.10%<br>(100/100)<br>(p=0.0044) |
| Percentage improvement (#CV splits MS-HBM outperforms Li2019) | gMS-HBM vs Li2019 | 5.60%<br>(100/100)<br>(p=0.1670) | 32.82%<br>(100/100)<br>(p=0.0025) | 62.99%<br>(100/100)<br>(p=0.0003) |
|  | cMS-HBM vs Li2019 | -0.69%<br>(39/100)<br>(p=0.8746) | 29.92%<br>(100/100)<br>(p=0.0068) | 42.67%<br>(100/100)<br>(p=0.0208) |
|  | dMS-HBM vs Li2019 | 4.48%<br>(100/100)<br>(p=0.2818) | 14.22%<br>(100/100)<br>(p=0.1887) | 54.56%<br>(100/100)<br>(p=0.0017) |

**Table S4.** Average prediction accuracies (COD) for all 58 behavioral measures and 36 behaviors with accuracies higher than 0.1 for at least one approach (“36 behaviors > 0.1”) across different parcellation approaches. Prediction was based on individual-specific functional connectivity. The mean accuracy and standard deviation were calculated across 100 20-fold cross-validations. The percentage improvement and number of cross-validations that MS-HBM algorithms outperform Schaefer2018 and Li2019 across 100 20-fold cross-validations (shown in brackets) were reported. Red font indicates statistical significance after correcting for multiple comparisons with false discovery rate q < 0.05.

|  |  | All 58 behaviors | 36 behaviors >0.1 |
| --- | --- | --- | --- |
| Accuracy (COD) across 100 20-fold CVs (mean ± std) | gMS-HBM | 0.0156±0.0010 | 0.0266±0.0014 |
| | cMS-HBM | 0.0149±0.0009 | 0.0257±0.0014 |
| | dMS-HBM | 0.0147±0.0009 | 0.0252±0.0014 |
| | Schaefer2018 | 0.0120±0.0009 | 0.0212±0.0014 |
| | Li2019 | 0.0121±0.0009 | 0.0213±0.0014 |
| Percentage improvement (#CV splits MS-HBM outperforms Schaefer2018) | gMS-HBM vs Schaefer2018 | 29.79% (100/100) (p=0.0026) | 25.41% (100/100) (p=0.0028) |
|  | cMS-HBM vs Schaefer2018 | 24.21% (100/100) (p=0.0207) | 21.22% (100/100) (p=0.0198) |
|  | dMS-HBM vs Schaefer2018 | 22.17% (100/100) (p=0.0195) | 18.97% (100/100) (p=0.0202) |
| Percentage improvement (#CV splits MS-HBM outperforms Li2019) | gMS-HBM vs Li2019 | 28.90% (100/100) (p=0.0051) | 24.79% (100/100) (p=0.0049) |
|  | cMS-HBM vs Li2019 | 23.36% (100/100) (p=0.0312) | 20.63% (100/100) (p=0.0289) |
|  | dMS-HBM vs Li2019 | 21.33% (100/100) (p=0.0324) | 18.39% (100/100) (p=0.0332) |

**Table S5.** Average prediction accuracies (COD) for 13 cognitive measures, NEO-5 personality measures and emotional measures across different parcellation approaches. Prediction was based on individual-specific functional connectivity. The mean accuracy and standard deviation were calculated across 100 20-fold cross-validations. The percentage improvement and number of cross-validations that MS-HBM algorithms outperform Schaefer2018 and Li2019 across 100 20-fold cross-validations (shown in brackets) were reported. Red font indicates statistical significance after correcting for multiple comparisons with false discovery rate q < 0.05.

|  |  | 13 Cognitive measures | NEO-5 | Emotional recognition (ER) |
| --- | --- | --- | --- | --- |
| Accuracy (COD) across 100 20-fold CVs (mean ± std) | gMS-HBM | 0.0412±0.0020 | 0.0202±0.0028 | 0.0032±0.0014 |
| | cMS-HBM | 0.0377±0.0021 | 0.0181±0.0029 | 0.0041±0.0013 |
| | dMS-HBM | 0.0409±0.0020 | 0.0135±0.0025 | 0.0021±0.0015 |
| | Schaefer2018 | 0.0384±0.0022 | 0.0086±0.0027 | -0.0016±0.0012 |
| | Li2019 | 0.0366±0.0021 | 0.0087±0.0024 | -0.0002±0.0012 |
| Percentage improvement (#CV splits MS-HBM outperforms Schaefer2018) | gMS-HBM vs Schaefer2018 | 7.33%<br>(99/100)<br>(p=0.3331) | 156.85%<br>(100/100)<br>(p=0.0027) | 976.79%<br>(100/100)<br>(p=0.0049) |
|  | cMS-HBM vs Schaefer2018 | -1.82%<br>(31/100)<br>(p=0.8214) | 129.41%<br>(100/100)<br>(p=0.0151) | 1113.46%<br>(100/100)<br>(p=0.0023) |
|  | dMS-HBM vs Schaefer2018 | 6.72%<br>(98/100)<br>(p=0.4079) | 68.76%<br>(100/100)<br>(p=0.1681) | 742.63%<br>(100/100)<br>(p=0.0130) |
| Percentage improvement (#CV splits MS-HBM outperforms Li2019) | gMS-HBM vs Li2019 | 12.58%<br>(100/100)<br>(p=0.1122) | 146.07%<br>(100/100)<br>(p=0.0033) | 887.80%<br>(100/100)<br>(p=0.0469) |
|  | cMS-HBM vs Li2019 | 2.96%<br>(77/100)<br>(p=0.7384) | 120.15%<br>(100/100)<br>(p=0.0167) | 1109.74%<br>(100/100)<br>(p=0.0206) |
|  | dMS-HBM vs Li2019 | 11.94%<br>(100/100)<br>(p=0.1567) | 62.72%<br>(100/100)<br>(p=0.2047) | 626.97%<br>(100/100)<br>(p=0.1116) |

## Footnotes

1 A gradient map can be thought of as a geological map with values representing the height of the land. Thus, low gradient values correspond to “valleys/bowls” and high gradient values correspond to “hills/peaks”. The watershed algorithm (https://en.wikipedia.org/wiki/Watershed_(image_processing)) works by placing “seeds” at the minima of the gradient map. The seeds are then grown (imagine rain flooding the geological landscape) until the boundaries of grown seeds (parcels) meet other grown seeds (parcels). The boundary vertices between parcels are set to 1, while non-boundary vertices are set to 0, thus resulting in a binarized gradient map.

